# Multiplicative couplings facilitate rapid learning and information gating in recurrent neural networks

**DOI:** 10.1101/2025.07.11.663676

**Authors:** Xiaohan Zhang, Mohamad Altrabulsi, Wenqi Xu, Ralf Wimmer, Michael M. Halassa, Zhe Sage Chen

## Abstract

The mammalian forebrain is the seat of higher cognition with architectural parallels to modern machine learning systems. Specifically, the cortex resembles recurrent neural networks (RNNs) while the thalamus resembles feedforward neural networks (FNNs). How such architectural features endow the forebrain with its learning capacity, is unknown. Here we take inspiration from empirical thalamocortical discovery and develop a multiplicative coupling mechanism between RNN-FNN architectures that collectively enhance their computational strengths and learning. The multiplicative interaction imposes a Hebbian-weight amplification onto synaptic-neuronal coupling, enabling context-dependent gating and rapid switching. We demonstrate that multiplicative feedback-driven synaptic plasticity achieves 2-100 folds of speed improvement in supervised, reinforcement and unsupervised learning settings, boosting memory capacity, model robustness and generalization of RNNs. We further demonstrate the efficacy and biological plausibility of multiplicative gating in modeling multiregional circuits, including a prefrontal cortex-mediodorsal thalamus network for context-dependent decision making, a cortico-thalamic-cortical network for working memory and attention, and an entorhinal cortex-hippocampus network for visuospatial navigation and sequence replay. Taken together, our results demonstrate the profound insights into neuroscience-inspired computation that enable multi-plastic attractor dynamics and computation in recurrent neural circuits.

## Main

Recent advances in NeuroAI have taught us good lessons to design brain-inspired AI systems^1–3^ (Hassibi et al., 2017; Zador et al. 2023; Kozachkov et al., 2023). The list of neuroscience-inspired neural network examples includes the convolutional neural network (CNN) motivated by hierarchical visual systems^4^ (LeCun et al., 1998), the capsule neural network (CapsNet) inspired from microcolumns in the cerebral cortex^5^ (Sabour et al., 2017), the deep Q-network (DQN) motivated by experience replay^6^ (Sorokin et al., 2015), and the Transformer motivated by the hippocampal formation^7^ (Whittington et al., 2022).

Mammalian brains contain neural networks with predominantly recurrent and feedforward connectivity. Recurrent connections between excitatory and inhibitory neurons are commonly found in neocortical structures. Recurrent neural networks (RNNs) have been widely used to model the cerebral cortex with strong or weak recurrent excitatory and inhibitory neurons^8–15^ (e.g., Sussilo and Abbott, 2009; Song et al. 2016; Yang et al., 2019; Pollock and Jazayeri, 2020; Zhang et al., 2021; Zhang et al., 2022; Zhang et al., 2025; Lakshminarasimhan et al., 2024). In contrast, some subcortical areas have predominantly feedforward neural network (FNN) structures, with rare or no recurrent connectivity. The reciprocal interactions between these cortical and subcortical structures form a common RNN-FNN loop architecture.

The ability of learning things rapidly with few examples is a desirable property for biological and artificial systems. The brain is flexible and robust, enabling us to perform efficient computations in dynamic and uncertain environments as well as with a small number of labeled examples (‘fewer-shot learning’) or with minimal supervision. However, unsupervised or reinforcement learning (RL) often suffers from learning instability when the sample batch size is small, creating an obstacle for few-shot and sequential learning. Therefore, designing RNNs with *rapid, robust*, and *stable* learning characteristics has been of great importance. Motivated by the biological RNN-FNN loop structure that enables rapid learning in animals and humans, here we hypothesize that multiplicative interactions^16^ (Jayakumar et al., 2020) is the key ingredient for flexible and robust learning and propose a multiplicative gating-based RNN-FNN architecture that resembles the thalamocortical architecture in the brain. Two key features are noteworthy. First, this framework accommodates the interaction between a fast-weight readout memory (‘FNN’) and a slow-weight network (‘RNN’), where the slow weights are dynamically modulated by the fast-weight network output in a multiplicative manner. The use of fast weights for modeling sequences and improving neural network performance has been reported in several lines of work^17–20^ (Hinton and Plaut, 1987; Schmidhuber, 1992; Ba et al., 2016; Irie et al., 2021). Second, multiplicative synaptic-neuronal interactions can be mathematically represented as a Hadamard product between a recurrent connectivity matrix and a lower-rank outer-product matrix, allowing us to impose hardwired or context-dependent constraints onto RNN connectivity and implement a Hebbian-weight amplification or suppression in synaptic plasticity. While our proposed idea is relatively simple, it not only offers biological support and flexibility for modeling multi-area neural circuits with bidirectional interactions but also yields profound benefits in machine learning applications.

Through extensive computer experiments and benchmark comparisons, we show that the proposed dynamic, multiplicative synaptic-activity couplings significantly improve the learning speed, flexibility, robustness, and generalization in both rate-based and spiking-based RNN models for a wide range of tasks, including cognition, working memory (WM), attention, spatial navigation, pattern recognition, sentiment analysis, and robotic control. We demonstrate the computational advantages of ‘dynamic synapses’ used in RNNs across various machine learning paradigms, ranging from supervised learning based on back-propagation-through time (BPTT) algorithm^10,21,22^ (e.g., Song et al., 2016; Bellec et al., 2020; Xue et al., 2023), biologically inspired unsupervised spiking-timing-dependent plasticity (STDP) algorithm^23,24^(Fernandez et al., 2024; Dong et al., 2023), and RL based on policy gradient optimization^25^ (Song et al., 2017). Furthermore, we demonstrate the strengths of multiplicative gating in modeling brain circuits with the loop, chain or hierarchy structure between recurrent and feedforward modules. Neuroscience examples include a prefrontal cortex-mediodorsal thalamus (PFC-MD) network in context-dependent decision making^14^ (Zhang et al., 2025), a cortico-thalamic-cortical structure that mimics PPC-pulvinar-PFC network in performing WM and attention, and a hierarchical entorhinal cortex-hippocampus (EC-DG-CA3-CA1) network during visuospatial navigation and sequence replay. Notably, our proposed modeling paradigm for three different neural circuits provides numerous predictions that well match experimental data in animal studies, supporting the biological plausibility of multiplicative gating. Finally, we discuss the broad implications of multiplicative gating mechanisms in neuroscience and AI.

## RESULTS

### From vanilla RNN to multiplicative RNN-FNN coupling

Without the loss of generality, we start with a rate-based vanilla RNN and express its neural dynamics 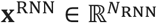 via a continuous-time differential equation:

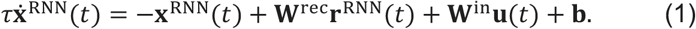

where *τ* denotes the time constant, and dynamics evolve at a time incremental of d*t*. In Eq. (1), **W**^rec^ denotes an *N*_RNN_-by-*N*_RNN_ recurrent connection weight matrix of *N*_RNN_ hidden RNN units that describes the state transition and may incorporate biological connectivity (e.g., Dale’s principle or cell-type-specific connectivity), and **W**^in^ denotes a connection weight matrix between an *m*-dimensional external input **u** to the *N*_RNN_-dimensional hidden recurrent units; **b** denotes a bias term, and **r**^RNN^ = *ϕ*(**x**^RNN^) denotes the transformed nonnegative firing rate activity through an elementwise nonlinear activation function *ϕ*. In neuroscience or engineering applications, various choices of the activation function include the ReLu, softplus, logistic sigmoid, and hyperbolic tangent functions.

#### Additive gating with FNN feedback

Next, we consider an RNN-FNN network with reciprocal connectivity and bidirectional interactions (**Fig. 1a**). A traditional view of the FNN’s role is to gate the RNN activity as another additive external input. Mathematically, the following two equations describe the additive dynamics of rate-based RNN and FNN, respectively

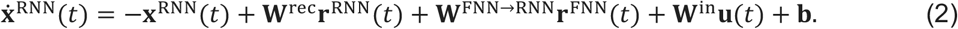

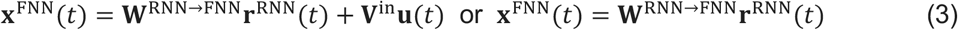

where **b** denotes a bias term in a general setting (sometimes assumed to be zero); **W**^FNN→RNN^ and **W**^RNN→FNN^ denote the respective *N*_FNN_-by-*N*_RNN_ (usually *N*_FNN_ ≤ *N*_RNN_) and *N*_RNN_-by-*N*_FNN_ weight matrices between FNN and RNN units, **V**^in^denotes the connection weight of the input to recurrent units (in some special cases, **V**^in^ = **0** when the FNN receives no direct external input); and **r**^FNN^ = *ϕ*(**x**^FNN^). To simplify the mathematical notation, we assume **V**^in^ = **0** and let

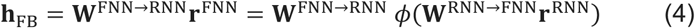

denote an efference copy of the random or learnable projections of RNN feedback activity; the RNN dynamics described in Eq. (2) can be rewritten as

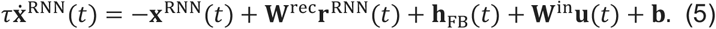

**Figure 1.**
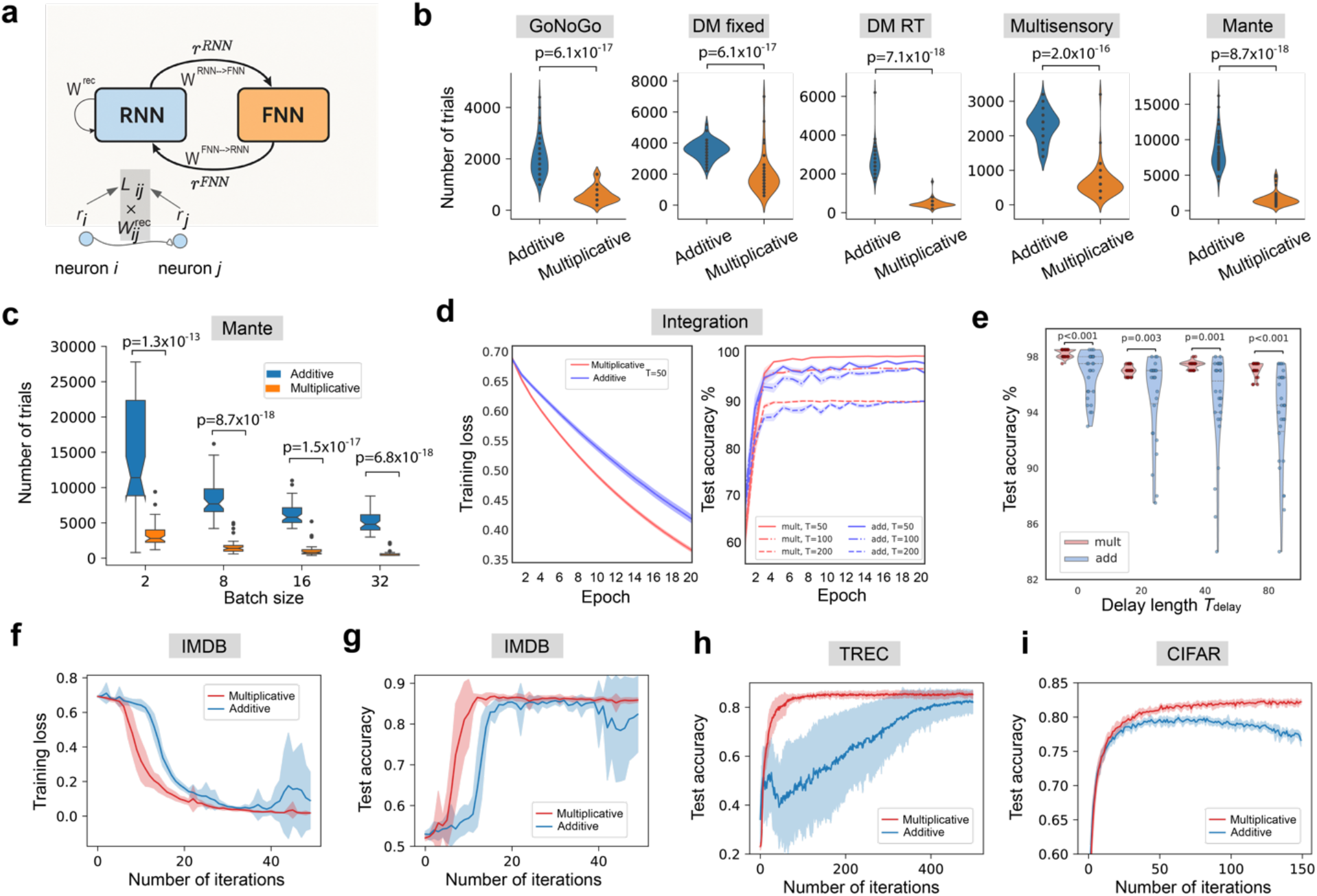
Multiplicative RNN-FNN couplings facilitate rapid learning in supervised learning benchmarks. **(a)** Schematic of bidirectional RNN-FNN loops and multiplicative synaptic-neuronal gating modeled as 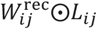. **(b)** Convergence speed comparisons on five WM and DM benchmarks based on supervised learning (*N*_RNN_=128, *N*_FNN_ = 128, batch size 8). Monte Carlo experiments (n=50 random seeds, p-values were derived from the rank-sum test). **(c)** Convergence speed comparison on the Mante task benchmark with respect to four different numbers of batch size (p-values were derived from the rank-sum test, n=50 for each condition). **(d)** Comparison of convergence curves of training loss (left, sequence length *T* = 50) and testing classification accuracy (right, sequence length *T*=50,100,200) on the integration problem. Shaded area denotes s.e.m. (n=30). **(e)** Violin plot comparison of generalization in testing classification accuracy (sequence length *T* = 50, plus varying delay length *T*_delay_) in the delayed integration task. P-values were derived from paired t-test (n=30) with FDR correction. **(f)** Comparison of training loss curves in the IMDB classification benchmark between self-gated and vanilla RNNs. The horizontal axis denotes the number of learning iterations (number of epochs × training sample size / batch size). Batch size: 256. Shaded area denotes s.e.m. (n= 10 random seeds). **(g)** Same as **f**, except for test accuracy curves. **(h)** Comparison of test accuracy curves in the TREC classification benchmark between self-gated and vanilla RNNs. The horizontal axis denotes the number of learning iterations (number of epochs × training sample size / batch size). Shaded area denotes s.e.m. (n= 10 random seeds). **(i)** Same as **h** but for the CIFAR classification benchmark. Shaded area denotes s.e.m. (n= 10 random seeds).

We refer this as an additive RNN-FNN coupling in that the feedback from the FNN output **h**_FB_(*t*) strengthens or suppresses the RNN activity **x**^RNN^ through additive gating.

#### Multiplicative gating with FNN feedback

In contrast to Eq. (5), a multiplicative RNN-FNN coupling allows the FNN feedback activity to directly modulate **W**^rec^:

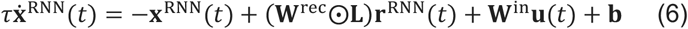

where ⨀ denotes the matrix Hadamard product between two matrices **W**^rec^ and **L** = **h**_FB_**h**_FB_^*T*^ of the same size, and **L** is a low-rank symmetric matrix (i.e., rank(**L**)< *N*_RNN_). In general, the mask **L** is a modulatory matrix that may represent structural or cell-type connectivity constraints^10,14^ (Song et al., 2017; Zhang et al., 2025), context-dependent gating, or learned modulation (**Supplementary Note 1**). Multiplicative interactions allow fine-grained control over which dynamic recurrent connections are strengthened in time such that the FNN activity **r**^FNN^ can rapidly change the hidden RNN state.

For notation simplicity, let 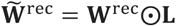 denote the effective masked recurrent weight matrix; from linear algebra rearrangement, 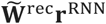 can be rewritten as *diag*(**W**^rec^**D**_r_**L**^*T*^) = *diag*(**W**^rec^(**D**_r_**L**^*T*^)) = *diag*(**W**^rec^(**D**_r_**L**)) (**Supplementary Note 2**), where **D**_r_ = *diag*(**r**^RNN^) denotes a diagonal matrix with the diagonal being a vector **r**^RNN^. If **h**_FB_ is a non-zero vector, **L** will be a rank-1 matrix, and the rank of 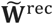 will be lower-bounded by **W**^rec^. Since matrix multiplication is associative, we can rewrite **Lr**^RNN^(*t*) = **h**_FB_(**h**_FB_^*T*^**r**^RNN^(*t*)); the inner product between the feedback **h**_FB_ and the RNN activity **r**^RNN^(*t*) resembles multiplicative attention. It is noteworthy that the two RNN models described in Eq. (5) and Eq. (6) have different dynamics despite sharing the same degrees of freedom in the model parameter space.

In a special case where the bidirectional RNN-FNN weights {**W**^FNN→RNN^, **W**^RNN→FNN^} are excitatory and ***ϕ*** is a ReLu function, the outer product **h**_FB_**h**_FB_^*T*^will produce a scaled copy of autocorrelation of **r**^RNN^. To see it clearly, we rewrite Eq. (6) in a scalar form:

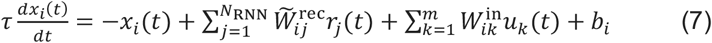

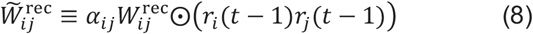

where *α*_*ij*_ ≥ 0 denotes a nonnegative scaling coefficient. Equation (8) implies a Hebbian-weight amplification mechanism in that the synaptic strength between neurons *i* and *j* is multiplicatively gated by a nonnegative Hebbian term *r*_*i*_*r*_*j*_----a product termed ‘multiplicative units.’ The magnitude of 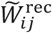 is locally amplified by the presynaptic and postsynaptic activity together, and the sign of excitatory (positive) or inhibitory (negative) weight 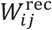 preserves Dale’s principle (Note: if the RNN-FNN weights are not purely excitatory, we can also impose a monotonic logistic sigmoid function to the feedback term assure the gating coefficients are non-negative). The gradient optimization of RNN weights in Eq. (7) can be written in a chain rule: 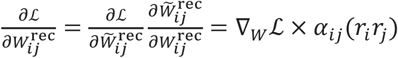, where *α*_*ij*_ (*r*_*i*_ *r*_*j*_) denotes a dynamic activity-dependent gating factor modulating the vanilla gradient ∇_*W*_ℒ computed from the loss function ℒ. This implies that local Hebbian masking controls the flow of information, allowing the RNN to dynamically enhance or suppress synaptic strengths based on correlated pre- and post-synaptic activities.

#### Multiplicative gating with time-delay feedback

When **W**^RNN→FNN^ and **W**^FNN→RNN^ are both identity matrices, the RNN-FNN loop will be equivalent to a self-gated RNN with only time-delay feedback. Sharing similar concepts of embedded Hopfield auto-associative memory^19^ (Ba et al., 2016) and multi-plasticity network (MPN)^26^ (Aitken and Mihalas, 2023), the self-gated RNN dynamics is described by

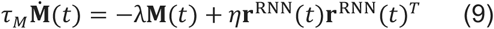

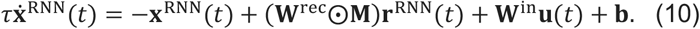

where *τ*_*M*_ denotes a time constant for the fast-weight dynamics of time-embedding associative memory of RNN activity; the mask matrix **M**, as a form of Hebbian synapses (*M*_*ij*_ ≥ 0), computes a moving average of the autocorrelation of **r**^RNN^. In Eq. (10), the recurrent weights between presynaptic and postsynaptic neurons are multiplicatively gated and self-stabilized by their associative memory traces (**Supplementary Note 3**).

In RNN-FNN models with either additive or multiplicative couplings, the update of recurrent weights **W**^rec^ and non-recurrent weights {**W**^in^, **W**^FNN→RNN^, **W**^RNN→FNN^} depends on the synaptic plasticity rule. In the subsequent sections, we will demonstrate that multiplicative gating in the RNN-FNN architecture outperforms the additive gating counterpart for a wide range of learning paradigms and for both rate-based and spiking neural networks.

### Multiplicative couplings facilitate rapid learning

First, we trained RNN-FNN models, with either additive (Eq. 5) or multiplicative couplings (Eq. 6), to perform five representative benchmarks of WM and decision-making (DM) tasks^11^ (Yang et al., 2019): Go/NoGo, DM mixed, DM_RT, Multisensory, and Mante^9^ (**Methods** and see **Supplementary Table 1** for details). We compared their convergence speed by running 20 different random seeds and standard BPTT algorithm with identical network size (*N*_RNN_=128 and *N*_FNN_=128), weight initialization, convergence and assessment criterion and default hyperparameters (e.g., learning rate, regularization parameter) (**Supplementary Table 2**). No extra effort was applied for hyperparameter optimization. Among all benchmarks, we found that multiplicative couplings improved the convergence speed of the additive coupling model by ~2-10 folds based on a fixed batch size of 8 (**Fig. 1b**). While decreasing the batch size generally slowed down the learning speed, multiplicative couplings facilitated rapid learning even in the presence of a small batch size (**Fig. 1c** and **Extended Data Fig. 1**).

To gain an intuition of the reason why multiplicative gating achieves faster convergence speed, we employed the multiplicative RNN-FNN to perform a simple evidence integration task^26^ (Aitken and Mihalas, 2023). We employed a minimal network size (*N*_RNN_=3 and *N*_FNN_=3) that was nevertheless sufficient to learn this simple task (training sequence length *T* = 50, time constant *τ* = 2, *dt* = 0.1). Notably, the multiplicative coupling model was able to learn the task rapidly with a small batch size of 4 (left panel, **Fig. 1d**). Next, to test the model generalization, we used various configurations of input length (*T* = 50, 100, 200) and found that multiplicative coupling outperformed additive coupling with better testing accuracy in both speed (4 epochs vs. 20 epochs) and steady-state performance (right panel, **Fig. 1d**). Furthermore, to test WM in the context of integration, we inserted a continuous delay period *T*_delay_ (during which the input was absent or noise interference was present) between the first and second halves of input sequence (so that the total length is *T* = 25 + *T*_delay_ + 25) and found that multiplicative coupling yielded consistently better test accuracy (i.e., higher mean and lower variance) for a wide range of delay durations (*T*_delay_= 0, 20, 40, 80; **Fig. 1e**). Given initial conditions, dimensionality reduction analysis (**Methods**) revealed that multiplicative gating and additive gating networks formed distinct low-dimensional clusters in their 3-by-*T* dimensional neural trajectories (**Extended Data Fig. 2a**). Theoretical analysis further revealed that multiplicative coupling models maintained low variance in the internal update compared to the additive coupling models (**Supplementary Note 4**), matching the results of computer simulations and offering an interpretation of the accuracy discrepancy between them (**Extended Data Fig. 2b-d**).

Next, we compared the performance between the multiplicatively self-gated RNN (Eq. 10) and the vanilla RNN (Eq. 1) on a sentiment analysis benchmark using the IMDB movie review dataset, in which the high-dimensional text data were preprocessed and subsequently fed into the RNN (**Methods**). We computed the classification accuracy on the 20% held-out testing data. For an identical parameter setup, the multiplicatively gated RNN achieved faster convergence than the vanilla RNN (**Fig. 1f**) and reached ~87.5% mean accuracy plateau (**Fig. 1g**). By monitoring the gradient magnitude of the loss function with respect to the recurrent weights 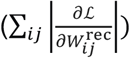 during RNN training, we found that the multiplicative RNN’s gradient rose rapidly at the beginning of training and decayed faster towards convergence (**Extended Data Fig. 3a**).

Finally, we conducted experiments on two machine learning benchmarks: Text Retrieval Conference (TREC) and CIFAR-10 tasks and observed consistent better results in multiplicative gating (**Fig. 1h,i** and **Extended Data Fig. 3b,c**). Put together, learning was more stable with multiplicative coupling, whereas the vanilla RNN could run into the risk of overfitting, resulting in a decrease in test accuracy.

### Multiplicative couplings enable robust and persistent dynamics

To gain insights into the efficacy of the multiplicative coupling mechanism, we used the “DM fixed” (with WM delay) benchmark to test the role of feedforward weights {**W**^FNN→RNN^, **W**^RNN→FNN^} with the following experiments. First, we randomized the feedforward weights and froze them during training. Additionally, we randomly set a subset of fixed bidirectional feedforward weights to zero (sparsity level: 0.2, 0.4, 0.6, 0.8). The sparsity level defines the percentage of zeros among all feedforward connections. Surprisingly, these sparse, random non-recurrent connections had little impact on the convergence speed of RNN (**Fig. 2a**). Notably, the median convergence speed was relatively stable regardless of the selection of unit connectivity and the sparsity level, suggesting that the plasticity of recurrent weights **W**^rec^ plays the key role in determining the RNN equilibrium state, whereas random and sparse readouts of the recurrent units and/or random feedback projections are sufficient to produce context-dependent gating (**Fig. 2b**).

**Figure 2.**
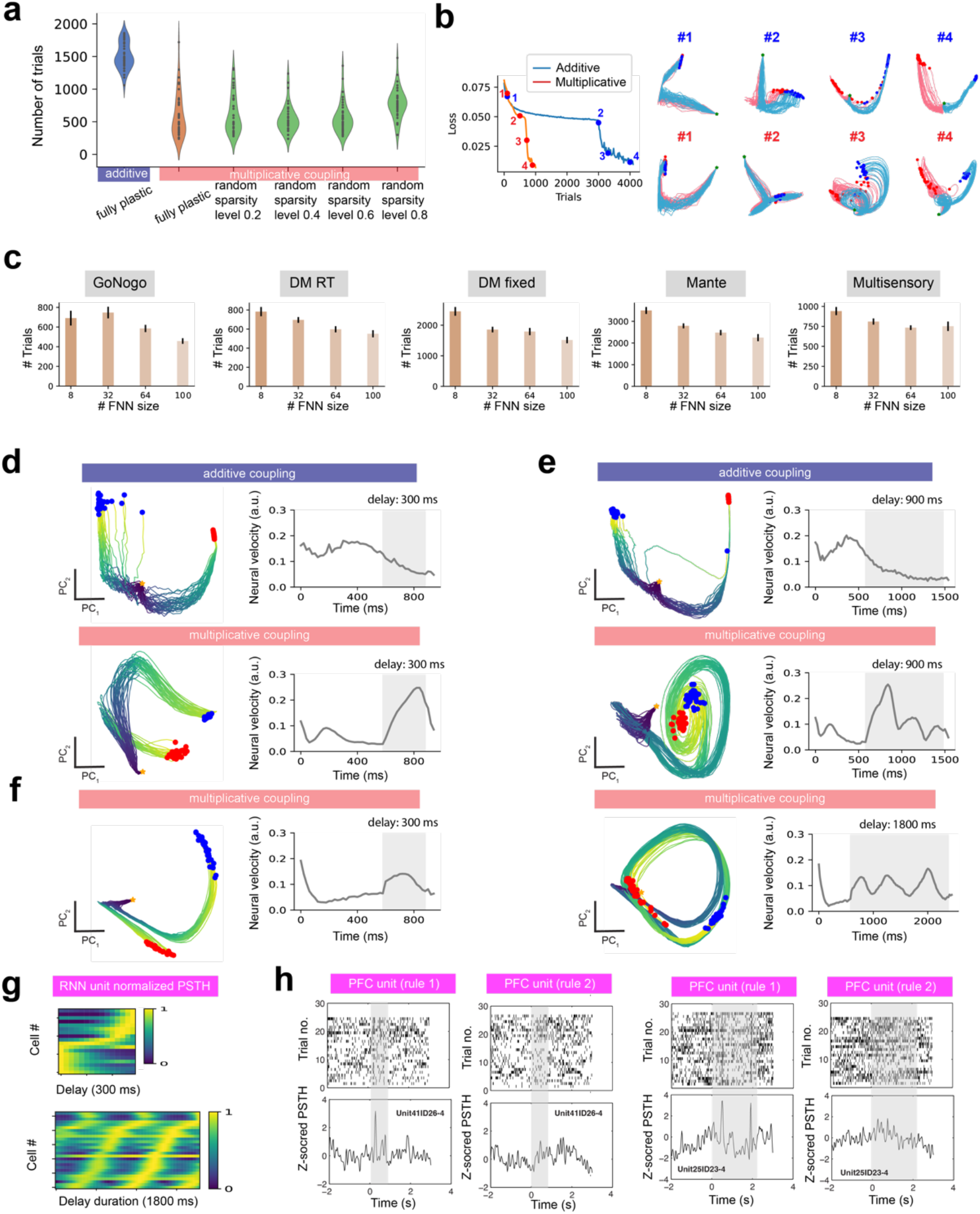
Multiplicative RNN-FNN couplings enable robust and persistent dynamics. **(a)** Sparse, random non-recurrent RNN-FNN connections had little impact on the learning speed. **(b)** Representative training convergence curves. Evolution of neural trajectories in principal component subspace (PC_1_ and PC_2_) throughout learning. Each line represents a trial, with trial time color coded. Star symbol * denotes the trial onset, and circles denote the trial offset, with blue/red color representing different outcomes. **(c)** Impact of the FNN size on the convergence speed of five cognitive task benchmarks in the multiplicative gating setting. Error bar denotes s.e.m. (n=50 independent models). **(d)** Upon convergence, comparisons of neural trajectories and neural velocity between additive and multiplicative couplings. Neural velocity analysis showed that the dynamics of RNN-FNN with additive coupling (top panel) decayed quickly to a fixed point during the WM delay period (shaded area), but the dynamics with multiplicative coupling (bottom panel) generated ‘bump’ activity. Each line represents a trial. Blue and red dots represent the end points of trials from two distinct classes. Star symbol represents the trial onset. Duration of delay period: 300 ms. **(e)** During extended delay (900 ms), the additive coupling model maintained the fixed point, whereas the multiplicative coupling model showed oscillatory dynamics. **(f)** In some cases, multiplicative couplings generated limit-cycle like attractor dynamics during extended delay (1800 ms). **(g)** The RNN-FNN with multiplicative coupling generated periodic sequence representations from the RNN population activity during 1800-ms delay period. Each heatmap shows the normalized peri-stimulus time histogram (PSTH) of a subset of RNN units. **(h)** Spike rasters of two representative mouse PFC neurons and their z-scored PSTHs for encoding two conflicting rules in a cross-modal decision-making task. During shorter delay (shaded area [0, 0.9] s, left), one single peak was observed, while two peaks emerged during extended delay (shaded area [0 2.1] s, right), resembling quasi-periodicity encoding.

To investigate the impact of FNN on learning speed in the multiplicative coupling setting, we systematically varied the size of FNN and repeated five WM and DM benchmark experiments using a fixed minibatch size of 8 and sparsity level of 0.2. We observed a general trend in slower convergence with a decreasing number of FNN units (**Fig. 2c**), but the results were relatively stable. These results suggest that lower-dimensional FNN (*N*_FNN_<*N*_RNN_) can read out random projections of context-dependent information for subsequent RNN connectivity gating, which further stabilize the attractor dynamics in learning.

We hypothesized that the feedback activity **h**_FB_ provided task-relevant gating information which could lead to context-dependent RNN attractor dynamics. While the objective is the same, the solution may be different between additive and multiplicative gating strategies. An illustration of a two-dimensional RNN-FNN system supported our intuition that multiplicative gating based on different feedback activity may introduce dynamic phase reconfiguration (see **Supplementary Note 5** and **Extended Data Fig. 4** for details) and further facilitate flexible cognitive control. Furthermore, in the actual “DM fixed” benchmark, we examined the multiplicative gating matrix **L** = **h**_FB_**h**_FB_^*T*^ computed from the FNN feedback and found that task-dependent structures gradually emerged from low-dimensional neural embeddings in the PCA subspace (**Fig. 2d**). We noticed that multiplicative gating produced a smoother trajectory evolution during the learning process (**Supplementary Video 1**). After the completion of learning, multiplicative gating enabled the RNN dynamics to maintain persistent activity during WM, as evidenced from the trajectory of neural velocity 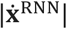 with a sharp increase at the start of WM; in contrast, the neural velocity in additive RNN-FNN coupling decayed gradually during WM (**Fig. 2d**, top panel). Interestingly, when the original WM delay period was extended by 3-6 folds (900-1800 ms), the RNN-FNN with multiplicative coupling sometimes generated damping oscillators (**Fig. 2e**) or limit cycle-like attractors (**Fig. 2f**) during the extended delay period that enabled correct classification, whereas the attractor was unstable in additive RNN-FNN coupling and frequently led to misclassification trials (**Fig. 2e**, top panel).

To systematically test the model robustness with extended delay, we modified the DM task setting (i.e., training with maximum stimulus coherence instead of a wide range of coherence statistics). To test generalization, we gradually varied the WM delay period (300, 400, 500 and 900 ms) and assessed the task performance of two model classes (multiplicative vs. additive, n=50 for each class). We found that the multiplicative models obtained higher mean accuracy in task performance (**Extended Data Fig. 5**), suggesting that WM was maintained more robustly based on oscillatory dynamics.

Of note, the presence of oscillatory or limit-cycle attractor during the extended delay period implies the quasi-periodicity of neuronal firing during WM. This phenomenon was not only evidenced in our computer experiments (**Fig. 2g**) but also experimentally observed in single-unit recordings of task-performing mouse PFC in a context-dependent cross-modal decision-making task^27^ (Rikhye et al., 2018; **Methods; Fig. 2h** and see **Extended Data Fig. 6** for more examples). In this case, sequential population activity was reflected by the oscillatory dynamics in neural subspace^28^ (Lebedev et al., 2019).

### Multiplicative couplings accelerate reinforcement learning

One conceptual limitation of supervised training for RNNs in cognitive tasks is that each trial is treated independently, and the current trial outcome (error or success) does not affect the next trial. To overcome this limitation, we adopted an actor-critic RL approach to train the RNN-FNN model (**Fig. 3a**), where the network parameters were updated based on recurrent policy gradient (**Methods**). We first tested two RNN-FNN models on the five WM and DM benchmarks and compared the relative RL convergence speed between additive and multiplicative couplings. One commonly observed problem in RL is learning instability given a small batch size. In our experimental setup (batch size: 16), we found that multiplicative couplings improved the stability and convergence by ~5-8 folds (**Fig. 3b**).

**Figure 3.**
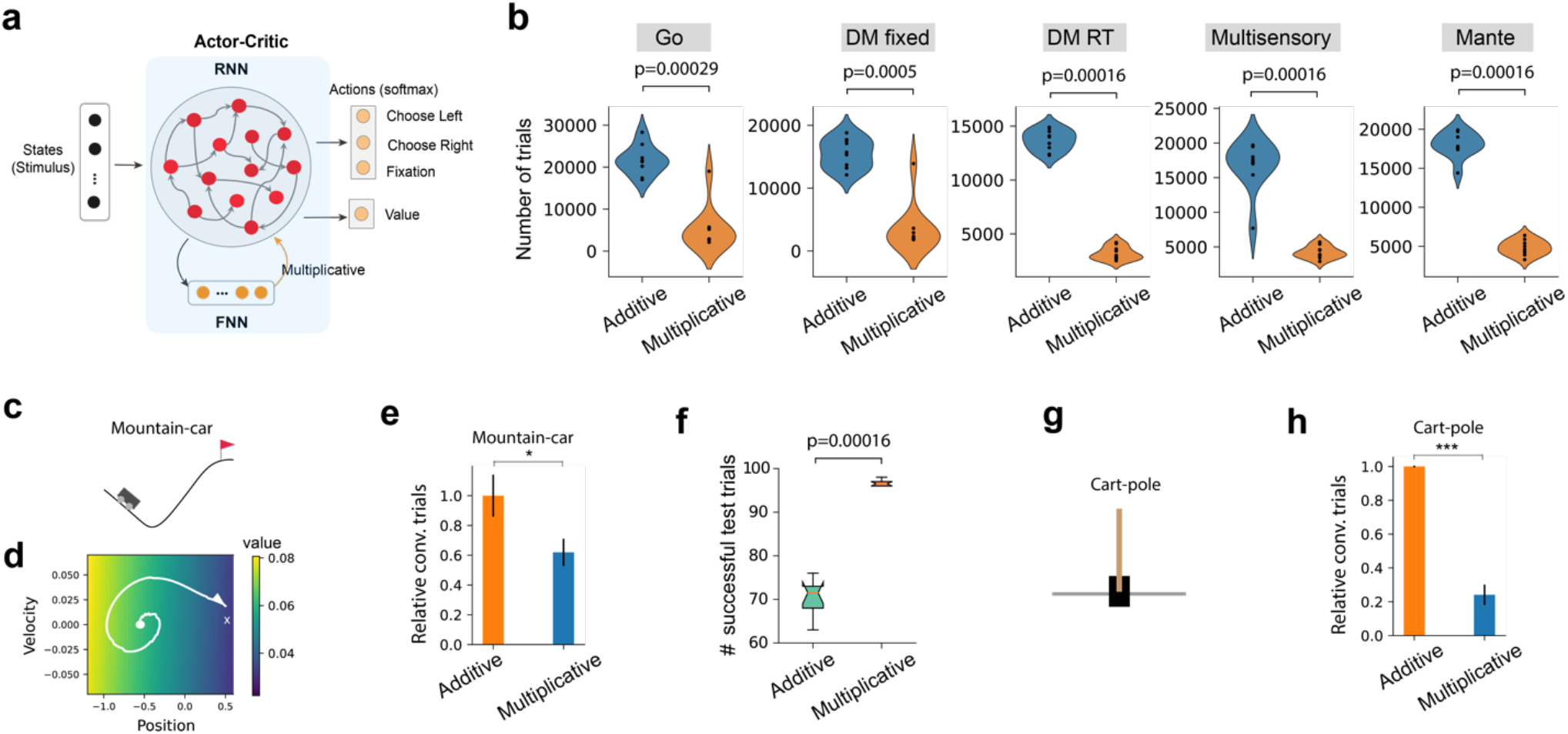
Multiplicative RNN-FNN couplings improves the RL speed. (a) Schematic of actor-critic RL applied to RNN-FNN (*N*_RNN_=128, *N*_FNN_=64) training for cognition and control tasks. The RNN was trained to produce a policy (the strategy that an agent chooses action given a state based on softmax) and a value function estimate. (b) Comparison of RL convergence speed on the five WM and DM benchmarks (batch size: 16) shown in Fig. 1b (all p-values were calculated from the rank-sum test, n=10). (c) Schematic of the mountain-car problem. (d) The inferred approximate value function in the position-velocity space. The white trajectory represents a successful trial from the initial condition (‘dot’) to the target condition (‘cross’). (e) Relative convergence speed comparison on the mountain-car benchmark (*, p=0.023, rank-sum test, n=15; (*N*_RNN_=64, *N*_FNN_=32). (f) Comparison on the number of successful trajectories of the trained RL agents among 100 test trials of the mountain-car problem based on initial conditions (p=0.00016, rank-sum test, n=100). (g) Schematic of the cart-pole problem used in the actor-critic RL (*N*_RNN_=64, *N*_FNN_=32). (h) Relative convergence speed comparison on the cart-pole benchmark (***, p=9.7×10^−9^, rank-sum test, n=30).

We further tested the RNN-FNN models on two classic continuous control and RL benchmarks: the mountain-car and cart-pole problems (**Methods**). The objective of controlling mountain-car (**Fig. 3c**) was to learn a state-action (***s, a***) policy or the approximation of a value function *V*(***s, a***), where the continuous state set ***s*** consists of the position and velocity of the car, and the discrete action set ***a*** consists of three options: ‘accelerate forward’, ‘accelerate backward’, and ‘do nothing’. Upon RL convergence, we inferred the 2D state value function (**Fig. 3d**). Our results showed that multiplicative coupling improved the RL convergence speed by ~40% (**Fig. 3e**; p=0.023, rank-sum test), especially the results held for a small minibatch size. Furthermore, we also run additional 100 test trials to assess the model generalization. We found that multiplicative coupling produced a greater number of successful trajectories (criterion: episodes with return > −200) than the additive coupling (**Fig. 3f**; p=0.00016, rank-sum test), suggesting that multiplicative couplings boost learning stability and control, leading to more consistent goal-reaching behavior. Similar results were found in the cart-pole problem (**Fig. 3g**), whose objective was to apply a force to the cart and prevent the pole falling over. In this case, multiplicative coupling achieved rapid convergence with ~75% improvement over additive coupling (**Fig. 3h**; p=9.7×10^−9^, rank-sum test). Taken together, these benchmark experiments support that multiplicative couplings in RNN-FNN models improve learning speed and stability in RL applications.

### Multiplicative gating in spiking RNNs improves biologically realistic learning speed

Spiking neurons are an important hallmark of the biological brain in stochastic computation. Next, we extended the analysis to the spiking RNN (SRNN) composed of spiking units. We developed a SRNN model with multiplicative self-coupling as well as its non-coupled counterpart (**Methods**). We trained these two classes of SRNN models based on both unsupervised, biologically plausible STDP and supervised BPTT learning rules, and then compared their performances on a simulated line detection task and a real-world MNIST benchmark.

With the same hyperparameter setup (**Supplementary Table 3**), the convergence of SRNN was influenced by the batch size and network size. In the line detection task (**Fig. 4a**), we varied these two parameters and found consistent improvement of convergence speed in the multiplicatively gated SRNN across all conditions (**Fig. 4b**), based on the same convergence assessment and initial weight conditions. Generally, relative speedup (~10-50%) was more pronounced for a larger network size. In STDP, we systematically varied two hyperparameters (*λ, η*) in Eq. (8) and repeated each setup 100 Monte Carlo runs with batch size of 1. In the case of multiplicative couplings, consistent improvement was found in both convergence speed and test accuracy; results were robust for various choices of hyperparameters (**Fig. 4c**). Similar observations were found in BPTT (**Fig. 4d** and **Extended Data Fig. 7** for various batch sizes). Note that multiplicative couplings achieved low variance in all results (especially in BPTT), whereas a larger network size without multiplicative coupling did not always yield a good test accuracy due to overfitting.

**Figure 4.**
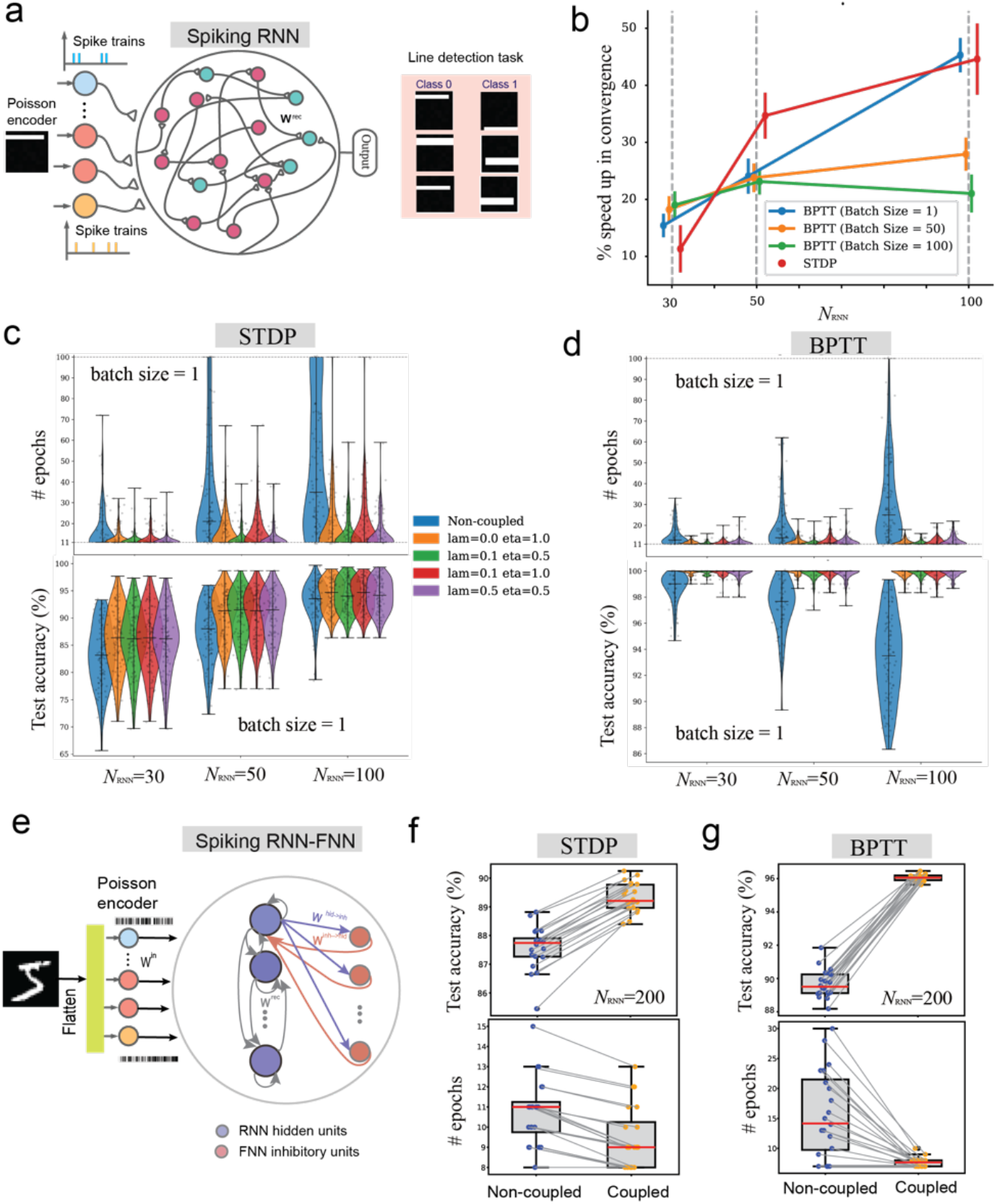
Spiking RNNs (SRNNs) applied to the line detection and MNIST benchmarks. (a) Schematic of the SRNN (left) for the line detection problem (right). (b) Relative convergence speedup of multiplicative couplings versus non-coupling with respect to the batch size and network size. Error bar denotes s.e.m. (n=100 random seeds). (c) Violin plot comparisons of STDP convergence speed and accuracy between SRNNs with respect to the hyperparameters in associative memory (*λ, η*) and the network size *N*_RNN_. Results were robust for a wide range of hyperparameter setups. Horizontal lines indicate median statistics of the distributions. (d) Same as **c** but for BPTT. (e) Schematic of the spiking RNN-FNN applied to MNIST handwritten digit classification. The 28×28 images were first flattened and then passed onto a Poisson encoder to produce 784-dimenisonal parallel spike trains. The input spike trains were sent to *N*_RNN_ =200 hidden units via an input weight matrix **W**^in^. The excitatory units were fully connected via a plastic recurrent weight matrix **W**^rec^. Additionally, the excitatory units received input from *N*_FNN_ =50 inhibitory units through fixed inhibitory synapses **W**^inh→hid^ and projected the reciprocal output to inhibitory units through fixed excitatory synapses **W**^hid→inh^. (f) Box plot comparisons of STDP test accuracy (p<10^−10^, Wilcoxon signed-rank test, n=20) and convergence speed (p = 0.0172, Wilcoxon signed-rank test, n =20 random seeds). Batch size: 1. (g) Same as **g** but for BPTT (p<10^−10^ for accuracy comparison and p=0.0005 for convergence comparison., Wilcoxon signed-rank test, n = 20). Batch size: 256.

We further applied the SRNN to solve a MNIST handwritten digit classification benchmark. As a control model (without coupling) for comparison, we modified a previously proposed spiking neural network^29^ (Diehl and Cook, 2018) with additional inhibition mechanism. The network consisted of one SRNN containing 200 hidden units and an additional spiking FNN containing 50 inhibitory-only neurons (i.e., no lateral inhibition) (**Fig. 4e**). Therefore, it can be viewed as a special spiking RNN-FNN (**Methods**). In both STDP and BPTT settings, we reported consistently better testing accuracy and training convergence speed with multiplicative self-coupled gating (**Fig. 4g**), supporting that multiplicative coupling in SRNNs may achieve good learning generalization for challenging problems.

### Modeling cognitive flexibility in a biologically constrained thalamocortical network

One important hallmark of cognitive flexibility is the capability of performing neural computation rapidly to accommodate dynamic, context-switching environments. We adopted the RNN-FNN modeling framework to an excitatory-inhibitory (E/I) PFC network that has reciprocal connections with feedforward excitatory-only MD thalamus (**Methods**). Additive coupling within the E/I-PFC-MD network in a prior study has shown improved WM for rule maintenance and insensitivity to noise interference in cognitive tasks^14^ (Zhang et al., 2025).

To extend the investigation of multiplicative gating in the PFC-MD network, we first trained the E/I-PFC-MD model to perform a sequential context-switching task based on the BPTT algorithm (**Fig. 5a**), where the objective is to learn a cue-to-rule mapping uncertainty in a context-dependent DM manner. We trained the model to learn *Context 1* in a batch manner. Subsequently in context switches (*Context 1 to Context 2* and subsequent *Context 2 to Context 1’*), we adapted the model parameters on a trial-by-trial basis. Notably, the multiplicative PFC-MD coupling facilitated rapid learning during context switching (**Fig. 5b**).

**Figure 5.**
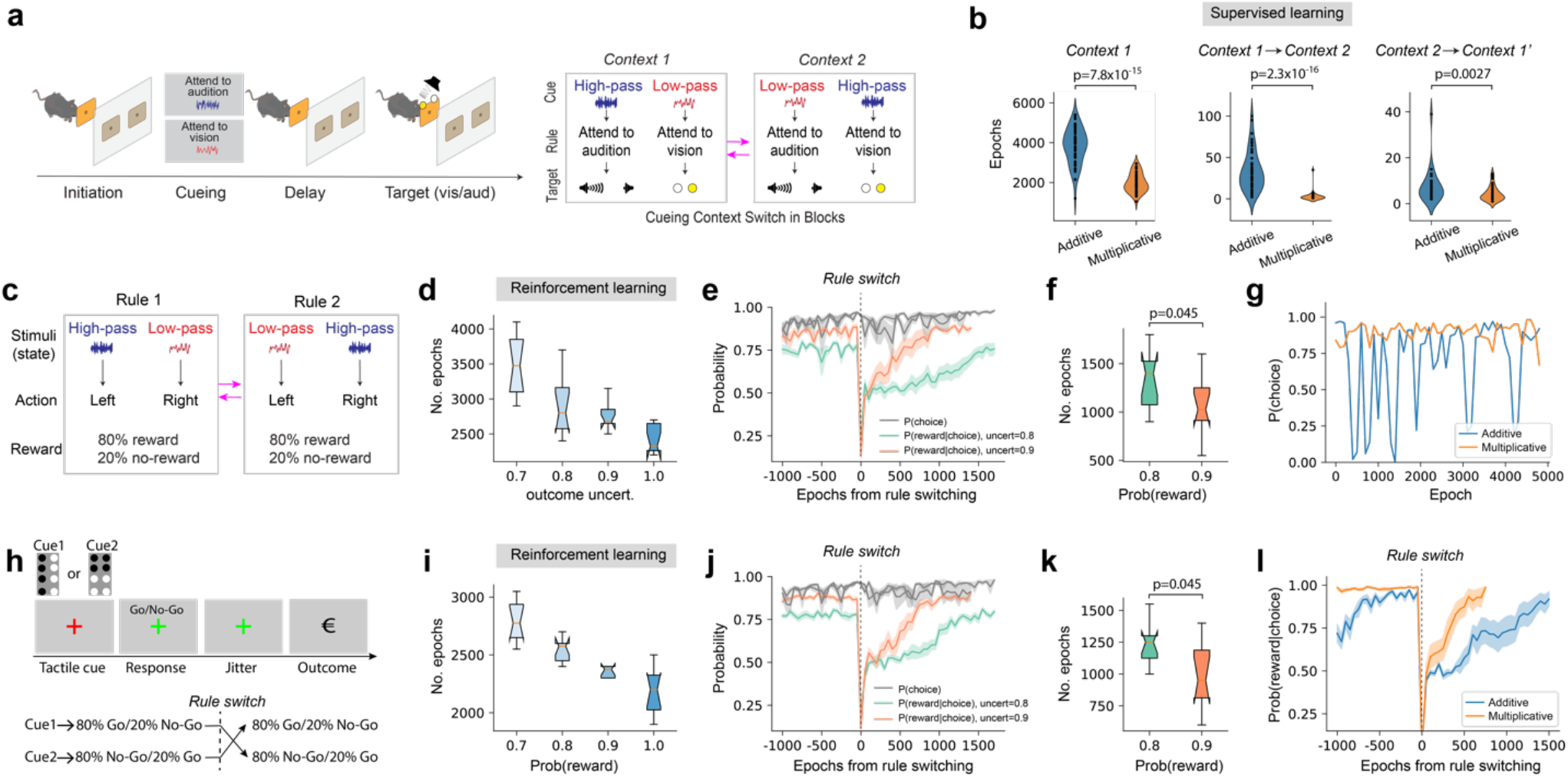
Excitatory-inhibitory (E/I) PFC-MD model for two cognitive flexibility tasks. **(a)** Schematic of the context-switching 2AFC task. **(b)** Multiplicative PFC-MD coupling substantially improved the convergence speed and enabled few-shot learning during context-switching in supervised learning (*N*_RNN_=128, *N*_FNN_=64; batch size: 16, n = 50 random seeds). **(c)** Schematic of context-switching with task outcome uncertainty. **(d)** Outcome uncertainty negatively impacted the RL convergence speed for rule 1. Results are shown for multiplicative coupling as the additive coupling setup failed to converge in RL (*N*_RNN_=128, *N*_FNN_=64; batch size: 16). Box plot shows median and 25%-75% percentiles (n=10). **(e)** Probability of *P*(*choice*) and *P*(*reward*|*choice*) during RL for multiplicative coupling. Two reward uncertainty levels (80% and 90%) are shown: the RL speed for rule switch was faster for 90% reward probability, but the *P*(*choice*) curves for two uncertainty levels were similar. Time 0 indicates the timing of rule-switch. Shaded area indicates s.e.m. (n=10). **(f)** Outcome uncertainty negatively impacted rule switch latency. Box plot shows median and 25%-75% percentiles (n=10). **(g)** Multiplicative coupling produced stable decision probability *P*(*choice*) during RL, whereas the learning curve in additive coupling was unstable (batch size: 2). **(h)** Schematic of the probabilistic Go/NoGo task used for RL (*N*_RNN_=128, *N*_FNN_=64; batch size: 16). (**i-k**) Same as **d-f**, except for the probabilistic Go/NoGo task. **(l)** Comparison of probability *P*(*reward*|*choice*) curves between additive and multiplicative couplings in the probabilistic Go/NoGo task without reward uncertainty. Shaded area indicates s.e.m. (n=10).

We next generalized the task setting to accommodate reward uncertainty such that reward was only given with 80% probability when a correct decision was made (**Fig. 5c**). Because supervised learning treated individual trials independently and could not track inter-trial feedback, we trained the E/I-PFC-MD model based on a policy gradient algorithm known as actor-critic RL (**Methods**) and found that multiplicative couplings enabled the network to learn the task rapidly and successfully (**Fig. 5d**). Importantly, multiplicative coupling not only produced high reward (i.e., reward probability *P*(*reward*|*choice*)), but also produced stable decision-making probability *P*(choice) (**Fig. 5e**). The reward uncertainty also affected the switch latency (**Fig. 5f**). Notably, in the case of small batch size, additive coupling produced unstable *P*(choice) curve, whereas multiplicative coupling was consistently stable (**Fig. 5g**). Lastly, we trained the E/I-PFC-MD model with actor-critic RL to perform a probabilistic Go/NoGo reversal learning task^31^ (**Fig. 5h**, Wang et al., 2023). Faster switching speed was again observed with multiplicative gating; the convergence speed reduced expectedly with an increasing level of reward uncertainty (**Fig. 5i**), and the average curve of proportion of correct responses matched the reported human experimental result (**Fig. 5j**). While the reward uncertainty negatively impacted switch latency (**Fig. 5k**), faster switch was still achieved in multiplicative gating (**Fig. 5l**). Taken together, the results from these two cognitive tasks support that multiplicative couplings improve cognitive flexibility for adapting to new contexts in a dynamic environment.

### Modeling cortico-thalamic-cortical network in working memory and attention

The higher-order thalamic nuclei such as the pulvinar has been suggested to play an important role in regulating cortico-cortical communications^32–34^ (Cortes et al., 2024; Perry et al., 2022; Kohn et al., 2020); yet the exact gating mechanism of the transthalamic pathway remains unclear for WM and attention. To model the cortico-thalamic-cortical loop, we assumed that two RNNs that represent two cortical structures (such as the PFC and PPC) have bidirectional connectivity with the FNN, such as the mediodorsal pulvinar; additionally, two fully connected RNNs have reciprocal and yet sparse cortico-cortical connections between excitatory neurons (**Fig. 6a**). Furthermore, we assumed that PPC-PFC connectivity is modulated by the pulvinar through multiplicative coupling. This is a special case of RNN-FNN coupling, where two interconnected cortical areas are viewed as a general RNN, with each subregion modeled by its own RNN dynamics and structural connectivity (**Methods**).

**Figure 6.**
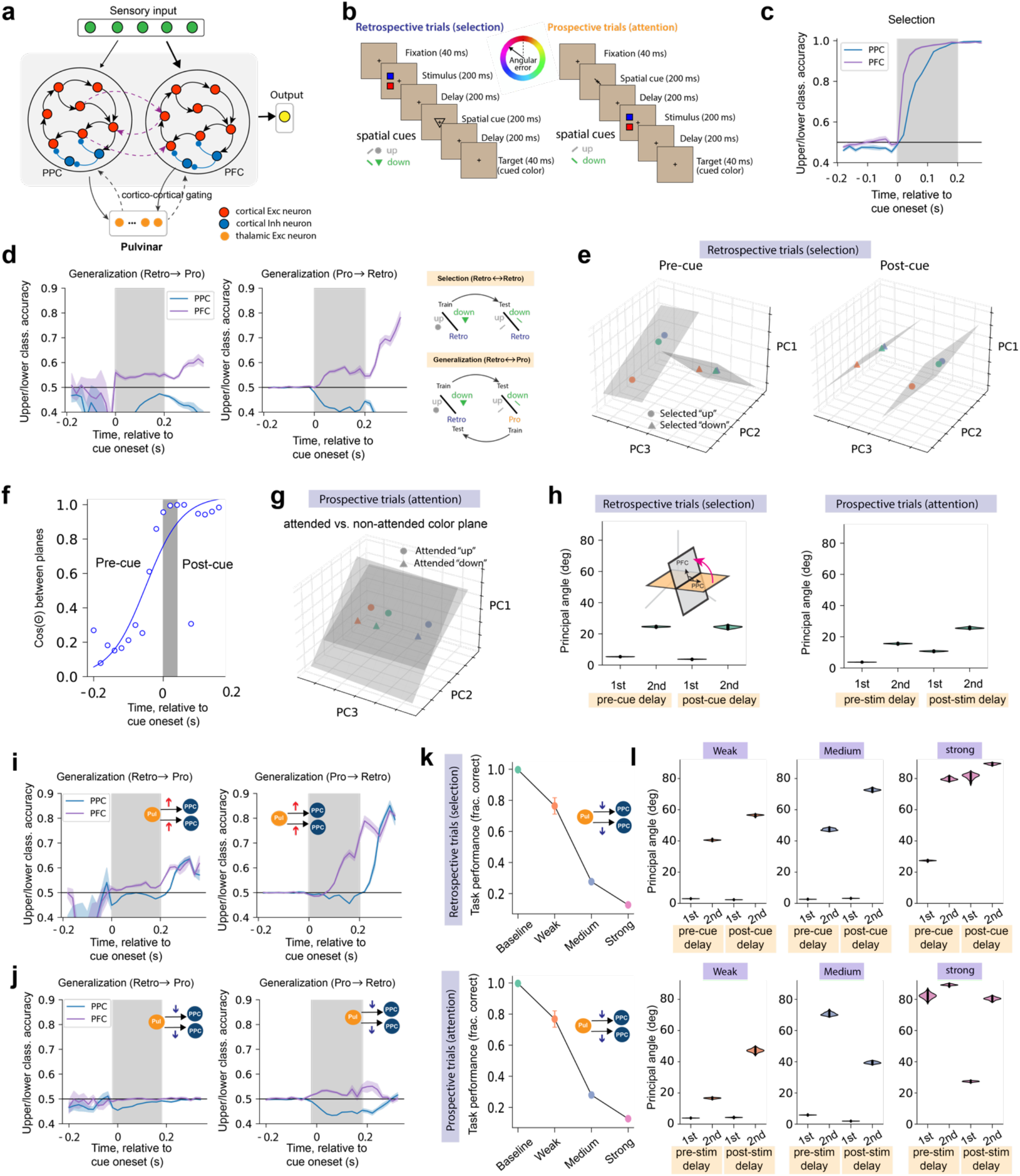
Pulvinar activity for gating cortico-cortical connectivity in working memory and attention tasks. **(a)** Schematic of the PPC-Pulvinar-PFC network, which was equipped with pulvinar activity-dependent gating for PPC-PFC connectivity. Red and blue circles denote cortical excitatory and inhibitory neurons, respectively. Dashed lines represent weaker connections or sparser connectivity. **(b)** Schematic of the modified WM and attention task adapted from REF^35^. The retrospective trials (“selection”) and prospective trials (“attention”) swapped the stimuli and spatial cues (circle/triangle or upward/downward bar), where the stimuli consisted of red and blue squares. **(c)** PFC and PPC population classification accuracy curves relative to the cue onset during selection condition. Shaded area denotes s.e.m. (n=10). **(d)** PFC and PPC population classification accuracy curves relative to the cue onset during two directed (Retro→Pro and Pro→Retro) generalization conditions. Shaded area denotes s.e.m. (n=10). Right: training and testing in selection and generalization settings based on retrospective and prospective trials. **(e)** Model prediction showed that principal subspaces between PFC subpopulation activities were orthogonal during the pre-cue period ([-200, 0] ms) and were nearly parallel during the post-cue period ([150, 200] ms) in retrospective trials. Shaded planes denote two planes represented by the selected ‘up’ and ‘down’-tuned PFC units. **(f)** Quantification of panel **d**, where every dot represents a sample computed at every time bin (dt=20 ms). Blue line shows the best-fitting logistic function for all sample points. **(g)** The angle between attended and non-attended planes in attention ‘Pro’ trials was nearly parallel. **(h)** Comparisons of the first and second largest principal angles during the pre-cue and post-cue delay periods in WM (selection) and attention tasks. Geometric illustration of principal angle was shown. Violin plots were generated (n=20). **(i)** Compared to panel **d**, activation of the pulvinar improved both PPC and PFC upper/lower classification accuracies in two generalization (‘Retro→Pro’ and ‘Pro→Retro’) settings. Shaded area denotes s.e.m. (n=10). **(j)** Compared to panel **d**, suppression of the pulvinar decreased the PFC upper/lower classification accuracy in two generalization (‘Retro→Pro’ and ‘Pro→Retro’) settings. Shaded area denotes s.e.m. (n=10). **(k)** Reducing the pulvinar impact by weakening pulvinar→PPC and pulvinar→PFC connection strengths reduced classification accuracy in both selection (top) and attention (bottom) tasks. Weak, medium, and strong reduction produced monotonic reduction is task performance. Error bar denotes SD (n=20). **(l)** Associated with performance reduction in panel **k**, weak, medium, and strong reduction of pulvinar input dramatically changed principal angles between the PPC and PFC. Two subspaces were unaligned when the principal angle became nearly orthogonal (n=20).

Motivated by the monkey PPC-PFC circuits that performed the WM and attention tasks^35^ (Panichello and Buschman, 2021; **Fig. 6b**), we trained two models to conduct retrospective (WM) and prospective (attention) tasks: (i) a PPC-Pulvinar-PFC network with multiplicative pulvinar gating on cortico-cortical connectivity (**Methods**) and (ii) a standard PPC-PFC network (without thalamus). Upon completion of model training, we examined the population representations of PPC and PFC in both WM and attention trials (**Fig. 6c,d**). To assess the emergent properties of the trained models and compare them with experimental findings from macaque monkeys, we conducted two types of computation. First, we employed a support vector machine (SVM) for population decoding analysis separately for the PFC and PPC during the selection trials (‘Retro→Retro’) and subsequently applied the trained decoders to test the generalization trials for the PFC and PPC, respectively. We computed the two-class classification accuracy in both selection and generalization modes. Second, we identified the ‘up’ and ‘down’ rule-tuned units among the PFC population and identified two two-dimensional planes separately from the ‘up’ and ‘down’ subpopulations based on principal component analysis (PCA). During retrospective trials, we further quantified the angle between the two planes before and after cue presentation (same as the procedure describe in REF^35^). Specifically, two orthogonal subspaces suggest an angle of 90 degrees, whereas two parallel subspaces imply an angle of 0 degree. Notably, the multiplicative PPC-Pulvinar-PFC model produced qualitatively and quantitatively similar results (**Fig. 6c-f**) as the experimental data (**Extended Data Fig. 8a-c**) in all decoding analyses (**Extended Data Fig. 8a**; same as Fig. 2c of REF^35^) as well as the PFC subspace relationship (**Extended Data Fig. 8b,c**; same as Fig. 4a,b of REF^35^) compared to the standard model (**Extended Data Fig. 8d**). We noticed that the PFC population showed an earlier rise than the PPC population in decoding accuracy in the selection trials (**Fig. 6c**). Furthermore, the pretrained PFC decoder performed well in the generalization trials (‘Retro→Pro’ and ‘Pro→Retro’), whereas the pretrained PPC decoder failed to generalize (for geometric insights, see the characterization of principal angles in **Extended Data Fig. 8e**). Notably, the PFC population subspaces showed a dramatic shift in their angles from the pre- to post-cue periods in retrospective trials (**Fig. 6f**), supporting that selection transformed WM representations in the PFC to match the changing task demands^35^. Interestingly, the thalamic gating strength had a reverse logistic shape before and after cue presentation (**Extended Data Fig. 8f**). Furthermore, we found that the attended and non-attended planes were nearly parallel in the attention task (**Fig. 6g** and **Extended Data Fig. 8g**), matching the experimental result (see Extended Data Fig. 10a of REF^35^). To gain more geometric insights, we measured the principal angles spanned by the PFC and PPC subspaces during delay periods in respective WM and attention tasks (**Fig. 6h**). In the WM task, we observed that the dominant principal angle was close to 0 degree during both pre-cue delay and post-cue delay, suggesting that PPC and PFC subspaces shared at least one common direction (i.e., at least one shared neural activity pattern). In the attention task, the results were quantitatively similar.

We further tested whether pulvinar activation in the PPC-pulvinar-PFC model can change the PPC-PFC angle between the ‘up’ and ‘down’ planes and affect the up vs. down classification accuracy accordingly. Remarkably, our experimental predictions showed that pulvinar activation not only substantially boosted PPC decoding accuracy from the chance level to a comparable PFC decoding accuracy level in both ‘Retro→Pro’ and ‘Pro→Retro’ settings but also improved the rise speed in PFC decoding accuracy curve (**Fig. 6i**). The improvement in decoding accuracy also confirmed our geometric intuition about the principal angle between the PPC and PFC subspaces. In contrast, pulvinar suppression diminished the PFC decoding accuracy in generalization trials (**Fig. 6j**).

In model prediction, we also found that weakening the pulvinar→PPC and pulvinar→PFC connectivity strengths reduced the final task performance in both WM and attention tasks (**Fig. 6k**) and dramatically changed the two principal angles (**Fig. 6l**), especially the second largest principal angle. There was a mirror pattern in principal angles between the pre-cue (or post-cue) delay in retrospective trials and the post-stimulus (or pre-stimulus) delay in prospective trials. These perturbation experiments showed that pulvinar suppression caused some dimensions to become partially or completely unaligned, whereas pulvinar activation dynamically modulated thalamocortical and cortico-cortical connectivity to align the PPC-PPC subspaces.

In addition to these model predictions (**Fig. 6i-l**), multiplicative gating of cortico-cortical connectivity substantially improved the task learning speed, varying from ~2.5 to more than 10 folds (**Extended Data Fig. 8h**). Put together, these results strongly support that multiplicative thalamic gating through the pulvinar plays a role in aligning two cortical subspaces for mediating cortico-cortical communications and coordinating long-range connected cortical neurons in learning.

### Modeling an entorhinal-hippocampal network in visuospatial navigation

To demonstrate the benefit of multiplicative gating beyond the thalamocortical network, we further considered a hierarchical RNN-FNN network that resembles the entorhinal cortex-hippocampus (EC-DG-CA3-CA1) architecture involving multiple recurrent and feedforward modules (**Fig. 7a**): EC and CA3 have densely recurrent connections, whereas DG and CA1 have little or no recurrent connections. In the entorhinal-hippocampal network, we assumed that EC received visual and velocity inputs as the primary sources of information before propagating them to the hippocampus. Both EC and CA3 were modeled as RNNs, while DG and CA1 were modeled as FNNs. We assumed that (i) the connections between CA3 and CA1 follow a topographically organized pattern^36,37^ (Brivanlou et al., 2004; Hongo et al., 2015); (ii) the EC→DG and EG→CA3 pathways show sparse connectivity^38,39^ (Yassa and Stark, 2011; Rebola et al., 2017); (iii) bidirectional EC-CA1 connections were sparsely connected. We trained the EC-DG-CA3-CA1 network with either additive or multiplicative interactions between EC and CA1 to perform a two-dimensional (2D) visuospatial navigation task^13^ (Zhang et al., 2022; **Methods**). The hierarchical network integrated both velocity and visual signals to produce an output of 2D position of the simulated trajectory.

**Figure 7.**
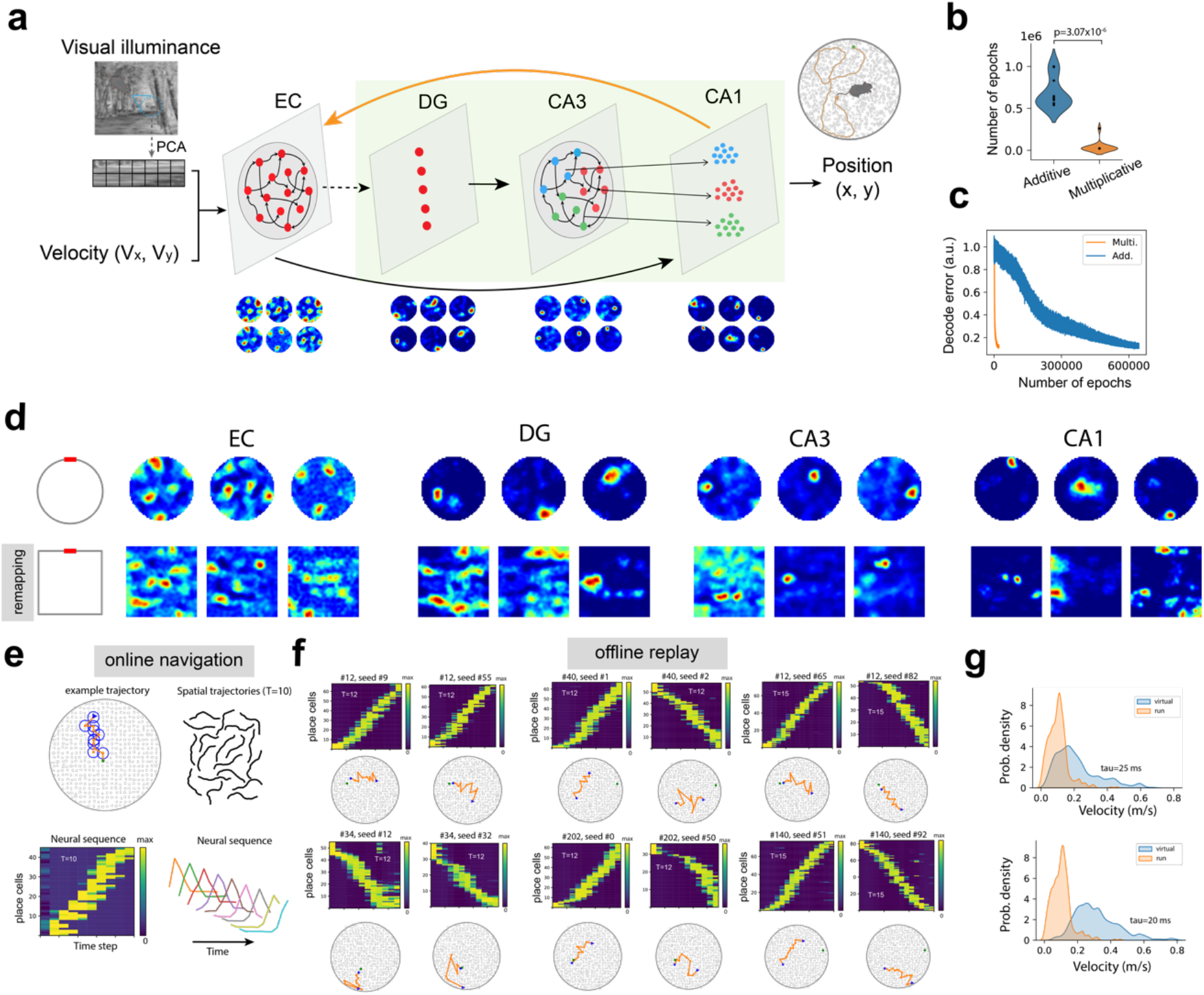
The entorhinal-hippocampal network in the visuospatial navigation task and offline sequence replay. **(a)** A schematic of the EC-DG-CA3-CA1 circuit model used for a two-dimensional spatial navigation task with multisensory cues. Visual cues were represented by principal components from an (8×8)-by-(8×8) image patch, while spatial cues were conveyed through two-dimensional velocity inputs (V_x_, V_y_). Both visual and spatial inputs were fed to EC, whereas the output pooled together the activity of three hippocampal modules (DG, CA3 and CA1) to generate an estimate of the 2D position. The EC has reciprocal connectivity with CA1 and has weak connectivity with DG, and CA3 and CA1 have topographically aligned connectivity (coded by three colors). Two models were trained, one with assumed multiplicative EC-CA1 coupling, and the other with additive EC-CA1 coupling. In the trained model with multiplicative coupling, grid-like activity patterns emerged in the EC population, whereas place cell-like firing patterns emerged in the DG, CA3, and CA1 populations. **(b)** Comparison of convergence speed between multiplicative and additive couplings (n=20, rank-sum test, p= 3.07×10^−6^). **(c)** Representative convergence curves for training the entorhinal-hippocampal network with multiplicative (blue) versus additive (orange) coupling, measured by the position decoding error. **(d)** Upon switching from a circular to a square-shaped environment, remapping of firing rate patterns emerged in EC, DG, CA3, and CA1. **(e)** Illustration of a sequence of activation of CA1 place cells during visuospatial navigation. Top left: trajectory overlaid with overlapping place fields. Top right: examples of spatial trajectories used in training (sequence length: 10). Bottom left: neural sequence ordered by the activation latency of place cells. Bottom right: nine sorted place cells in sequence, along with their mean firing rate profiles. **(f)** Representative examples of CA1 reactivated sequences during an off-line state. Multiplicative coupling showed various temporal sequences triggered by a random noise input. Each curve represents a virtual trajectory (sequence length: 12-15); green circle denotes the agent’s initial position, blue star and triangle symbols denote the start and end positions of the replay trajectory, respectively. The corresponding neural sequences of selected place cells were visualized as heat maps and the cell order was ranked by a consistent order (yet different units were reactivated among the CA1 population), where each row represented the normalized firing activity of a place cell and was sorted based on the peak latency. **(g)** Distribution statistics of virtual speed of sequence replay based on two different time constants. In comparison, a smaller time constant led to a faster replay speed.

We first noted substantial (up to ~100 folds) learning speed improvement using multiplicative EC-CA1 coupling (**Fig. 7b,c**; p= 3.07×10^−6^, rank-sum test), whereas additive EC-CA1 coupling sometimes (1 out of 20 runs) failed to converge within a given maximum number of epochs. Examining the emergent tuning properties of units at different layer of the hierarchy showed grid cell patterns in EC and localized place cell patterns in CA1, consistent with some early findings of the RNN modeling literature^40,41,13^ (Cueva and Wei, 2018; Sorscher et al., 2023; Zhang et al., 2022). Emergent place cell patterns also appeared in both DG and CA3. We further tested the impact of the change of environment on individual unit tunings and found that changing the shape of environmental enclosure resulted in various degrees of unit receptive field remapping across the network hierarchy (**Fig. 7d**), matching the experimental findings^42^ (Colgin et al., 2008). Additionally, CA3 units in the EC-CA1 multiplicative coupling model showed more stable dynamics than CA1 units under the same input perturbations, as evidenced in the stability of population decoding accuracy (**Extended Data Fig. 9a,b**). This was possibly due to the connectivity difference between the ‘slow-weight RNN’ and ‘fast-weight FNN’. Experimentally, the CA3 place fields were reported to be more stable during plasticity than the CA1 place cells^43^ (Dong et al., 2021).

During spatial navigation, CA1 place cells exhibited sequential activation based on their place field locations (**Fig. 7e**). To model the offline (sleep or immobile wakefulness) state, we turn off the sensory input and presented only a small random noise input to EC given a random initial position. We also reduced time constant to accelerate the speed of replay. By examining the internal representations in CA1, first, we found virtual ‘forward’ and ‘reverse’ neuronal sequences generated from intrinsic multiplicative coupling dynamics, with the sequence length (range: 10-15) matching or longer than the training sequence length. Some sequences were local replay (i.e., near the initial position), whereas others were remote replay (i.e., distant from the initial position). Second, ~8-16% of total CA1 units were reactivated during sequence replay, reminiscent of hippocampal theta and ripple sequences. Third, given the same initial position condition, different noise inputs or random seeds could trigger distinct sequential activation of randomly selected CA1 place cells (**Fig. 7f**). Conceptually, the replay sequence generation can be represented as a function of multiple factors: *f*(*τ, noise, rand*_*seed, init*_*pos*), and the replay sequence length is determined by the intrinsic memory capacity. Fourth, from all CA1 replayed trajectories, we estimated the average virtual replay speed (mean±SD: 0.21±0.13 m/s for *τ* = 25 ms; for 0.31±0.13 m/s *τ* = 20 ms), which was ~4-5 times faster than the mean run speed (mean±SD: 0.08±0.01 m/s) (**Fig. 7g**). Fifth, when CA1 units exhibited replay, CA3 units also displayed synchronous and coordinated sequence replay showing similar or dissimilar spatial trajectories (**Extended Data Fig. 9c**); additionally, the timing of CA3 unit reactivation was (~1 temporal bin) faster than the onset of CA1 unit reactivation (**Extended Data Fig. 9d**), consistent with the notion that CA3 driving CA during hippocampal ripple events^44^ (Nakashiba et al., 2009). Sixth, and most importantly, the control model with additive EC-CA1 coupling did not yield any meaningful neural sequence structure during the offline state (results not shown). Put together, these neural circuit modeling applications (**Figs. 5–7**) demonstrate the strengths and flexibility of multiplicative coupling in model prediction and validation of experimental findings.

## DISCUSSION

Our proposed multiplicative gating mechanism has several conceptual links to several other neural gating models proposed in the neuroscience literature. First, early visual processing involves complex recurrent and feedforward structures and bidirectional loops in the visual retino-engicute-V1 hierarchy. One theory of visual attention^45^ (Olshausen et al., 1993) suggested that the pulvinar may serve as a control signal to dynamically route information through visual cortices. In the attention model, each control unit in the pulvinar modulates a single or local group of synapses; the output node of the routing circuit provides an input to associative memory, and the output of associative memory is then fed back and correlated with the input to drive the control units. According to the proposal of multiplicative gating of cortical information flow^45,46^ (Koch and Poggio, 1992; Olshausen et al., 1993), a control thalamic unit may modulate the strength of a cortico-cortical synapse through nonlinear neuronal-synaptic gating.

Second, our notion of activity-dependent plasticity in RNN is conceptually in line with the proposal of ‘dynamic synapses’ of the MPN^26,47^ (Aitken and Mihalas, 2023; Aitken et al., 2024) or even the early generic concept^48–50^ (Tsodyks et al., 1998; Dror and Tsodykis, 2000; Barak and Tsodyks, 2005). It was shown in the mean-field theory framework that the addition of dynamic synapses to RNN introduces sharp increases in population spikes as well as sensitivity to the input duration during the WM delay period^50^ (Barak and Tsodyks, 2005), revealing transient synchronization and persistent attractor dynamics that matched experimental findings in PFC neurons and motor cortical neurons in the delay periods^51,52^ (Wang et al., 2006; Riehle et al., 1997). The prediction of our model with multiplicative gating (**Fig. 2e**) also matched this observation. In the MPN framework^26^, the feedforward synaptic weight matrix was assumed to be multiplicatively gated by a synaptic modulation matrix; the MPN showed slower multi-plasticity attractor dynamics different from vanilla RNNs and achieved comparable task performance in integration tasks. Our work, inspired and developed independently from thalamocortical gating, is a generalization of this concept. Similarly, we showed that multiplicative synaptic plasticity in RNN-FNN models produces emergent rich neural dynamics (such as fixed points, line attractors, damped oscillations, and limit cycles, **Fig. 2d,e**) during WM, which requires neither oscillatory inputs nor two coupled oscillatory populations^53^ (Pals et al., 2024). Additionally, we showed that multiplicative gating within RNN-FNN enables rapid switching or context-dependent computation (**Fig. 6**). This opens new doors to theoretical analyses of dynamic synapses, neural state hopping, and multi-plasticity network dynamics.

Third, our proposed multiplicative coupling mechanism shares the same philosophy with the neuron-synapse coupling theory, which posits that synapses influence neurons by shaping network dynamics and neurons influence synapses through activity-dependent plasticity^54–57^ (Clark and Abbott, 2024; Haydon and Carmignoto, 2006; Krishnamurthy et al., 2022; Todo et al., 2019). Our proposal also supports the synaptic gating theory^58^ (Katz, 2003). The goalkeeper neurons can influence information transmission, commonly observed in the thalamus, hippocampus and basal ganglia^59,60^ (Boukadoum and Gisger, 2011; Foresco and Grace, 2003). Furthermore, multiplicative interactions between slow-weight and fast-weight dynamics have been proposed in dynamic synapses of the cortex^50^ (Barak and Tsodyks, 2007) and in ‘tripartite synapses’ based on synaptic-astrocyte coupling^61^ (Gong et al., 2024).

More generally, multiplicative interactions are computationally powerful in modeling gating, attention, feature fusion, and integration mechanisms through elementwise multiplication or Hadamard product^16,62^ (Jayakumar et al., 2020; Wu et al., 2016). For example, attention can act as a gatekeeper and filter to influence memory and determine what information is processed in memory processing. The concept of multiplicative gating has also been explored in several versions of discrete-time RNNs, including variants of Long Short-Term Memory (LSTM)^63^ (Hochreiter and Schmidhuber, 1997) and Gated Recurrent Unit (GRU) network^64^ (Chung et al., 2014), which employ gating mechanisms to mitigate the vanishing gradient. Specifically, the gating mechanism uses neurons that multiply the outputs of other neurons---the so-called ‘multiplicative units’; such multiplicative interactions allow the input or feedback activity to dynamically modulate effective RNN connectivity. These gating mechanisms allow for flexible control of information flow and network dynamics through time. The input-gating based multiplicative RNN (mRNN) architecture assumes the recurrent weight matrices as a function of the input and represent the hidden state transition as a three-way tensor of input-dependent connections. Therefore, mRNN may gate the information through learnable multiplicative interactions so that every input can synthesize its own recurrent weight matrix by determining the gain^59,65–67^ (Sutskever et al., 2011; Sussillo et al., 2016; Wu et al., 2016; Krause et al., 2017). Multiplicative interactions were also adopted in the deep, feedforward Transformer architecture^68^ (Vaswani et al., 2017), which amplifies or suppresses different parts of the input for attention and efficiently models long-range sequence dependencies. Combining different multiplicative gating mechanisms within the RNN-FNN structure is also possible. For instance, FNN can incorporate self-attention and layer normalization operations as well as multiple layer structures to learn high-dimensional embeddings^3^ (Kozachkov et al., 2023).

The RNN-FNN architecture is computationally appealing for several reasons. The idea of using two networks (either two FNNs or one RNN plus one FNN) for learning temporal sequences has a deep root in machine learning^17–19^ (Hinton and Platt, 1987; Schmidhuber, 1992; Ba et al., 2016), such that the first network learns to produce context-dependent weight changes for the second network, and FNN produces an output transformed into intermediate weight changes for the fast-weight network which facilitates rapid synaptic changes or few-shot learning.

### Insight into NeuroAI

Hierarchical RNN-FNN networks with multiple recurrent and feedforward loops are omnipresent in biological brains. Neurons within cortical networks are recurrently connected, whereas neurons within many subcortical structures were loosely connected. FNN provides a contextual input that may rapidly changes the RNN state through multiplicative coupling. Through dynamic RNN connectivity masking, information is regulated between recurrent units based on their correlative activities--- a neuronal doctrine reminiscent of Hebbian plasticity. Through the feedback-gating mechanism, context information-rich FNN can serve as a control signal and impose attention through history-dependent associative memory. A growing line of evidence has suggested that the thalamus plays a role of gate controller or gatekeeper for the cortex, ranging from sensory gating, attention, to thalamocortical and cortico-cortical information regulation^69–71^ (McCormick and Bal, 1994; Gisiger and Boukadoum, 2011; Halassa and Sherman, 2021), where the searchlight of attention was modulated by the inhibitory thalamic reticular nuclei (TRN)^72–74^ (Crick, 1984; Hassaa et al., 2014; Chen et al., 2015). The thalamus also provides contextual feedback to the cortex and routes information thorough reciprocal thalamocortical and corticothalamic loops. The cortico-thalamic-cortical loop involving the bidirectional interactions between the recurrent PFC and the feedforward MD thalamus is crucial for flexible cognitive control^75^ (Scott et al., 2024). As a plausible form of multiplicative thalamocortical interactions, the thalamus output can influence the cortical state through a specific pathway that targets on cortical vasoactive intestinal peptide (VIP)-expressing interneurons. The VIP-targeted MD neurons with dopamine receptor-expression may amplify prefrontal signals and regulate the strength of functional connectivity within the PFC via local disinhibitory circuit motifs^76^ (Mukerjee et al., 2021). Furthermore, optogenetic MD activation can amplify local PFC connectivity between paired excitatory neurons and enable rule-specific neural sequences to emerge, whereas MD suppression reduces PFC connectivity between excitatory neurons (see Fig. 3 of REF^30^). By modulation of interneurons, the MD can act as a source of gatekeeping^60^ (Floresco and Grace, 2003). Accumulating lines of experimental evidence from mice and tree shrews^30,76,77^ (Schmitt et al., 2917; Mukerjee et al., 2021; Lam et al., 2025) have led us to hypothesize that multiplicative thalamocortical couplings allow the PFC to integrate context-rich thalamic signals and maintain robust WM, enabling context-dependent decision-making. Consistent with our prior thalamocortical modeling results^14^ (Zhang et al., 2025), the current model prediction (**Fig. 2e**) lends additional support for this hypothesis. The extension to multiplicative cortico-pulvinar-cortical gating for attention and information routing further generalizes this notion, supporting that the role of pulvinar in regulating cortico-cortical communication^34,78^ (Halassa and Kastner, 2017; Kohn et al., 2020). Our model predictions (**Fig. 6i-k**) on pulvinar manipulation provide new experimentally testable hypotheses that call for future experimental validation. The same computational principle is likely to apply to the thalamocortical system in the motor cortex-motor thalamus-basal ganglia loop in motor planning or control^79,80^ (Bosch-Bouju et al., 2013; Logiaco et al., 2021).

Multiplicative gating finds its efficacy while modeling the hierarchical entorhinal-hippocampal system. Not only multiplicative gating demonstrated rapid learning in the visuospatial navigation task, but it also showed neuronal tuning and offline replay matching experimental findings. Our proposed model has the capability of producing forward and reverse replay-like sequences in the CA1 and CA3 modules in the absence of sensory input, offering a unifying framework to study neural representations during online and offline learning for spatial and non-spatial tasks. The task-optimized multiplicative gating-empowered entorhinal-hippocampal network can be viewed as a continuous attractor network (CAN)^81^ (Khona and Fiete, 2022) that produces internal generation of an experienced or novel sequence or spatial trajectory. Since the reactivated CA1 units from each replay event are random samples of the whole CA1 population, they can also contribute to off-line learning through Hebbian plasticity. To date, experimental evidence on the functional roles of CA1 feedback (through the subiculum) in gating EC has been reported^82,83^ (Basu et al., 2016; Haam et al., 2018). Of note, direct evidence of CA1 gating on EC synapses requires future investigations.

Flexible and robust context-dependent computation is a hallmark of the brain across various cognitive, sensory and motor tasks^9,84,85^ (Mante et al., 2013; Remington et al., 2018; Qi et al., 2025). Nevertheless, the underlying neural basis or mechanism remains incompletely understood^86^ (Heald et al., 2023). One of these proposals is to control a multiplicative gain of the input^84^ (Remington et al., 2018) or control the effective connectivity or dynamics of the network^80^ (Logicaco et al., 2021). The context information, being latent in the collected neural data, possibly explains a large degree of variance in neuronal firing variability. Detailed mathematical analyses of synaptic-neuronal coupling^47,52^ may further reveal insights into multiscale synaptic plasticity and attractor dynamics.

Multiplicative interactions may serve as a unifying framework for a wide range of ANN computational motifs, such as gating, attention layers, hypernetworks, and dynamic convolutions^16^ (Jayakumar et al., 2020). Low-rank matrix factorization or low-rank RNN connectivity also assume representations based on multiplicative interactions. Hebbian activity-dependent synaptic gating unifies the connectivity and activity-dynamics and promotes stability in various forms of learning. Such paradigm-independent advantages in learning are useful for modeling multiregional and hierarchical circuit structures or training variants of RNNs to perform cognitive and control tasks.

Multiplicative couplings offer a flexible framework for rapid, stable, and flexible learning. Our empirical observations of improved model generalization on the held-out data (**Fig. 1d** and **Fig. 4f**) also suggest its implicit role in adaptive and dynamic weight regularization. Another benefit of multiplicative gating is to facilitate few-shot learning, enabling agents to explore dynamic environments or adapt to new tasks quickly based on limited trials or samples. An effective strategy for achieving few-shot learning is through meta-learning (a.k.a. ‘learning to learn fast’)^87,88^ (Tyulmankov et al., 2022; Shervani-Tabar and Rosenbaum, 2023), where FNN acts like a meta-learning engine to learn new contexts rapidly. In the context-switching task, RNN plays the role of pretrained base learner (‘slow weights’) for performing the desired task, whereas the meta-optimizer adapts ‘fast weights’ to modulate FNN activity and subsequently changes the context-dependent RNN dynamics. In a general setting, the mask matrix **L** can be parameterized by a low rank matrix (**Supplementary Note 1**). Depending on the task demand, the low-rank modulated network can learn ‘what’ and ‘where’ to gate, fulfilling the respective attention and routing roles. Therefore, this multiplicative gating framework can generalize low-rank RNNs^89^ (Dubreuil et al., 2022) and expand the representation power for continual learning or task-dependent computation.

Finally, our findings suggest new paths to build biologically realistic RNN-FNN architectures with enhanced functionality for modeling complex multi-area brain structures. Take the thalamus as an example, the MD thalamus receives either direct or indirect dopamine input from the ventral striatum, a subcortical rewarding area that is important for both goal-driven and habitual behaviors; integration of thalamo-striatal and cortico-striatal systems^90^ (Haber and Calzavara, 2009) into the cortico-thalamic-cortical loop may produce a modeling paradigm characterizing behavioral deficits in both aversive motivation regulation and cognitive control^91^ (Grahek et al., 2019). Furthermore, cortico-cortical connectivity can be modulated jointly by the thalamus (such as the MD and pulvinar) and top-down modulatory input in a multiplicative manner (e.g., **W**^rec^⨀**L**_1_⨀**L**_2_), where **L**_1_ and **L**_2_ denote two masks representing two independent multi-timescale gating mechanisms. Additionally, multiplicative gating may control interareal information flow in a brain state-dependent manner. Take hippocampal-neocortical interactions as an example, we may envision that recurrent intracortical connectivity is gated by fast-weight hippocampus-driven auto-associative memory (ASM) during online task learning; whereas in an off-line memory consolidation state, the connectivity can be gated by top-down cortico-hippocampal activity (e.g., long-range VIP+ interneurons projections from the neocortex to CA1^92,Malik et al. 2022^) based on hetero-associative memory (HSM) to coordinate bidirectional communications. Therefore, circuit modeling with multiplicative gating may serve as a starting point to investigate various interesting hypotheses and provide experimentally testable predictions.

## METHODS

### Benchmark tasks

#### Working memory (WM) and decision-making (DM) benchmarks

We adopted five established cognitive benchmark tasks (Yang et al., 2019): *Go/NoGo, Decision Making (DM fixed), Delayed Decision Making (DM RT) Multisensory DM* and *Mante*^9^ (Mante et al., 2013), where were adapted from REF^11^(Yang et al., 2019). The Python software implementation is available at https://github.com/gyyang/multitask.

#### Continuous-drift integration task

We adapted and altered the “simple integration task”^26^ (Aitken and Mihalas, 2023). In this task, the network was aimed to determine whether the mean of a noisy one-dimensional stimulus sequence was positive or negative. In the training set, half of the sequences had negative means (class 1), uniformly sampled from [-0.12, 0]; the other half had positive means (class 2), sampled from [0, 0.12]. Within an input sequence, each stimulus input was drawn from a Gaussian distribution centered at the sequence mean, with variance depending on the sequence length 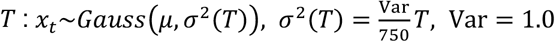. This choice was to keep the total signal-to-noise ratio (SNR) independent of *T*. The final input is a fixed ‘Go’ signal that prompts the RNN-FNN model to output its decision. To perform the two-class integration task, we trained a total of 60 RNN-FNN models (30 with multiplicative coupling, 30 with additive coupling) on 800 training sequences of constant length *T* = 50 and tested on 200 testing sequences, each with a varying length 50, 100, and 200. We initialized trainable weights {**W**^rec^, **W**^FNN→RNN^, **W**^RNN→FNN^} from *Gauss*(0, 0.1), and set untrainable **W**^in^ = [1, 1, 1]^*T*^. We used the logistic sigmoid activation (instead of ReLu) for improving the stability of learning. The integration timestep and time constant were set to *dt* = 0.1 and *τ* = 2.0 which ensured sufficiently continuous dynamics while enabling stable training for both coupling types. Models were trained on 800 training sequences (length *T* = 50) and evaluated on 200 testing sequences of lengths 50, 100, and 200. Furthermore, we investigated the effect when a pure-noise delay (delay length: *T*_delay_) was inserted at the midpoint of test sequences; the noise during this delay period was set to have the same variance as the rest of the sequence.

#### Cart-pole and mountain-car benchmarks

In the context of RL, we studied several continuous control benchmarks. Cart-pole, also known as inverted pendulum problem, is a standard neural controller benchmark^93^ (Geva and Sitte, 1993). The lower end of a pole is mounted on a cart in such a way that the pole can only swing in a vertical plane parallel to the direction of motion of the cart. To balance the pole the cart is pushed back and forth on a track of limited length, and it has become a benchmark to evaluate specific RL algorithms for continuous control^94^ (Barto et al., 1983). The state of the cartpole system is described by four variables: the horizontal position of the cart *x*, its velocity 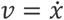, the pole angle *θ*, and the angular velocity 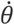. The force *F* applied to the cart provides the controlling action. In terms of Newtonian mechanics, the equations of motion are described by two equations

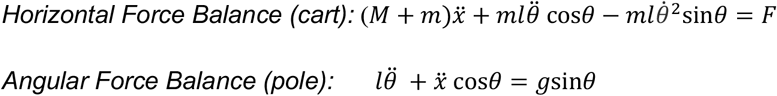

where *M* and denote *m* the mass of cart and pole, respectively. The cart and pole dynamics interact in a nonlinear and coupled manner. The second derivatives of the motion, namely the cart acceleration 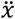 and angular acceleration 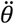, are described as follows

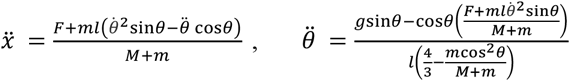

RL is aimed at learning the approximation of state-action value function *Q*(***s, a***). The state set consists of the 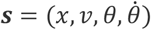 of the cart, the action set *a* consists of three options: “move left”, “move right”, and “no action”. The reward function is given by 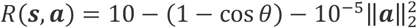. The trial terminates when |*x*| > 2.4 or |*θ*| > 0.2.

Mountain-car is a classic benchmark for testing the performance of RL algorithms. In this problem, an underpowered car must climb a steep hill to reach a goal located at the top of the hill. The state of the Mountain-car environment is represented by two continuous variables, the position *x* and velocity 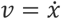 of the car. The action space is discrete (“accelerate forward”, “accelerate backward”, and “no action”), and the positive reward is related to the car’s vertical offset, such as *R*(***s, a***) = −1 + height. The agent also receives a negative reward for every time step it takes to reach the goal (i.e., the target height). The agent is aimed at learning how to control the car’s acceleration to climb the hill and reach the goal as quickly as possible while using the least amount of energy. During testing trials, we also monitored the accumulated reward, if the reward value was below −200, suggesting that failed convergence after 200 trials.

#### MNIST benchmark

The MNIST (Modified National Institute of Standards and Technology) dataset consists of 70,000 28 × 28 pixels grayscale images of handwritten single digits between 0 and 9. The digits have been size-normalized and centered in a fixed-size image. The goal is to train the images into ten categories using 60,000 training samples (a subset of 10,000 samples for validation) and test the model with additional 10,000 held-out samples. The dataset is publicly available (http://yann.lecun.com/exdb/mnist/).

#### IMDb Movie Reviews benchmark

The IMDb benchmark is a binary sentiment analysis dataset consisting of 50,000 reviews from the Internet Movie Database (IMDb). The dataset contains an even number of positive and negative reviews. Only highly polarizing reviews are considered. A negative review has a score ≤ 4 out of 10, and a positive review has a score ≥ 7 out of 10. No more than 30 reviews are included per movie. The dataset is publicly available (https://paperswithcode.com/dataset/imdb-movie-reviews). For the input representation, we adopted a pretrained word2vec model to obtain the 300-dimensional word embedding vectors.

#### TREC-6 benchmark

The Text Retrieval Conference (TREC) benchmark is a dataset used for six-class classification in machine learning, where the classes include ‘human’, ‘location’, ‘numeric information’, ‘entity’, ‘abbreviation’, ‘description and abstract concepts.’ The training set contains 5,452 questions, and the test set contains 500 questions. The dataset is publicly available (https://cogcomp.seas.upenn.edu/Data/QA/QC/).

#### CIFAR-10 benchmark

The CIFAR-10 (Canadian Institute for Advanced Research, 10 classes) benchmark is a multi-class classification task in machine learning. The dataset consists of 60000 32×32 color images in 10 classes (‘Airplane’, ‘Automobile’, ‘Bird’, ‘Cat’, ‘Deer’, ‘Dog’, ‘Frog’, ‘Horse’, ‘Ship’, ‘Truck’), with 6000 images per class. There are 50,000 training images and 10,000 test images. The dataset is divided into five training batches and one test batch, each with 10,000 images. The test batch contains exactly 1000 randomly selected images from each class, and the training batches contain exactly 5000 images from each class. The dataset is publicly available (https://www.cs.toronto.edu/~kriz/cifar.html). For the input representation, we preprocessed the image with a three-layer convolutional neural network (CNN) and fed the 64×8 features into the RNN. To process the 32×32×3 input tensor, the first and second hidden layers of the CNN consist of 2D convolutional filters (Conv_1: 3×32, Conv_2: 32×64, 3×3 kernel, padding 1) followed by batch normalization, ReLu activation function, and 2×2 max-pooling. The final layer consists of flattened features that are sent to a fully connected neural network with ReLu activation.

### Supervised learning for the RNN-FNN model

The RNN-FNN model with multiplicative or additive coupling was trained by minimizing a loss function with back-propagation through time (BPTT). We used the mean-squared-error (MSE) loss function with *L*_*2*_ regularization and default regularization parameter of 0.1. The model parameters were optimized using Adam optimizer, with default configuration of hyperparameters (learning rate parameter 0.0005; weight decay regularization parameter 0.1). The optimization procedure continued until the desired accuracy was reached. The recurrent connection matrices were initialized using Kaiming initialization^95^ (He et al., 2016), the input weights were initialized using a uniform distribution between −1 and 1, and the output weights were initialized with a Gaussian distribution of mean 0 and standard deviation 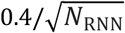. The convergence criterion was set as either the validation loss was below a preset threshold, or the test accuracy was above a threshold (e.g., 98%). The ratio of training and validation sample size varied according to the specific benchmark. In BPTT, we monitored and computed the magnitude of gradients during training using the visualization tool “TensorBoard” from PyTorch. The *L*_1_ norm of the gradient strength was plotted to compare between the additive and multiplicative couplings.

### Policy gradient and actor-critic RL algorithm

The policy gradient is an approach to solve RL problem, which targets at modeling and optimizing the policy directly. Without loss of generality, let *π*_*θ*_(*a*|*s*) denote the stochastic policy modeled by a parameterized function with respect to *θ*, which takes the state of the environment as the input and gives the actions’ probabilities as the output. The objective function *J*(*θ*) is written as

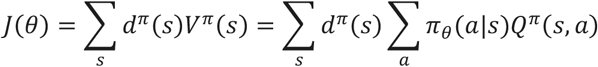

where *d*^*π*^(*s*) denotes the stationary distribution of Markov chain for *π*_*θ*_; *V*^*π*^(*s*) or *Q*^*π*^(*s, a*) denote the value of state *s* or state-action (*s, a*), respectively when we follow a policy *π*; and *V*(*s*) or *Q*(*s, a*) denote the state and state-action value functions measures the expected return of state *s* or state-action (*s, a*), respectively. We directly optimized the policy gradient ∇_*θ*_*J*(*θ*) = *E*_*π*_[*Q*^*π*^(*s, a*)∇_*θ*_ln*π*_*θ*_(*a*|*s*)] using gradient descent, where the expectation is defined with respect to the policy. The policy gradient theorem^96^ (Sutton and Barto, 2018) establishes the theoretical foundation for various policy gradient algorithms. The actor-critic model is one type of policy gradient RL algorithm that combines aspects of both policy-based methods (“actor”) and value-based methods (“critic”), which aims to address the limitations of each method when used individually. In the actor-critic framework, the actor learns the policy *π*_*θ*_(*a*|*s*) to make decisions, and the critic estimates the state or state-action value function.

In the WM and DM benchmarks, we assumed that each task had the following structure: a fixed period (1 timestep), stimulus period (25 timesteps), and decision period (10 timesteps). We employed an Advantage Actor-Critic (A2C) RL framework to optimize RNN-FNN model. Let *T* denote the maximum length of a trial. At each time step *t* ∈ {0,1, …, *T* − 1}, the RNN received input stimulus *s*_*t*_ and generated two outputs: a policy distribution *π*(*a*_*t*_|*s*_*t*_; *θ*) over discrete actions (“choose left”, “choose right”, or “maintain fixation”) via a softmax layer (“Actor”), and a state value function *V*(*s*_*t*_; *θ*_*v*_) estimate via a linear readout (Critic). The computation objective was to learn a policy that maximizes the expected cumulative reward:

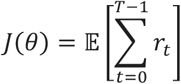

where *r*_*t*_ denotes the reward at time *t*.

The actor loss was computed using the policy gradient method:

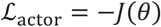

The gradient of ∇_*θ*_*J*(*θ*) for each trial was estimated using the A2C algorithm:

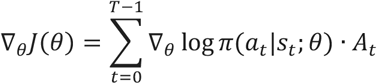

where the advantage *A*_*t*_ was given by the temporal-difference (TD) error or reward prediction error (RPE):

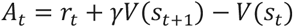

with discount factor *γ* = 0.99. The value function is further updated

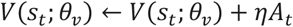

where *η* denotes the learning rate parameter.

The critic loss minimized the squared TD error:

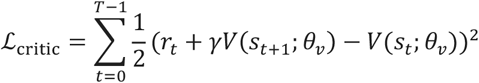

To encourage exploration, we added an entropy regularization term:

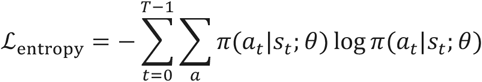

The total loss per trial was a weighted sum of the three terms:

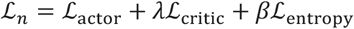

where *λ* = 1 and *β* = 0.001 are scaling coefficients for the critic and entropy terms, respectively. The final loss function was averaged across all *N*_*trials*_ trials:

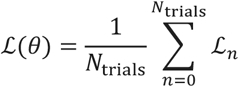

We optimized the parameters *θ* using the Adam optimizer (learning rate *η* =0.0001), with gradients computed via BPTT. To stabilize training and avoid exploding gradient in the recurrent dynamics, we applied gradient clipping with a maximum norm of 0.9.

### SRNNs for line detection and MNIST tasks

#### Neuron model

We considered two types of neurons: the standard non-coupled Leaky Integrate-and-Fire (LIF) neuron and the multiplicatively self-coupled Leaky Integrate-and-Fire (MLIF) neuron. For both types of neurons, if the membrane potential 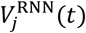 of the *j*-th hidden unit of the SRNN at timestep *t* reaches the threshold *V*_threshold_, the *j*-th neuron will emit a spike and 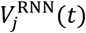 will be reset to *V*_reset_. For both LIF and MLIF neurons, the membrane potential 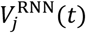 evolves according to the following differential equation:

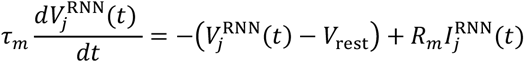

where *τ*_*m*_ is the membrane time constant, *V*_rest_ denotes the resting membrane potential, *R*_*m*_ denotes the membrane resistance, and 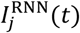 denotes the total synaptic input current to the *j*-th hidden unit at timestep *t*. The definition of 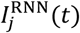 differs between the LIF and MLIF neuron models as follows:

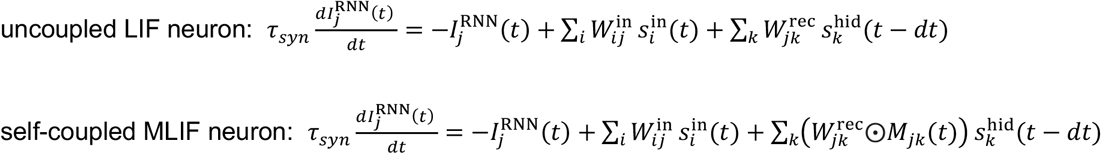

where *τ*_*syn*_ denotes the synaptic time constant; 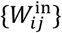 and 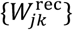 denote the input-to-hidden and hidden-to-hidden synaptic weights, respectively; 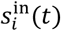 denotes the input spike received by the *i*-th input neuron at timestep *t*, and 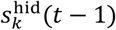 denotes the output spike emitted by the *k*-th hidden neuron at timestep *t* − 1; all binary spikes are generated by hard-thresholding the voltage against a voltage threshold *V*_threshold_.

In the MLIP neuron model, **M** = {*M*_*jk*_} denotes the time-embedded associative memory of the *j*-th hidden neuron that is defined as

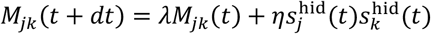

where (*λ, η*) denotes two hyperparameters, and their relative strengths determine the decay of the memory trace.

#### Network architecture and weight initialization

In the line detection task, we used a simple SRNN architecture with various sizes of hidden units (*N*_RNN_ = 30,50,100). For reproducibility, we set the random seeds for both PyTorch and NumPy for the training set and other sets of random seeds for the testing set. The input-to-hidden weights (**W**^in^) and hidden recurrent weights (**W**^rec^) were initialized using a standard Kaiming initialization^95^ (He et al., 2016), where each entry was sampled from a Normal Distribution with mean 0 and variance 2/ *N*_RNN_.

In the MNIST task, we developed a spiking RNN-FNN architecture: one recurrent SRNN consisted of *N*_RNN_ = 200 hidden recurrent units (without E/I constraints) and an additional spiking FNN with *N*_FNN_ =50 non-recurrent inhibitory neurons (**Fig. 4f**). For simplicity, we assumed that the inhibitory units in FNN received input through fixed excitatory synapses and projected back to the hidden layer with fixed inhibitory synapses. In total, the RNN-FNN model had four types of synapses: plastic input weights **W**^in^, plastic hidden recurrent weights **W**^rec^, fixed hidden-to-inhibitory weights **W**^hid→inh^, and fixed inhibitory-to-hidden weights **W**^inh→hid^. Empirically fixed hyperparameters of **W**^hid→inh^ = 22.5, and **W**^inh→hid^ = −120 were used based on the previous setup^29^ (Diehl and Cook, 2018).

#### STDP plasticity

The synaptic weights of the spiking RNN or spiking RNN-FNN were updated either using standard BPTT or STDP algorithm. We adopted a trace-based rule for STDP, where each synapse was associated with two trace variables: a presynaptic trace *υ*_pre_ and a postsynaptic trace *υ*_post_. Both traces decayed exponentially in time and increased whenever a spike occurred, as described by the linear differential equations below:

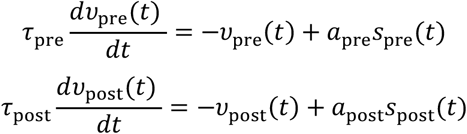

where *dt* = 0.001, *τ*_pre_ and *τ*_post_ are the time constants adjusted for specific tasks (see **Supplementary Table 2**); *a*_pre_ = *a*_post_ = 0.001 denote the scale factors that determine the contribution of each presynaptic or postsynaptic spike to the respective trace. Note that the time constant {*τ*_pre_, *τ*_post_} for the spiking event is much faster than the time constant *τ*_*m*_ for the neuronal membrane potential.

The synaptic weight update followed a Hebbian-like rule, where correlated the presynaptic and postsynaptic activity led to potentiation, whereas uncorrelated activity led to depression.

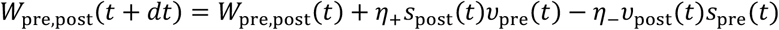

where *η*_+_ and *η*_−_ denote the learning rate parameters for potentiation and depression, respectively. In our experiments, we used *η*_+_ = 0.01, *η*_−_ = 0.0001 in the STDP update.

In the STDP application for the MNIST benchmark, we further implemented an adaptive strategy for the spiking threshold *V*_threshold_ as follows: *V*_threshold_(*t*) = *V*_threshold_baseline_ + *V*_theta_(*t*), where the adaptive component, *V*_theta_(*t*), evolved according to the following equation:

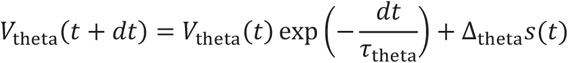

which was aimed to slightly increase the threshold with a small increment Δ_theta_ > 0 after a spike *s*(*t*)) was emitted.

#### Construction of synthetic training and testing samples

In the line detection task, we constructed the synthetic training and testing samples as follows. Each input consisted of a 10×10 grayscale image, and each image was first initialized with random pixel values and then multiplied by a noise level of 0.05. We generated two class labels: class 0 with a horizontal line present in rows 1-5 versus class 1 with a horizontal line present in rows 6-10 (**Fig. 5b**). The line had a thickness of 1 or 2 rows (chosen randomly with 50% probability). Finally, a random horizontal jitter from the set {−1, 0, 1} was selected, and the line was shifted left or right based on that amount. In total, 100 training images and 300 testing images were constructed.

#### Stimulus encoding

In the line detection task, we employed a Poisson encoder to convert the 10×10 grayscale images into spike trains for the SRNN input. At finite time steps *t* ∈ {1,2, …,50} and for every input pixel value *x*_*i*_ ∈ {0,1} (*i* = 1, …,100), we drew a uniform random variable *u* from the interval [0,1] with additionally imposed thresholding (such as a default setup: *u* < 0.1*x*_*i*_). We used a Poisson encoder to produce different spike trains on each forward pass or epoch. Since the sample size was rather small, we pre-encoded each image once into a deterministic spike train (over a simulation window of *T* = 50) and reused it across all training epochs. This procedure ensured us to obtain consistent spiking train inputs at each epoch and to avoid additional random fluctuations.

In the MNIST benchmark, the output layer corresponded to a 10-element vector of logits, each corresponding to a label. The input layer consisted of *N*_in_=28×28=784 input neurons (one for each image pixel). Each MNIST image pixel intensity was normalized between 0 and 1 that represented the firing probability of the input neuron, which was further encoded as a Poisson-distributed spike train for 200 timesteps. For each step, the *i*-th input neuron generated a spike 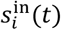 with a probability proportional to its normalized pixel intensity. At the end, the input consisted of a binary spike train matrix of size 784 × 200, where each entry indicates whether a spike occurred at a given time step. This encoder transformed static images into multiple spike trains that were subsequently fed into the SRNN. To impose a one-spike per time bin constraint, we assumed a refractory interval (5 ms) for each neuron after firing. During this period, the membrane potential was held constant to prevent spiking. Additionally, we imposed the one-spike constraint so that if multiple neurons simultaneously exceeded the spiking threshold within a single time bin, we randomly selected one neuron to emit a single spike.

#### Pattern classification

To enable our network to perform classification, we adopted a dynamic neuron-to-label assignment strategy^29^ (Diehl and Cook, 2015). Before training, we randomly assigned each hidden neuron to a label. Next, during each epoch and for each label *l*, we accumulated the spike counts *C*_*l,j*_ of the *j*-th hidden neuron whenever an input with label *l* was presented and tracked the total number of presentations for each label, denoted *N*_*l*_. At the end of each epoch, we computed the label assignment for the *j*-th neuron via the following equation

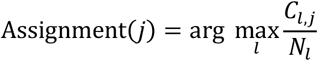

where both *C*_*l,j*_ and *N*_*l*_ were reset to zero at the start of each epoch. During testing, the final neuron-to-label assignments were fixed and used for classification. Specifically, for each test sample, we presented the input to the SRNN, collected spike counts from all hidden neurons, and then voted for the label with the highest total spike count among the neurons that were assigned to that label.

In the case of BPTT, a readout layer with hidden-to-output connections **W**^out^ was introduced to perform classification. Both the weights were initialized using a Kaiming uniform distribution. For each Poisson-spiking input sequence **x**(*t*), we generate an average output over the simulation window time *t* = 1, …,50, namely 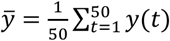. We then computed the cross-entropy loss ℒ between 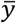 and the target output *y*_target_. We implemented the gradient descent using the “torch.autograd” function from the PyTorch-based Norse libraries (https://norse.ai/docs/), which automatically tracked the forward-backward pass in the calculations of all surrogate gradient 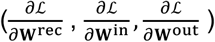 and applied the gradient chain rule through time. The surrogate gradient approximated the true gradient of spiking function with a smooth, differentiable function due to the non-differentiable nature of binary spike events. We used the Adam optimizer with a learning rate parameter of 0.1. In the line detection task, we varied the batch size (30, 50 and 100) to evaluate its impact on training convergence and test accuracy.

#### Convergence and performance assessment

We continuously monitored the network convergence at every step of training epoch. During inference, we froze all network weights and fed the test input to the network, determining the predicted class by applying the argmax function to the output layer activations. All models were evaluated based on the testing set. We defined a convergence threshold ϵ=0.005 (0.5%) such that the improvement was deemed negligible if the absolute difference in test accuracy between consecutive epochs was below ϵ. To guard against random minor fluctuations, we also introduced a patience hyperparameter 10 such that training would continue until 10 consecutive steps have met the convergence criterion. For the line detection task, we trained all models for 100 epochs. If the model had failed to converge at the 100^th^ epoch, then we treated the epoch 100 as the convergence point.

In the MNIST task, the criterion was similar except for using a convergence threshold of ϵ=0.01 (1%) and a patience hyperparameter of 5. To prevent runaway weight growth in STDP, we adopted a dynamic normalization strategy^29^ (Diehl and Cook, 2015) and normalized synaptic weights after each training iteration. An upper bound of synaptic weights was also imposed. For comparing multiplicative and additive couplings rather than achieving the optimal performance, we trained all models for a maximum of 30 epochs using a minibatch size of 256 (for BPTT) or 1 (for STDP). If the network failed to converge by then, we would treat the maximum epoch as the convergence point. We ran the experiments for 20 runs with different random seeds, where identical initial hyperparameter conditions were used for coupled and non-coupled models. Furthermore, in the STDP version of the MNIST task, we empirically imposed a maximum norm for all weights to assure the numerical stability.

#### Implementation

The SRNN training was implemented on a high-performance computing (HPC) cluster that used the Slurm Workload Manager (version 24.11). We created Slurm job files that specified the hidden layer size, hyperparameters, and a random seed. In the line detection task, we iterated over random seeds {0, 1, …,99} for each combination of hyperparameters, producing 100 Monte Carlo runs per setting. Each Slurm job file loaded the pre-encoded training and testing data, set the specified seed within both NumPy and PyTorch prior to model initialization, and then ran for a fixed number of epochs of training. After each epoch, the job computed testing accuracy for the subsequent analysis.

### Biologically constrained PFC-MD network in context-switching tasks

To model the recurrent PFC network, we implemented an excitatory-inhibitory (E/I) RNN while imposing Dale’s law constraint^13^ (Zhang et al., 2022). Briefly, the PFC consists of both excitatory and inhibitory units. According to Dale’s principle, the ratio of Exc and Inh units is 4:1. In our computer simulations, the total number of PFC units was 256 (*N*_PFC_=256), and we set *N*_exc_=205, and *N*_inh_=51. Mathematically, the synaptic connectivity constraint can be expressed by the following equation^13^ (Zhang et al., 2022)

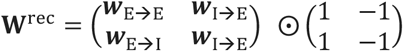

where the Hadamard product ⨀ imposes the structural connectivity between excitatory and inhibitory cell types.

To introduce PFC-MD couplings in the PFC-MD network, we first considered a standard model where the PFC receives an additional additive input from the MD output **r**^MD^ = ***ϕ***(**x**^MD^) scaled by the thalamocortical synapses **W**^MD→PFC^, and the MD receives the direct input from the PFC^14^ (Zhang et al., 2025)

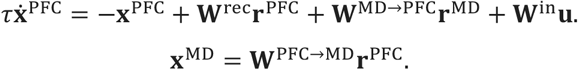

where **r**^PFC^ = ***ϕ***(**x**^PFC^) denotes the respective non-negative firing rate vector for the PFC, ***ϕ*** denotes a ReLu function and **u** denotes the external input. Let **h** = **W**^MD→PFC^**r**^MD^ = **W**^MD→PFC^***ϕ***(**W**^PFC→MD^**r**^PFC^) represents a copy of PFC activity; because the thalamocortical and corticothalamic synapses {**W**^MD→PFC^, **W**^PFC→MD^} are excitatory and ***ϕ*** is a ReLu function, we can further simply the term **h** = **W**^MD→PFC^**W**^PFC→MD^**r**^PFC^ and rewrite the PFC dynamics as follows

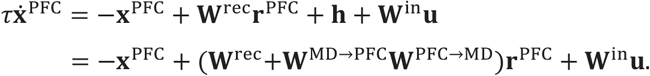

Let **L** = **hh**^*T*^denote an outer product of the efference copy that carries the sufficient statistics of **h**; we introduced the synaptic-neuronal coupling via PFC-MD multiplicative coupling governed by the following dynamic equation:

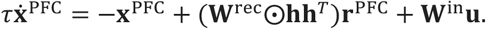

where ⨀ denotes the matrix Hadamard product between the recurrent weight matrix **W**^rec^ and cross-correlation matrix **hh**^*T*^ of the same dimensionality.

Finally, the network output is based on a linear readout of PFC population activity: **y** = **W**^out^**r**^PFC^ (for continuous output) or **y** = *σ*(**W**^out^**r**^PFC^) (softmax function for categorical output).

We trained the PFC-MD model to perform a context-dependent two alternative force choice (2AFC) task with auditory cues (**Fig. 5a**). During the 400-ms cueing period, auditory low-pass (LP, −1) or high-pass (HP, +1) pulse were presented to the network. Different cues match different rules (“attend-to-audition” vs. “attend-to-vision”). During 100-ms delay period, rule information needs to be preserved. During 100-ms target period with “divided attention”, the auditory and visual stimuli (independent from the cueing period) are simultaneously presented, and each attended target is associated with a unique outcome out of two choices (**Fig. 5a**). We further considered a switching context task in a block-by-block fashion (*Context 1*→*Context 2*→ *Context 1’*). In *Context 2*, the rule was reversed from *Context 1*, and in *Context 1’*, the rule was the same as *Context 1*. We first trained the PFC-MD network based on supervised learning and refer the reader to REF^14^ for technical details. As a generalization, the probabilistic reward was used in the reward period, producing an outcome uncertainty condition (**Fig. 5c**). We then trained the PFC-MD network based on Actor-Critic RL. While the technical details are similar, we used a different hyperparameter setup.

We also trained the PFC-MD model to perform a human probabilistic Go/NoGo reversal learning task with a minor modification from REF^31^ (Wang et al., 2023). In each block, two tactile patterns were use consistently throughout the experiments (one designated as the ‘Go’ pattern and one as the ‘NoGo’ pattern). Each trial began with a fixation cross, followed by a 200-ms tactile stimulation period presenting one of these two patterns to the participant’s right index finger. After the tactile cue, the fixation cross turned green, signaling the participant to choose a response (either ‘Go’ by pressing a button with their left index finger, or ‘NoGo’ by withholding a press). Following the response, after a 100 ms interval, the outcome was presented visually for 100 ms, indicating whether the choice was rewarded or not. The reward contingencies were probabilistic: 70% of trials with the first tactile pattern were rewarded if the agent chose ‘Go’, and 70% of trials with the second tactile pattern were rewarded if participants chose ‘NoGo’. By trial and error, the participant had to learn which of the two available options (‘Go’ and ‘NoGo’) had the higher reward probability for each of the two tactile patterns. Importantly, in each individual block, the association between tactile stimuli and responses was switched at a random trial (reversal). After rule switching, the participant needs to reverse their choice behavior to maximize reward.

### PPC-Pulvinar-PFC network in working memory and attention tasks

In a similar manner to E/I-PFC-MD modeling, we modeled the PPC and PFC as two E/I-RNNs, and the pulvinar as one excitatory neuron-only FNN. We assumed that the thalamo-cortical and cortico-thalamic connectivity is bidirectional and asymmetric. In a general setting, we assumed that the pulvinar receives cortical inputs from both PPC and PFC, and the pulvinar output multiplicatively modulates the cortico-cortical (PPC→PFC and PFC→PPC) connectivity as follows:

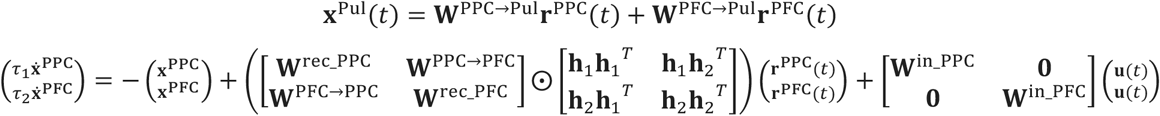

where **h**_1_ = **W**^Pul→PPC^**r**^Pul^, **h**_2_ = **W**^Pul→PFC^**r**^Pul^ and **r**^Pul^ = ***ϕ***(**x**^Pul^), where ***ϕ*** denotes the activation function.

To focus our study on thalamic gating of cortico-cortical connectivity, we considered a simplified version of the multiplicative gating of the full model:

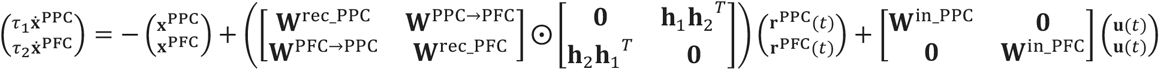

where **h**_1_**h**_2_^*T*^ and **h**_2_**h**_1_^*T*^ denote the thalamic gating. When *N*_PPC_= *N*_PFC_, we further used the inner product value (**h**_1_^*T*^**h**_2_ = **h**_2_^*T*^**h**_1_) to quantify the strength of thalamic gating.

For the network size, we set *N*_PPC_= *N*_PFC_ =200 (with E/I ratio of 4:1: *N*_exc_=160, and *N*_inh_=40), and *N*_Pul_=64. In the context of WM, PFC neurons tend to have a faster time constant and stronger recurrent synapses than PPC neurons^97^ (Murray et al., 2017), we assumed that the PFC has a faster time constant (*τ*_2_ = 40 ms) than the PPC (*τ*_1_ = 80 ms). Furthermore, we assumed that the PPC and PPC receive the sensory input with different strengths (i.e., **W**^in_PPC^ > **W**^in_PFC^). We also assumed the long-range cortico-cortical connectivity were sparse, with excitatory-to-excitatory connectivity only. Additionally, we assumed sparse thalamocortical and corticothalamic connections, with stronger connectivity (sparsity level 0.3) in the pulvinar-PPC pathway than in the pulvinar-PFC pathway (sparsity level 0.7). We further assumed that the output was generated by a linear readout of the PFC activity:

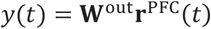

As a control, we also considered a PPC-PFC model without thalamic gating as well as a PPC-Pulvinar-PFC with additive thalamic gating. The simplified cortical dynamics were described as follows:

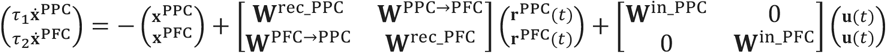

and the output equation is identical. Similarly, an additive cortico-thalamic-cortical model has the following RNN dynamics:

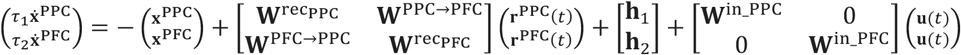

We trained the multiplicative gating and control models using the BPTT algorithm (based on similar setting as the previously described PFC-MD model) to perform both WM and attention tasks (**Fig. 6b**), which were modified from^35^ (Panichello and Buschman, 2021).

Briefly, the WM task (aka. “Retrospective trials”) starts with a fixation period of 80 ms, followed by a 200-ms stimulus period in which either 1 or 2 squares is presented at two possible locations. The color of each sample was drawn randomly from 8 evenly spaced points along a colored circle. Colors are independent across locations. Following a pre-cue delay of 200 ms, spatial cues appear indicating which sample (upper or lower) should be reported. Two sets of spatial cues are used: 45° clockwise vs. anticlockwise lines or triangle vs. circle symbols. After another 200-ms post-cue delay, the target appears for 80 ms with cued color, and the goal is to choose the colored region that corresponds to the color of the cued memory. In the attention task (aka. “Prospective trials”), the basic task structure is similar except that the spatial cues are presented 200 ms before the first delay period. Note that the functions of the first and second delay periods are swapped between the retrospective and prospective trials.

In population decoding analysis, we used a nonlinear support vector machine (SVM) classifier with a first-order polynomial kernel and default hyperparameters to classify upper vs. lower classes. The input features corresponded to the single-trial firing rate of individual units from the PPC or PFC. We used 512 simulated trials for training and 512 simulated trials for testing. We slid the moving window *dt*=20 ms and repeated the decoding procedure from time −0.2 s to 0.2 s (time 0 denotes the cue onset). The decoding accuracy was assessed by 5-fold cross-validation from 20 Monte Carlo simulations. We reported the mean ± s.e.m. decoding accuracy and compared our model with experimental data from REF^35^.

To measure the angle between two planes, we adopted the methods in REF^35^ based on the principal component analysis (PCA). We constructed the first three dominant principal subspace whose axes correspond to the first three eigenvectors of the covariance matrix and then projected the population vector for a given condition into this reduced dimensional subspace. We measured the cosine of the angle, cos(*Θ*), between the two planes spanned by the vectors that defined the “up” and “down” items, respectively: 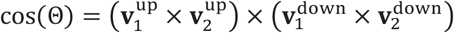. See REF^35^ for technical details of hyperplane construction and angle computation. To assess how cos(*Θ*) changed around cue onset, we used a logistic regression model of the form: 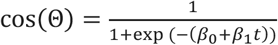, where *t* denotes the time relative to the cue onset. We also compared our model predictions with experimental data from REF^35^.

Furthermore, we computed the PCA subspaces separately for the PPC and PFC populations derived from the trained PPC-Pulvinar-PFC model. We adopted the same method described in REF^14^ and computed the subspace angles (“principal angle”). Briefly, let **C**_**1**_ and **C**_**2**_ be two *d*-dimensional PC projection matrices derived from two respective higher-dimensional neural population data (for instance, from separate PPC and PFC populations measured concurrently, or from the same PFC populations measured independently in two different tasks). The two *d*-dimensional subspaces have a total of *d* angles, denoted by {*θ*_1_, …, *θ*_*d*_}. The angle *θ*_1_ is defined as the largest angle determined by two dominant principal vectors (say 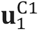 and 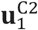) within the range of **C**_1_ and **C**_2_, respectively. Similarly, the angle *θ* is defined as the largest angle determined by vectors 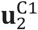 and 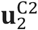 constrained to be orthogonal to 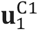 and 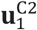, respectively. This definition is continued iteratively until *θ*_*d*_. In practice, these angles can be obtained as the cosine of the singular values of the inner product of two matrices: 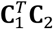. In our analysis, we computed the first and second largest principal angles (*θ*_1_ and *θ*_2_). Geometrically, an angle of 0 degree indicates that two subspaces are parallel, whereas an angle of 90 degree indicates that two subspaces are orthogonal. If *θ*_1_ = *θ*_2_, it often implies that the joint neural population activity primarily encodes a single task variable. On the other hand, if the gap between *θ*_1_ and *θ*_2_ is large, it implies that the joint neural population activity encodes two or more independent task variables.

### EC-DG-CA3-CA1 network in a visuospatial navigation task

In our computer simulations, training and testing were conducted by generating trajectories within a two-dimensional circular arena (radius of 1.3 m). To mimic an animal’s movement, we developed a framework for random walk generation, based on stochastic speed and head direction within a 2D space. Each trajectory spanned 10 steps. The starting position was sampled from a uniform distribution within the circular enclosure, with values ranging from −radius/2 to radius/2. The subsequent positions in the random walk were computed by integrating the starting point with generated Cartesian velocities. At each step, velocities were determined by independently sampling the speed and head direction. The simulation used an integration time step of 0.02 s. The initial run speed parameter was drawn from a Gaussian distribution with a mean of 0.08 m/s and a standard deviation of 0.01 m/s. The head direction was sampled from another Gaussian distribution, with zero mean and a standard deviation of 11.5 rad.

Based on the head direction, we defined a viewing area represented by an 8×8-pixel image (i.e., *N*_in_=64), corresponding to the raw visual input (e.g., pixel luminance). To reduce the data’s dimensionality, we applied principal component analysis (PCA) and projected the vectorized image onto the principal component (PC) subspace that captured ~90% of the variance in the visual stimuli. In sum, the input dimensionality was 2+2=4.

Following an established method^98^ (Banino et al., 2018), the place receptive field was modeled using a 2D isotropic Gaussian function to encode a 2D position **z**, defined as

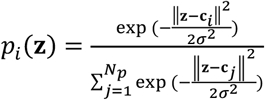

where *i, j* = 1,2, …, *N*_*p*_ and *N*_*p*_ = 512 representing the total number of place cells in the output. The term *c*_*i*_ specifies the center of two-dimensional (2D) place receptive field of the *i*-th place cell, and the parameter *σ* = 0.12 m determines the spatial width of each place cell.

The constructed EC-hippocampus circuit was derived based on the experimental knowledge of genetically identified hippocampus circuit projections (**Fig. 7a**). We assumed that the EC exhibits recurrent connection, the velocity and visual signal input from the EC enters the hippocampus and projects to the DG. The DG acts as a gatekeeper and is thought to perform pattern separation. Granule cells in the DG project to the CA3 region, which has a recurrent structure. The CA1 pyramidal neurons receive projections of CA3 neurons in a topographically organized manner, and then project back to EC, forming a loop. EC can also bypass DG and CA3 and send direct input to CA1. The number of neurons in each population is *N*_EC_ = *N*_DG_ = *N*_CA3_ = *N*_CA1_ = 512. Each brain area consists of both excitatory and inhibitory neurons. However, for modeling simplicity, we did not impose the E/I constraints onto each module.

In the case of additive coupling, we assumed that the RNN dynamics of EC was described by the following equation:

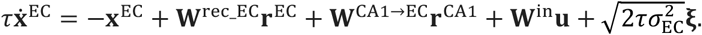

where *τ* = 100 ms represents the time constant, ***ξ*** denotes *N*_EC_-dimensional Gaussian noise, independently sampled from a standard normal distribution, and *σ*_EC_ = 0.05 defines the standard deviation; **W**^rec_EC^ is an *N*_EC_×*N*_EC_ matrix for recurrent weights within EC, **W**^CA1→EC^ is an *N*_CA1_×*N*_EC_ matrix representing CA1-to-EC projection weights, and **W**^in^ is an *N*_EC_ ×*N*_in_ matrix for the input connection (*N*_in_ =4). The neuronal firing rates **r**^EC^ were determined using a ReLU activation function: **r** = ReLU(**x**).

The FNN dynamics of DG was described by

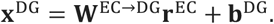

where **W**^EC→DG^ is an *N*_EC_×*N*_DG_ matrix representing projection weights from EC to DG, and **b**^DG^ is a bias vector.

The recurrent dynamics of CA3 was described by

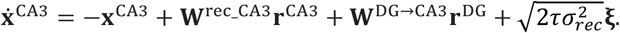

where **W**^rec_CA3^ is an *N*_CA3_×*N*_CA3_ matrix defining recurrent connection weights within CA3, **W**^DG→CA3^ is an *N*_DG_×*N*_CA3_ matrix representing the feedforward connection from DG to CA3; **r**^CA3^ and **r**^DG^ denote the respective neuronal firing rate vectors in CA3 and DG.

Finally, the FNN dynamics of the CA1 region were described by:

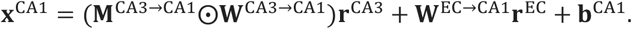

where **W**^CA3→CA1^ is an *N*_CA3_×*N*_CA1_ matrix representing projection weights from CA3 to CA1, and **b**^CA1^ is the bias vector. The masking matrix **M**^CA3→CA1^ is a block matrix used to define the feedforward structural connectivity, with ⨀ denoting the elementwise (Hadamard) product. Specifically, the block matrix **M**^CA3→CA1^ is defined as

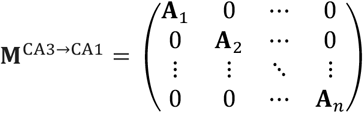

where each block **A**_1,_ **A**_2,_…, **A**_*n*_ is a 16×16 matrix filled with one and *n* = 32.

In the network output, we pooled the neuronal firing rates from all three hippocampal subfields (DG, CA3, and CA1) to encode the place cell activity at each time point. Thus, the predicted place cell activity was expressed as:

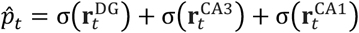

where *σ* denotes the vector-valued softmax function operated on the DG, CA3 and CA1 firing rate vectors. The weights **W**^EC→DG^, **W**^EC→CA1^, **W**^DG→CA3^ were initialized with a Gaussian distribution of mean 0 and standard deviation 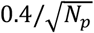 (where *N*_*p*_ denotes the number of neurons in the target area) whereas the remaining connection weights in the network were initialized to zero.

In the case of multiplicative coupling, we assumed that the EC dynamics were modulated by CA1 activity through the CA1→EC feedback. Let **h** = **W**^CA1→EC^**r**^CA1^, we imposed a multiplicative gating **W**^rec_EC^ via **hh**^*T*^. Specifically, the recurrent EC dynamics can be rewritten as

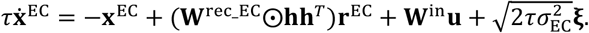

Compared to additive gating, the multiplicative gating model has the same degrees of freedom in terms of unknown parameters and identical DG, CA3, and CA1 dynamics.

During training, we used a loss function consisting of two terms: a cross-entropy term to compare the predicted and true place cell activities, and an *L*_2_-regularization term to penalize large recurrent weights. For a simulated trajectory of sequence length *T*, the loss is defined as:

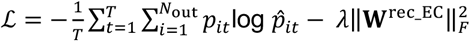

where *p*_*it*_ represents the true activity of the *i*-th place cell at time *t*; ‖·‖_*F*_ denotes the Frobenius norm of the matrix; 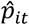 denotes the model-predicted activity; *λ* is the regularization coefficient which was set to 0.0001. In all navigation conditions, we set time constant *τ* to 30 ms, temporal bin size *dt* = 20 ms and training sequence length *T* = 10.

In replay analysis, we modified the network output and used CA1 only to encode the place cell activity at each time point. Thus, the predicted place cell activity was rewritten as 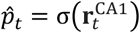. During the offline state, the speed of replay was typically much faster than the run speed. We reduced the ratio of *τ*/*dt* and set *τ* to 25 ms or 20 ms. We randomly selected an initial position within the environment and fed the network with small random noise. The network generated internal neural sequences for a length typically longer than 10. From each reactivated neural sequence, a unique spatial trajectory can be decoded or reconstructed based on their place fields. Furthermore, given the same initial position, different random noise or random seeds could produce different replay trajectories. We repeated the procedure for n=250 random positions, each with 5 random seeds uniformly sampled from 0 to 99. Finally, to estimate the average speed of each replay trajectory, we computed the sum of virtually traveled distance and normalized it by the total time (i.e., *dt* multiplied with the sequence length).

### Dimensionality reduction and neural trajectory visualization

To extract low-dimensional representation of population activity from the RNN-FNN network, we employed PCA to visualize neural trajectories during the task period (**Fig. 2**). Briefly, we grouped neural population activity into a matrix **X** ∈ ℝ^*N*×*T*^, where *N* denotes the number of units and *T* denotes the number of time bins. We performed PCA on temporally binned data matrix **X** during specific task periods and further extracted two dominant principal subspaces associated with the two or three largest eigenvalues. We projected the neural activity onto the dominant PC subspaces and visualized the 2D or 3D neural trajectory. For multiple trials, we augmented the trials and then computed trial-averaged neural trajectories. Furthermore, we computed the “neural velocity” norm based on the low-dimensional neural trajectory.

In the integration benchmark (**Extended Data Fig. 2**), we conducted dimensionality analysis (PCA followed by nonlinear embedding) for the 3-by-*T* (*T*=50 or 100) dimensional neural trajectories for two classes of trained RNN-FNN models: one with multiplicative and the other with additive coupling. The neural trajectories were generated by initial conditions of **x**(0) = (0,0,0) and fed with the identical input sequence **u**(*t*). We first conducted linear PCA for the 150-dimensional neural trajectories and used the first top 10 PC components for two-dimensional t-SNE embedding, with a default perplexity parameter of 10.

### Animal behavior and neurophysiology

Vgat*-*Cre mice (016962) and C57BL/6J mice (000664) were obtained from Jackson Laboratory. All mice tested were between 2-12 months of age and housed on a 12-h light/dark cycle. Male mice were used for behavioral testing to reduce potential confounds from placing mice both genders sequentially in the same behavioral testing environment, while mice of both sexes were used for all neurophysiological experiments. All experiments were carried out under protocols approved by the Committee on Animal Care at Massachusetts Institute of Technology (MIT) and conformed to NIH guidelines.

Mice were trained on the cross-modal decision-making task as previously described^27,30^ (Schmitt et al. 2017; Rikhye et al., 2018). The initial duration of delay period was 0.5-0.9 s by default. Here we further extended the delay period up to 2.1-2.5 s. This was achieved over several weeks by gradually increasing the time the mice were required to hold their nose into the initiation port in 50-100 ms intervals from session to session. The time was only increased if mice were able to initiate at least 50 trials in the previous session. At the behavioral level, mice were able to achieve comparable task accuracy.

Electrophysiological data was acquired and analyzed as previously described^27^ (Schmitt et al. 2017). Stereotrode recordings from the prelimbic part of the PFC were acquired using a Neuralynx multiplexing digital recording system (Neuralynx, Bozeman MT). Electrode signals were amplified, filtered between 0.1 Hz and 9 kHz and digitized at 30 kHz. Following acquisition, spike sorting was performed offline based on relative spike amplitude and energy within electrode pairs using the MClust toolbox. Spike trains of each unit were aligned to behavior trials using TTL time stamps, separated by trial type and plotted as raster and peri-stimulus time histograms (PSTHs). During the delay period, rule-tuned PFC units showed timing-specific peak firing pattens. The tuning was specific to exclusive to one rule (“Attend-to-vision” or “Attend-to-audition”) but not both^27,30^ (Schmitt et al. 2017; Rikhye et al., 2018).

### Statistics and reproducibility

Computer simulations on benchmarks were repeated with Monte Carlo runs with different random seeds (n=20-100). Computer modeling on neural circuits was conducted by training multiple (n=10-20) independent RNN-FNN models, with each based on the same network size and architecture but different initial conditions. The researchers were not blinded to allocation during experiments and outcome assessment. Statistics testing was conducted based on nonparametric unpaired random-sum and paired signed-rank tests, or paired t-test with FDR correction.

## Data availability

Benchmark datasets are available in the public. Computer simulations are available along with the source code. Source data are provided with this paper.

## Code availability

PyTorch is freely available at https://pytorch.org. Norse library is an open source PyTorch library available at https://github.com/norse. All custom software for benchmark and modeling experiments that supports the analysis and plots are collectively available at https://github.com/Xh-Zhang1/multiplicative and https://github.com/KhairKhair/spikingNN-multiplicative-coupling-

## Acknowledgements

We thank György Buzsaki for some valuable feedback. The work was supported by grants DA056394 (Z.S.C.), MH132642 (Z.S.C., M.M.H.) and MH139352 (Z.S.C., M.M.H.) from the US National Institutes of Health. A preliminary version of this work was presented in the annual COSYNE meeting, March 29, 2025.

## Author contributions

Z.S.C. conceived and supervised experiments, analyzed and interpreted the data, and wrote the paper. X.Z. and M.A. developed the computational models, performed experiments, analyzed and interpreted the data. W.X. and Z.S.C. derived the mathematical proof. R.W. and M.M.H. provided data from the mouse experiments. Z.S.C. wrote the manuscript, with additional inputs from all other authors. All authors read, edited and approved the manuscript. Z.S.C. acquired funding.

## Supplementary Information

Supplementary Notes 1-7 and Tables 1-3.

Supplementary Video: Comparison of the evolution of two-dimensional neural trajectories between additive and multiplicative couplings during learning. Screenshots were updated every 10 epochs.

---

**Extended Data Fig. 1.**
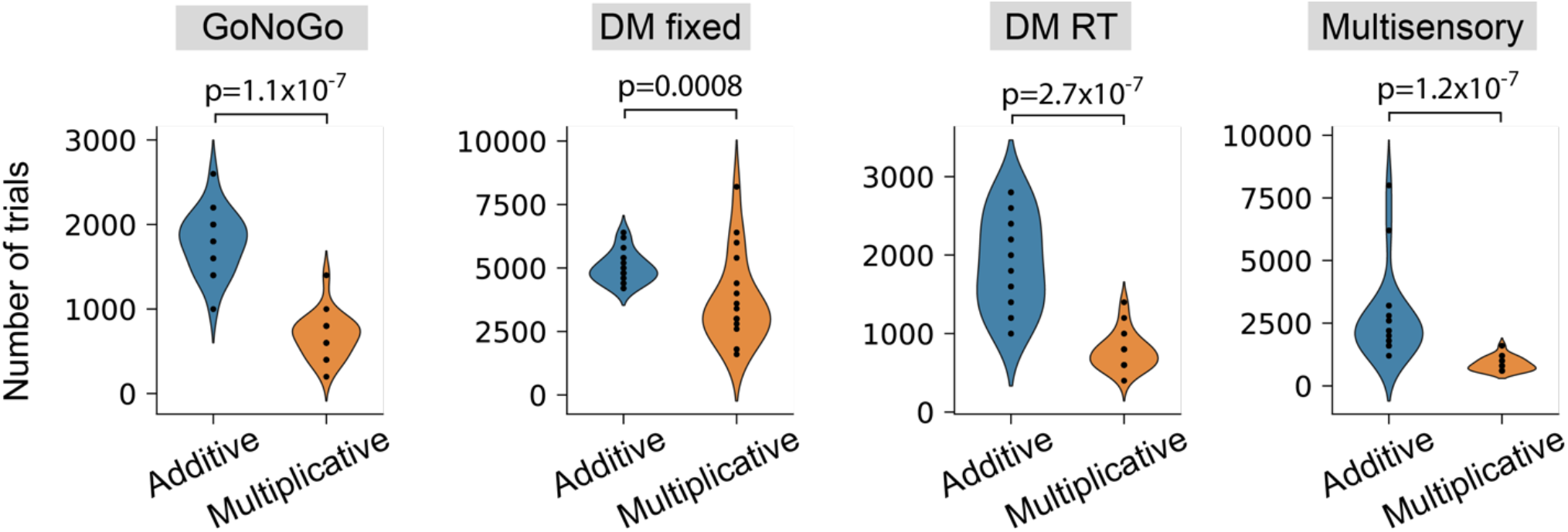
Additional results on the WM and DM benchmarks. Violin plot comparisons of convergence speed on four benchmarks with a minibatch size of 2. All p-values were computed based on the rank-sum test (n=20).

**Extended Data Fig. 2.**
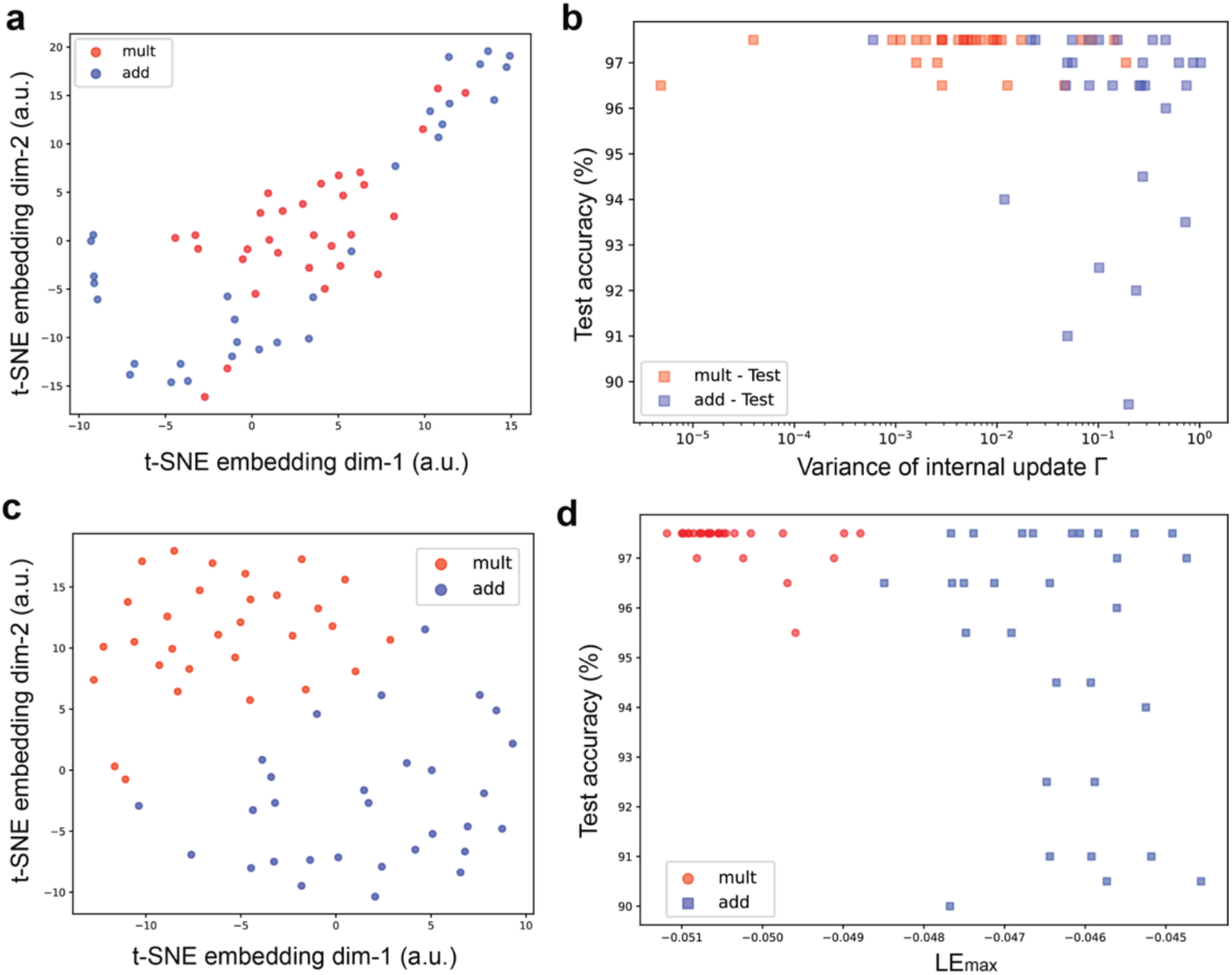
Additional test results of 3×3 RNN-FNN on the Integration benchmark. (**a**) Dimensionality reduction (PCA followed by t-SNE on top 10 PCs) analysis of neural trajectories during noisy-delay input period (*T* = 50). The two-dimensional embedding space (arbitrary unit) revealed distinct clusters between multiplicative (red) and additive (blue) gating RNN-FNN models (*τ* = 2, *dt* = 0.1, *n* = 30 for each class). Each model was initialized with the same initial state condition **x**^RNN^(0) and fed into an identical input sequence with length *T* = 100. **(b)** The scatter plot of testing accuracy of all models versus their respective variance of the internal update term **Γ** during the delay period (*T*_delay_ = 50; x-axis displayed in the log domain only for the better separation purpose) showed a negative correlation (Spearman’s rank correlation *ρ* = −0.455, *p*=0.000259, *n*=60). Each dot denotes a trained RNN-FNN model (*n* = 30 for each class). (**c**) Upon dimensionality reduction on the 3×3×100 Jacobian matrix traces, two-embedding space showed two separated classes. (**d**) The scatter plot of testing accuracy of all models versus their respective largest Lyapunov exponent estimates (Spearman’s rank correlation *ρ* = −0.578, *p*=1.3×10^−6^, *n*=60).

**Extended Data Fig. 3.**
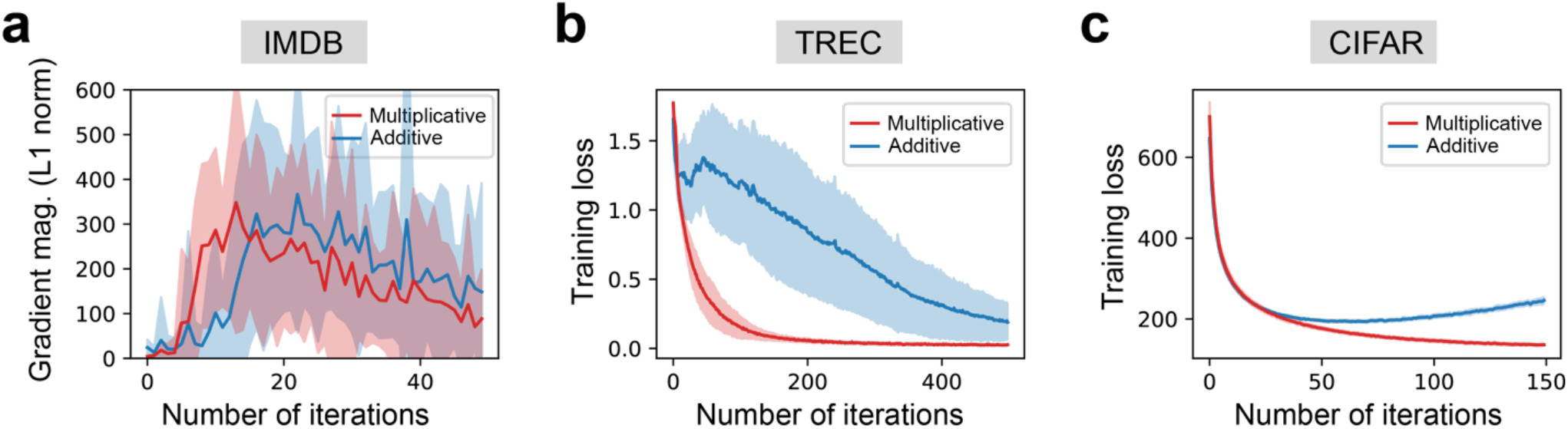
Additional results on the IMDB, TREC, and CIFAR benchmarks. (**a**) Comparison of RNNs’ gradient flow magnitude (L_1_ norm) in the IMDB benchmark explained why multiplicative coupling converged faster than the vanilla RNN. Shaded area denotes s.e.m. (n=50). (**b,c**) Training loss curves for the TREC (**b**) and CIFAR (**c**) tasks. Error bar denotes s.e.m. (n=20).

**Extended Data Fig. 4.**
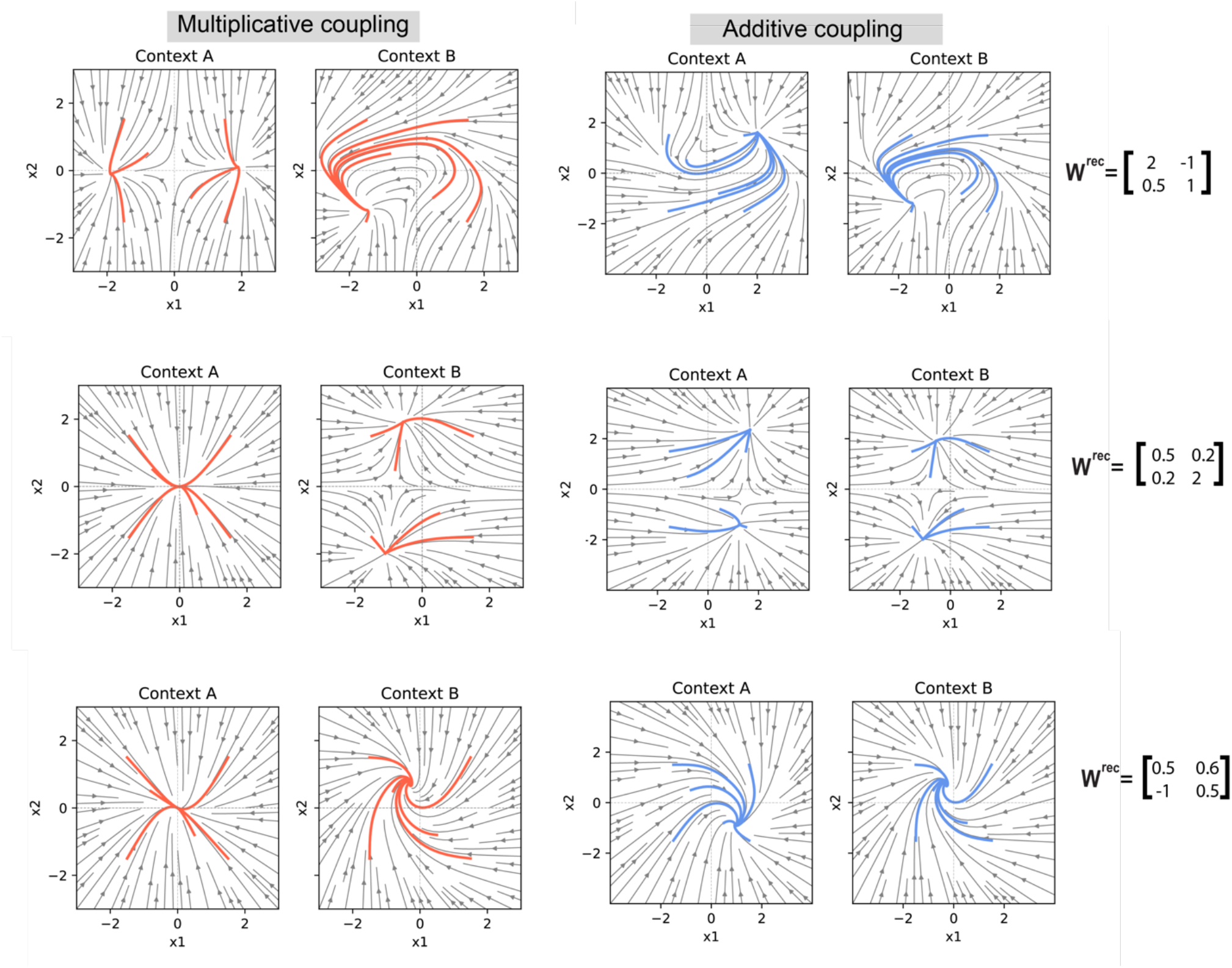
Illustrations and comparisons of phase portraits beween multiplicative and additive couplings in a 2×2 RNN-FNN architecture. In each example, two RNN-FNN networks (one with additive coupling and the other with multiplicative coupling) received the same yet context-dependent feedback vector **h** (i.e., **h**_context A_ ≠ **h**_context B_), but only multiplicative coupling showed distinct attractor dynamics in the 2D latent neural space **x**^RNN^ =[x_1_, x_2_]^*T*^ between two different contexts (Context A vs. Context B), thereby accommodating context-dependent gating. In contrast, additive RNN-FNN coupling showed rotational dynamics during context switching, which has also been previously demonstrated in a high-dimensional PFC-MD network (see Fig. 8g of REF^14^).

**Extended Data Fig. 5.**
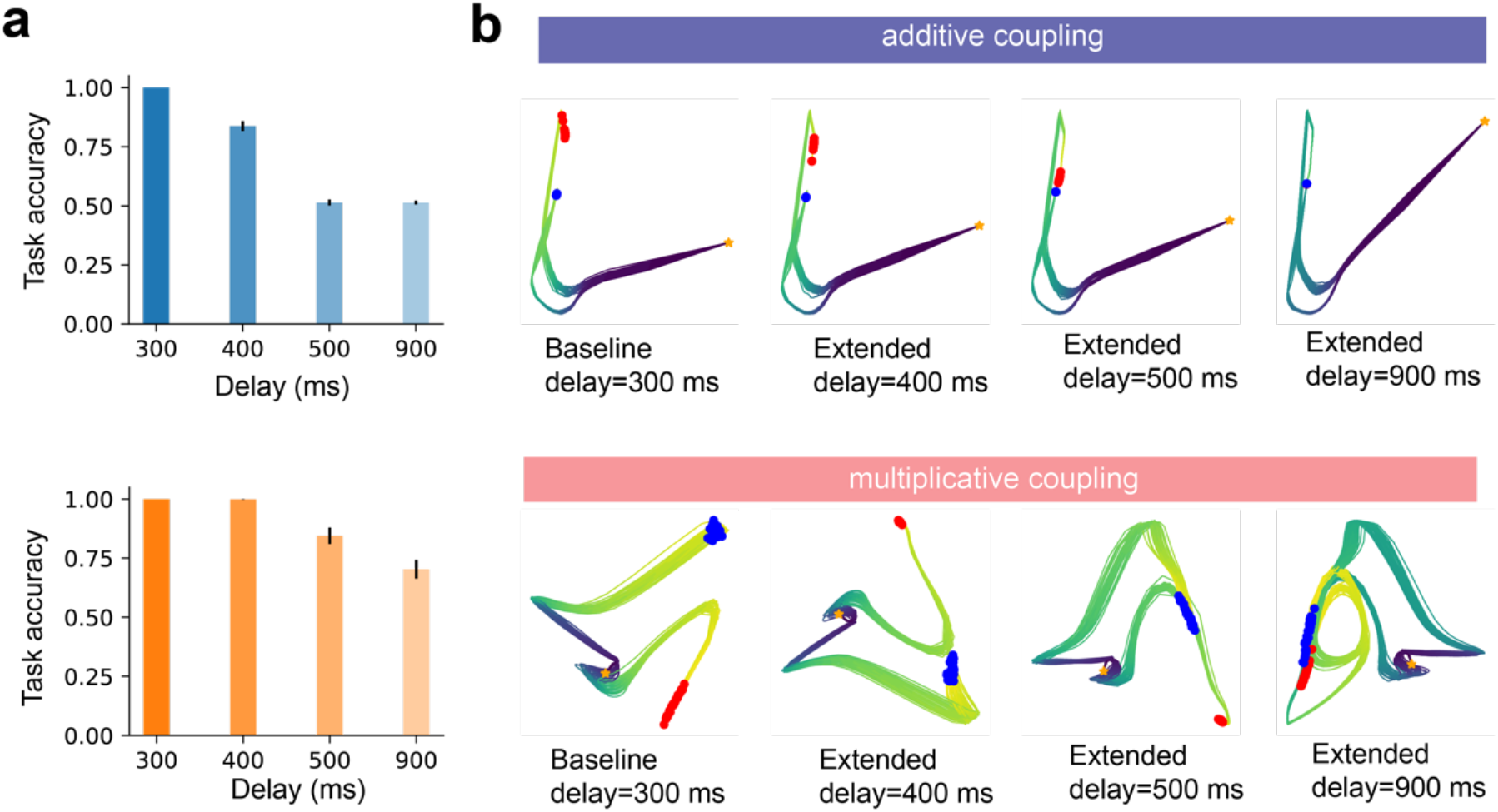
Testing the robustness of RNN-FNN models with respect to delay duration in the “DM fixed” benchmark. **(a)** Comparison of models’ test accuracy with various WM delay duration. Multiplicative couplings showed better model robustness with respect to extended delay, whereas additive couplings quickly dropped the performance to the chance level (50%). Error bar denotes s.e.m. (n=50). (**b**) Comparison of neural trajectories between additive and multiplicative couplings. Each line represents a trial. Blue and red dots represent end points of the trials from two distinct classes. Star symbol represents the trial onset. As the extended delay duration increased, the separation between blue and red dots became more difficult. In additive coupling, the shapes of neural trajectories were similar across four delay conditions, whereas in multiplicative coupling, the shapes of neural trajectories were more dynamic.

**Extended Data Fig. 6.**
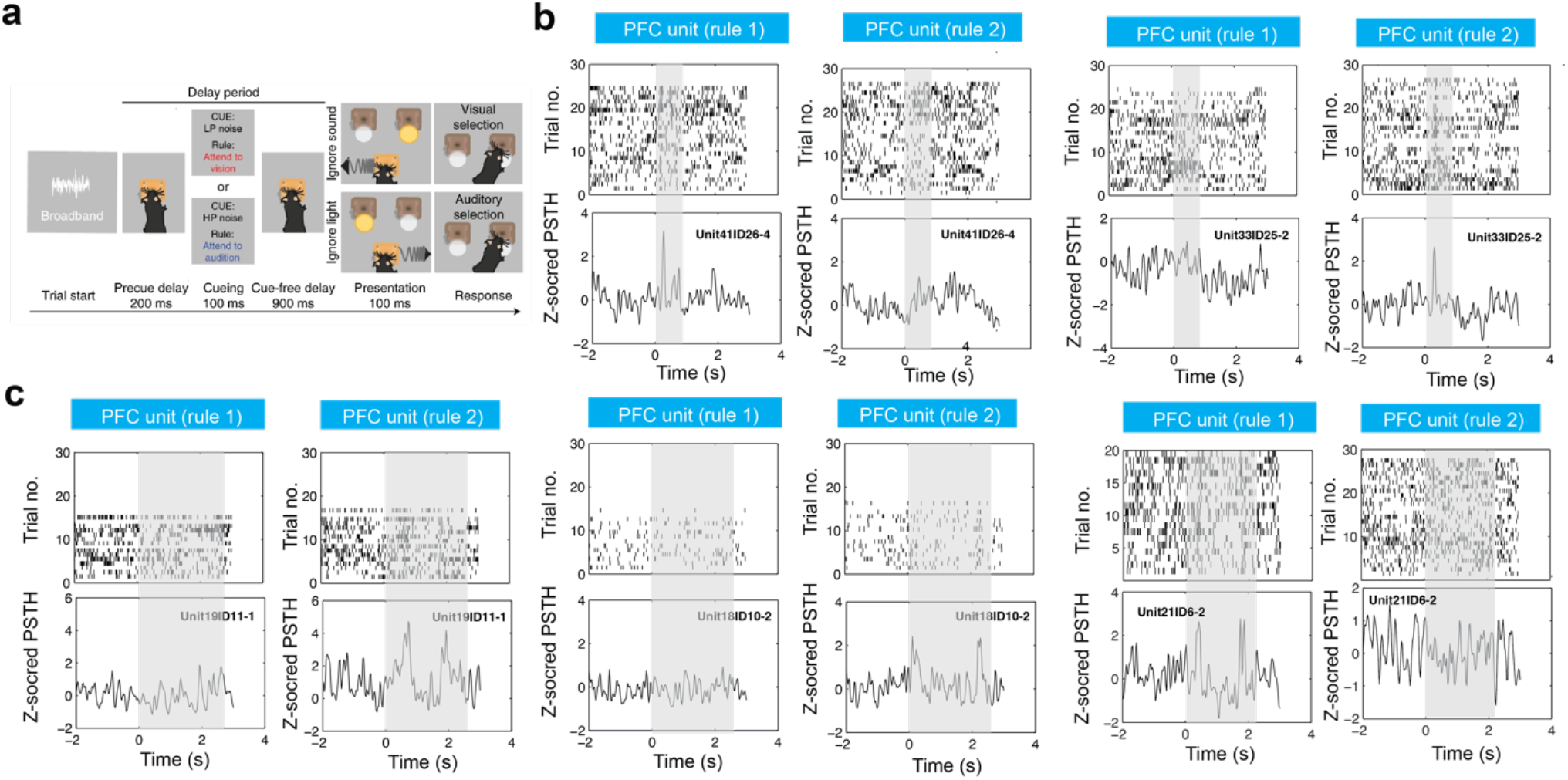
Schematic of the cross-modal decision-making task (a) and additional representative spike raster and PSTH examples of mouse PFC units (b,c). Panel **a** was adapted from REF^27^ with permission (© Springer Nature). Time 0 denotes the delay onset. Shaded areas in all panels denote the stimulus-free task delay periods. (**b**) During the shorter delay (shaded area [0, 0.9] s), one single peak was observed. (**c**) During extended delay period (shaded area [0 2.5] or [0, 2.1] s), two peaks emerged, resembling quasi-periodicity encoding during WM.

**Extended Data Fig. 7.**
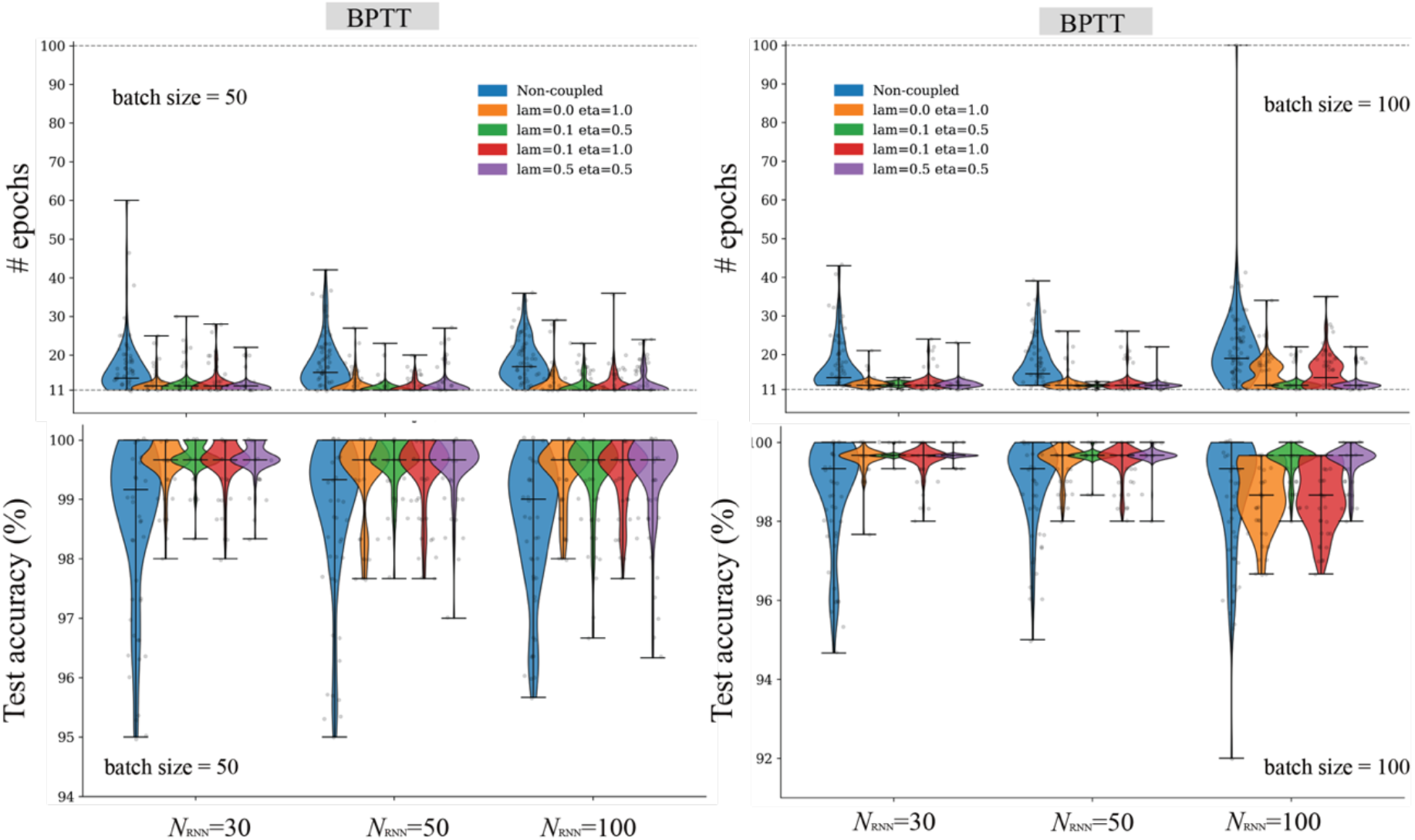
Additional training and testing results of SRNNs on the line detection benchmark. The convergence speed and test accuracy were shown with respect to various hyperparameters in associative memory (*λ, η*), batch size, and the network size *N*_RNN_.

**Extended Data Fig. 8.**
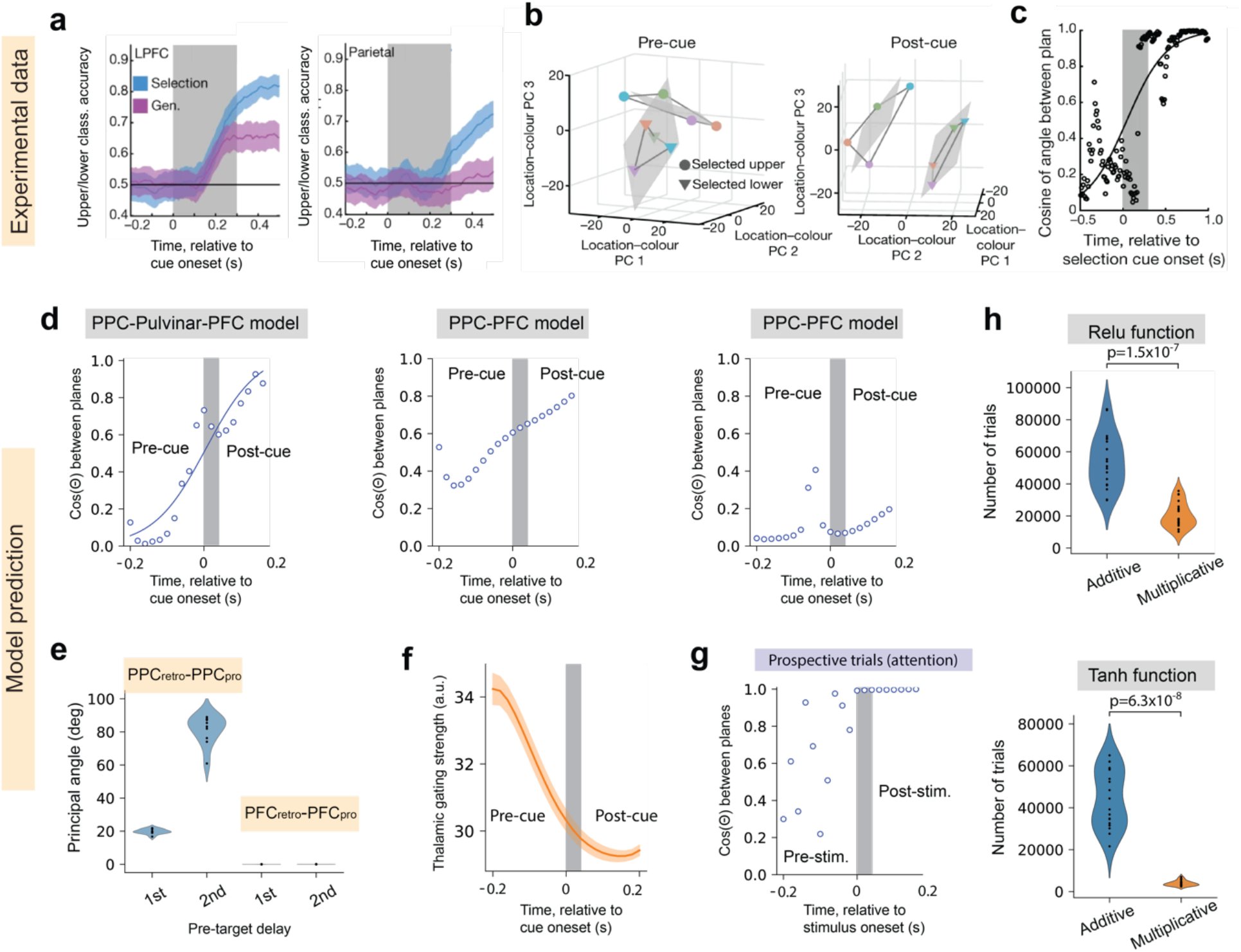
Additional results of multiplicative gating of PPC-PFC connectivity. (**a-c**) Experimental data matched our model prediction results (**Fig. 6d-f**), all panels were modified from REF^35^ with permission (© Springer Nature). (**d**) *Left panel:* Same as **Fig. 6f**, except that results were generated from another trained PPC-Pulvinar-PFC model. *Middle and right panels:* Two representative results generated from two independently trained PPC-PFC models. (**e**) The first and second principal angles of PFC_retro_-PFC_pro_ and PPC_retro_-PPC_pro_ subspaces computed from retrospective and prospective trials. Violin plots were generated (n=10). (**f**) Monitoring of pulvinar gating strength (**h**_1_^*T*^**h**_2_) before and before cue onset in retrospective trials, which had a reverse logistic shape compared to **Fig. 6f**. Shaded area denotes s.e.m. (n=8). (**g**) Cosine of the angle between two planes in attention trials. The angle of 0 indicates that the two planes were parallel during the post-stimulus period. (**h**) Based on two different choices of the activation function (top; Relu; bottom: tanh), multiplicative gating substantially improved the convergence speed of task learning (p-values, rank-sum test, n=20 for each violin plot).

**Extended Data Fig. 9.**
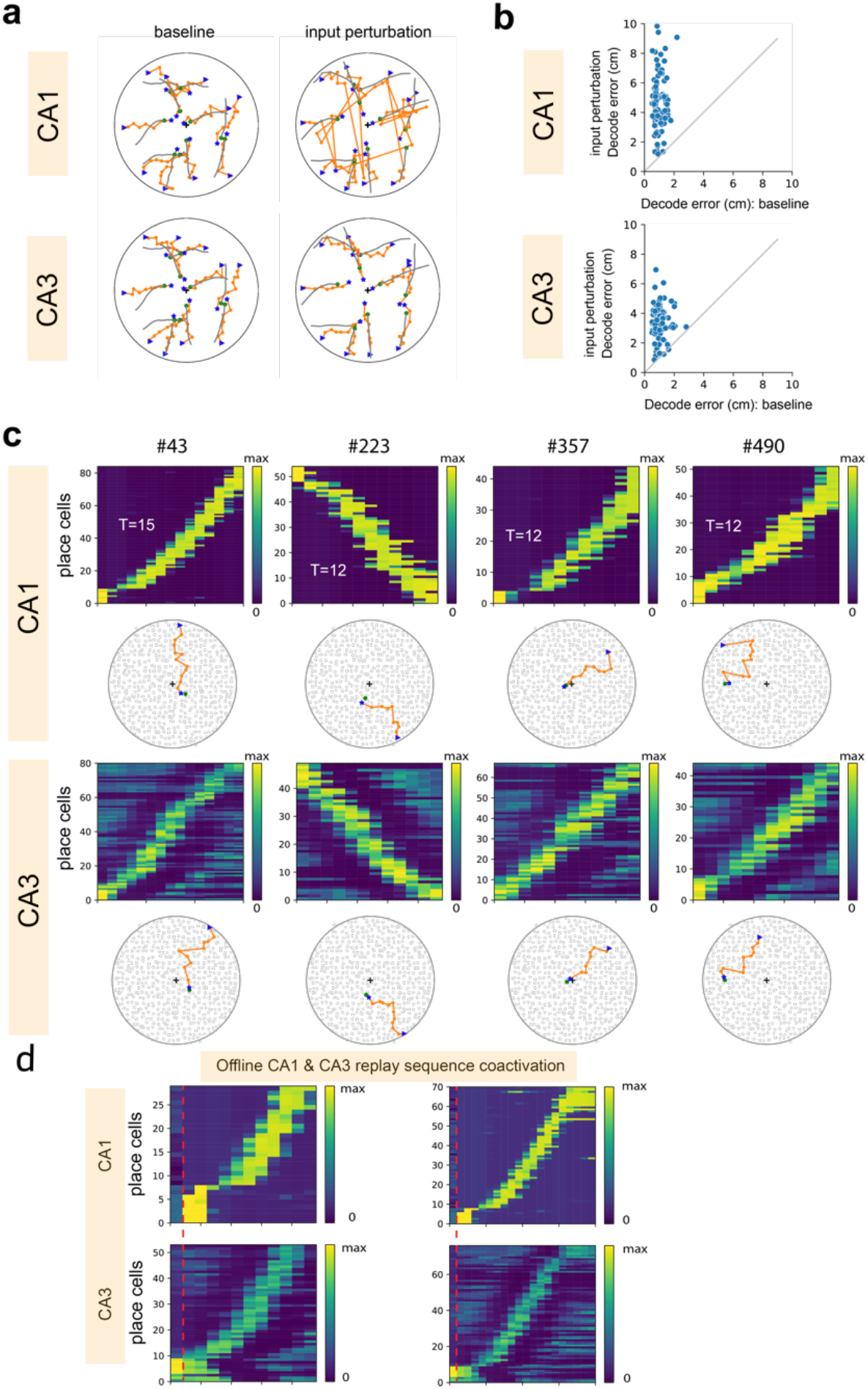
Additional results of the entorhinal-hippocampal network with multiplicative EC-CA1 coupling. **(a)** Representative illustration of CA1 vs CA3 population decoding for trajectory reconstruction without and with input perturbation. Green: true trajectory; Orange: reconstructed trajectory. Blue triangle and star symbols denote the start and the end of trajectories, respectively. (**b**) Scatter plot of mean decoding errors of simulated 2D trajectories (n=100) based on CA1 or CA3 population in baseline and with input perturbation. A bigger change in population decoding error induced by input perturbation suggested that CA1 place cell representations were more plastic or less stable than CA3 place cell representations. (**c**) When CA1 units showed replay sequence, CA3 units also produced neural sequences showing similar or dissimilar spatial trajectories. Note that the initial starting points were identical between CA1 and CA3 replay. (**d**) The timing of CA3 unit reactivation was slightly (i.e., one temporal bin, marked by vertical dashed lines) faster than the onset of CA1 unit reactivation. Two examples are shown.

## Supplementary Material

In this document, we present details of technical discussions, computer simulations, and mathematical derivations related to multiplicative gating mechanism.

### Supplementary Note 1: Functional classification of the mask matrix

We consider a general equation of multiplicative RNN dynamics with dynamic synapses 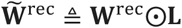 based on a mask matrix **L**

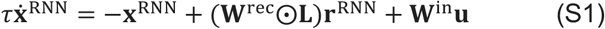

The choice of mask matrix **L** in (S1) modulates the flow of information and determines different functional types:

- *Static binary mask L*_*ij*_ ∈ {0,1}, which can enforce sparse structural connectivity (such as cortical microcircuits) and encourages local or modular dynamics.
- *Random continuous mask L*_*ij*_ ∈ Uniform(0,1) or *L*_*ij*_ ∈ Gaussian(*μ, σ*), which leads to randomized but tunable recurrent strength, can enrich the reservoir of dynamics.
- *Context or task-dependent mask L* = *f*_task_(*context, cue*), enables dynamic reconfiguration of recurrent connectivity, can lead to context-specific attractors, context-gating and flexible cognitive control.
- *Learnable (plastic or modulation) mask L* = *f*_*θ*_, which can be parameterized by *θ* and optimized during training, allows the network to learn what to gate and where (“meta learning”), and allows the network to perform structural generalization or continual learning.
- *Factorized mask* 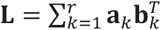 or 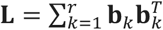, which consists of a low-rank matrix representation (where **a**_*k*_ or **b**_*k*_ denotes the *k*-th learnable basis vector). This will allow the network to learn “what” and “where” to gate information.
- *Structural mask*, which imposes a block-structure in the mask that are specific to subregions or cell types. For instance, 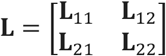 has a two-by-two block structure.
- *Compositional masks*, which implements a series of multiplication operations to impose concurrent multiple gating operations (e.g., **L** = **L**_1_⨀**L**_2_⨀**L**_3_). Individual mask matrices may represent hard-wired connectivity constraints, local Hebbian constraints, or global neuromodulation constraints.

### Supplementary Note 2: Proof of (*A* ⊙ *B*)*v* = *diag*(*AD*_*v*_*B*^*T*^)

#### Notations

Let ***A*** and ***B*** denote two *n*-by-*n* dimensional matrices of the same size; let ***v*** denote an *n*-dimensional vector and let ***D***_***v***_ denote a diagonal matrix with the entries of vector ***v*** along its diagonal.

#### Proof

First, we examine the left-hand-side (LHS) of the equation: (***A*** ⊙ ***B***)***v***.

Let ***C*** = ***A*** ⊙ ***B***, so *C*_*ij*_ = *A*_*ij*_*B*_*ij*_, then it follows that

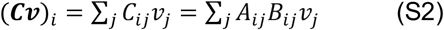

Next, we examine the right-hand-side (RHS) of the equation: *diag*(***AD***_***v***_***B***^*T*^).

We examine the elementwise entry for the matrix ***D***_***v***_***B***^*T*^:

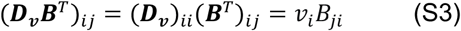

From the rule of matrix multiplication (e.g., ***ABC*** = ***A***(***BC***)), we now compute ***A***(***D***_***v***_***B***^*T*^)

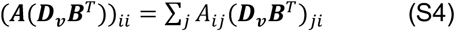

Finally, we substitute (***D***_***v***_***B***^*T*^)_*ji*_ = *v*_*j*_*B*_*ij*_, and rewrite (S4) as follows:

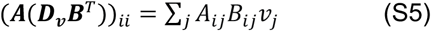

Comparing the LHS and RHS, it completes the proof.

### Supplementary Note 3: Self-gated RNN with time-delay feedback

Equation (9) describes the fast dynamics of auto-associative memory trace **M** computed from the RNN’s time-delay feedback. Employing the mask matrix **M** in multiplicative gating in equation (10) introduces dynamics dynamic synapses in the RNN and enforces adaptive Hebbian associative learning. Note that (**W**^rec^⨀**M**)**r**^RNN^ = *diag*(**W**^rec^**D**_r_**M**^*T*^)= *diag*(**W**^rec^**D**_r_**M**), and the term **D**_r_**M** can be expanded to the following sum: 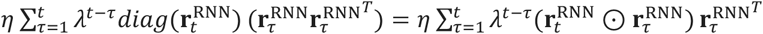.

In contrast, in the additive gating condition

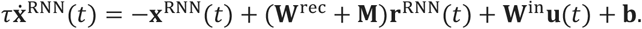

Projecting the firing rate vector **r**^RNN^ onto space **M** yields 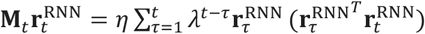, where the inner product 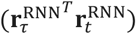 measures the similarity between the current and past RNN activities as an unnormalized attention score^19^ (Ba et al., 2016).

### Supplementary Note 4: Theoretical analysis and empirical studies of 3×3 RNN-FNN models

In the Integration benchmark, we adopted a simple 3-dimensional RNN-FNN network with either multiplicative or additive coupling between the RNN and the FNN. Let’s first rewrite the RNN equations with multiplicative and additive couplings respectively:

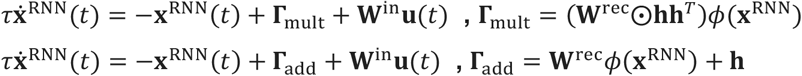

where **Γ**_add_ and **Γ**_mult_ denote the internal update term for the respective additive and multiplicative coupling models. To gain some mathematical insights and make analytic derivations, we assume that the activation function *ϕ* is a logistic sigmoid function with the output range between 0 and 1 and further make the statistical assumptions for model inference: (i) the initial state **x**^RNN^(0)=[0, 0, 0]; (ii) the external input **u** is 0 (e.g., during WM delay period) or is negligible (i.e., |**Γ**_mult_| ≫ |**W**^in^**u**|); (iii) the initial RNN weights follow a Gaussian distribution with zero mean and small variance: 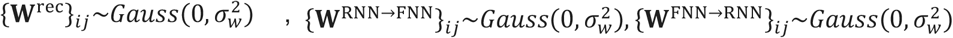, where 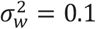.

Let 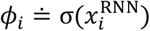, if 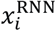 is drawn from a symmetric distribution centered at 0 (including Gaussian), then 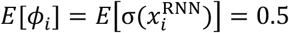, where *E*[·] denotes the mathematical expectation. If 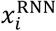 and 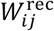 are mutually independent, then 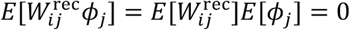. Therefore,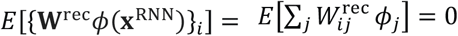. If 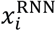 is Gaussian distributed with a zero mean and variance 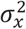, we can derive 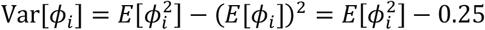. Depending on 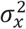, we may derive the lower and upper bounds 0 ≤ Var[*ϕ*_*i*_] ≤ 0.08 (for 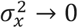 and 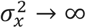, respectively). Similarly, 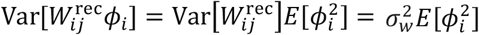 and 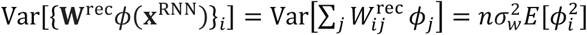, where we obtain the lower and upper bounds: 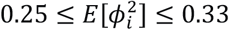.

#### Additive gating

Let *h*_*i*_ denote the *i*-th component of the FNN feedback vector **h**_FB_ = **W**^FNN→RNN^*ϕ*(**W**^RNN→FNN^**r**^RNN^). Let **z** = **W**^RNN→FNN^**r**^RNN^; it is easy to show that *E*[**z**] = **0**. Furthermore, we derive the first and second order moment statistics based on the mutual independence assumption:

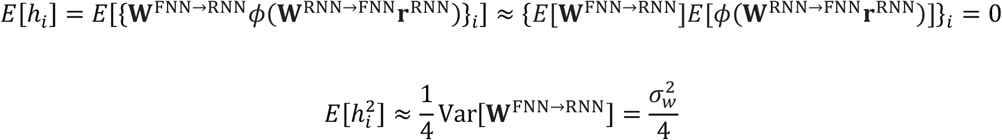

We introduce a new variable: {**Γ**_add_}_*i*_ = {**W**^rec^*ϕ*(**x**^RNN^)}_*i*_ + *h*_*i*_; taking the mathematical expectation yields *E*[{**Γ**_add_}_*i*_] = *E*[{**W**^rec^*ϕ*(**x**^RNN^)}_*i*_] + *E*[*h*_*i*_] = 0. The variance is given by 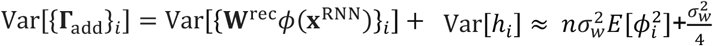. Finally, we have

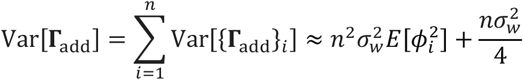

#### Multiplicative gating

To assure the gating term is nonnegative (without changing the algebraic sign of recurrent synaptic connection), we impose a logistic sigmoid function onto the feedback output such that **h**_FB_ = *σ*(**W**^FNN→RNN^*ϕ*(**W**^RNN→FNN^**r**^RNN^)) unless **W**^FNN→RNN^ is purely excitatory. Similarly, based on approximation and numerical stimulation, we derive the following first and second order moment statistics:

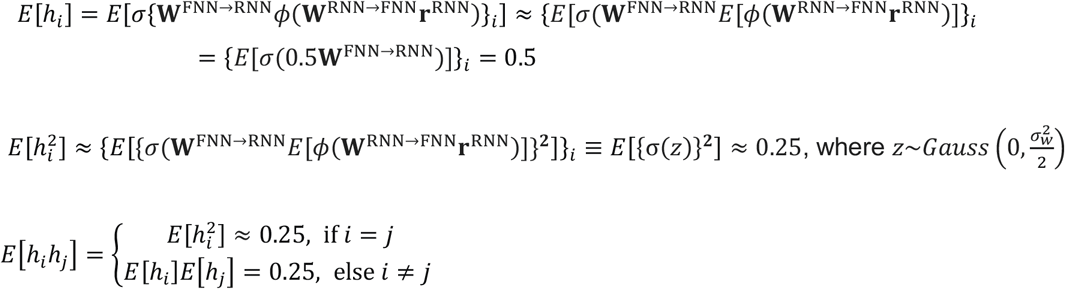

Furthermore, based on of the mutual independence assumption between 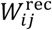 and *h*_*i*_ (and *h*_*j*_), we derive the following first and second moment statistics:

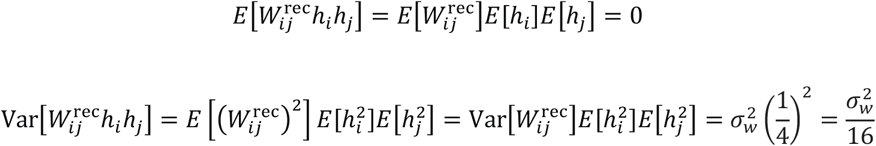

Let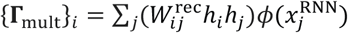; taking the mathematical expectation yields 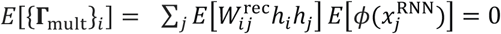. Taking the variance operation yields 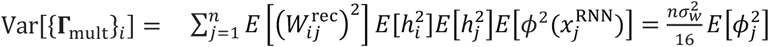. Finally, we have

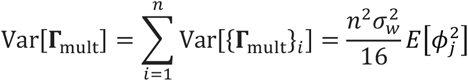

We compare the ratio of the corresponding variance terms between multiplicative and additive gating:

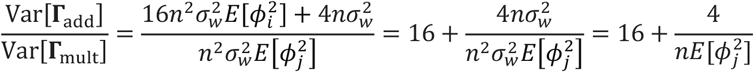

Plugging *n* = 3 into the above equation yields 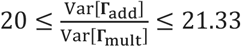 or 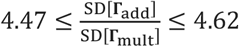.

The above analysis implies that the variance of information flow during incremental latent RNN state update is smaller in multiplicative gating, suggesting a more controlled stable gating mechanism. To validate this theory, we conducted the following experiments. First, we fed an identical random input to the populations of pretrained multiplicative and additive gating RNN-FNN models (*n* = 30 for each class). Each model was initialized with the same initial state condition **x**^RNN^(0) and fed into an identical input sequence **u**(1: 100) of length *T*=100 (25 inputs + 50 noisy delay + 25 inputs). Extracting only the time series during the delay period only, we generated 3-by-50 neural state output **x**^RNN^(1:50) and their corresponding response vector **r**^RNN^(1:50)= *ϕ*(**x**^RNN^(1:50)). We applied dimensionality reduction (PCA followed by t-SNE embedding analysis) on the neural trajectories **r**^RNN^. Our results showed independent of the choice of input sequence, two classes models showed distinct clusters in the two-dimensional embedding space, especially close within-class resemblance among the trained multiplicative coupling models (**Extended Data Fig. 2a**). Next, during the delay version of integration task, we computed the ensembled average of **Γ**_add_ and **Γ**_mult_ for each trained model during the delay period, where was averaged across time and then averaged across all testing sequences. Accordingly, we also computed the respective ensemble average testing accuracy for each model. We further compared the relationship between each model’s accuracy statistic with the update term **Γ** (**Extended Data Fig. 2b**). At the population level, our results showed that (i) the accuracy had a negative Spearman’s rank correlation (*ρ* = −0.455, *p*=0.000259, *n*=60) with the internal update term **Γ**; (ii) multiplicative couplings showed stabilized **Γ**_mult_ and high accuracy mean and low variance, whereas additive couplings showed high variability in both accuracy and **Γ**_add_, with the empirical statistics matching the theoretical derivation results shown in the table below. While these empirical results were calculated based on specific assumptions (*τ* = 2, *dt* = 0.1), the trend remained robust for a different setup close to a discrete-time setting (*τ* = 2, *dt* = 0.4; *ρ* = −0.403, *p* = 0.00142, *n* = 60).

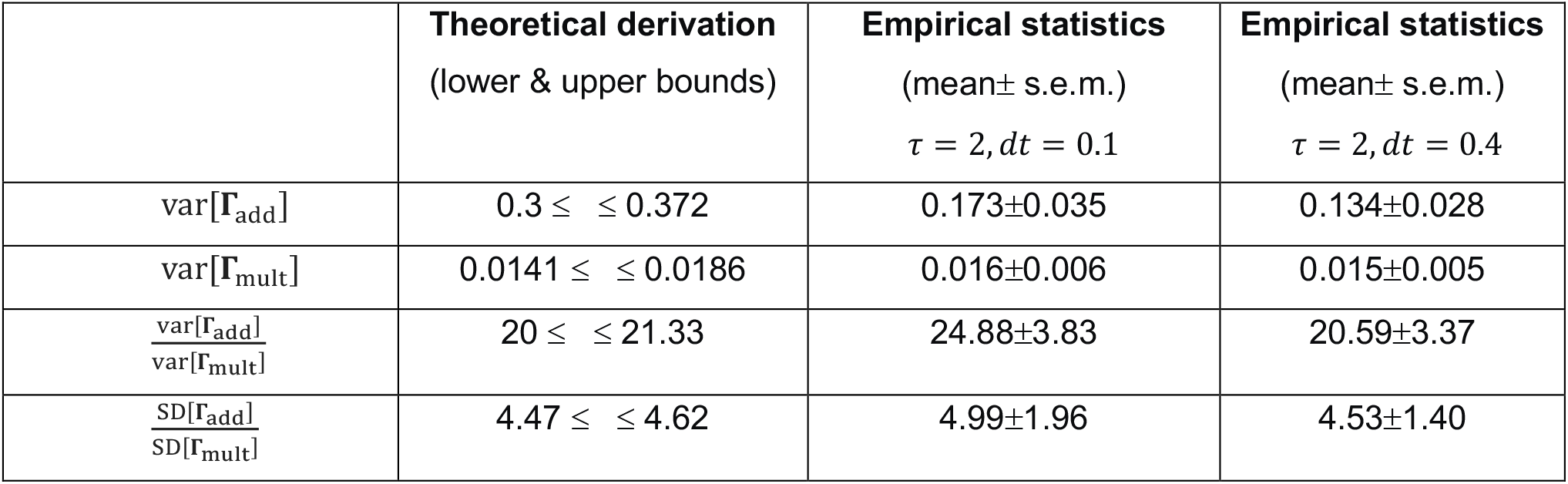

### Supplementary Note 5: 2×2 RNN-FNN computer simulations and phase portrait comparison

To gain geometric insights into context-dependent computation of multiplicative gating, we ran computer simulations with a low-dimensional RNN-FNN system.

In the 2×2 system (**Extended Data Fig. 4**), we manually set **W**^rec^ and generated two vectors that represent feedback activity **h**_FB_ for two different contexts. For simplicity, we assumed that each element of 2-by-1 vector **h**_FB_ was constant and was generated from a standard normal distribution. We randomly initialized the initial state **x**^RNN^(0) and set the external input **u** = **0**. In this case, **L** = **h**_FB_**h**_FB_^*T*^ was a rank-1 matrix. Under three different conditions of **W**^rec^, the RNN-FNN networks produced attractor dynamics such as fixed-points or saddle points that were visualized by the phase diagram. At the fixed point **x**^*^, we had 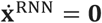 or

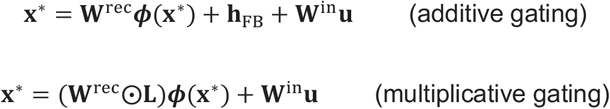

In the case of multiplicative gating, the phase portraits between two contexts were rather different, suggesting that the network could represent rich yet distinct dynamics under two different context inputs. In contrast, in the case of additive gating, the phase portraits between two contexts were qualitatively similar (up to some rotation ambiguity). Our extensive computer simulations showed that multiplicative gating-enabled dynamic synapses may introduce dynamic phase reconfiguration or exhibit phase transition with multiplicative interactions.

### Supplementary Note 6: Jacobian derivations of RNN-FNN models with additive and multiplicative couplings

Let **h**_FB_ = **W**^FNN→RNN^**r**^FNN^ = **W**^FNN→RNN^***ϕ***(**W**^RNN→FNN^**r**^RNN^), **L** = **hh**^*T*^ is a mask matrix. To simplify the mathematical derivation, we assume that ***ϕ*** is a ReLu function and **W**^FNN→RNN^ and **W**^RNN→FNN^ are excitatory (namely, all matrix entries are positive), as in the bidirectional MD-PFC connectivity; therefore, we can simplify the equation **h** = **Cr**^RNN^ and 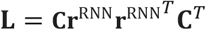, where **C** = **W**^FNN→RNN^**W**^RNN→FNN^.

To show the relationship between RNN connectivity and dynamics, we examine the Jacobian of the RNN. For notation simplicity, let 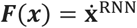; in the additive coupling setting, the Jacobian of the network is computed as

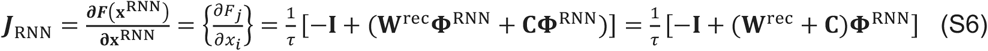

where **Φ**^RNN^ = *diag*{***ϕ***^′^(**x**^RNN^)} represents an *N*_RNN_-by-*N*_RNN_ diagonal matrix containing the derivative of ***ϕ*(x**^RNN^), and **I** denotes an *N*_RNN_-by-*N*_RNN_ identity matrix. Note that if all eigenvalues of ***J***_RNN_ have real parts <0, the RNN will have stable fixed point(s); if some eigenvalues have real parts > 0, then possible oscillations or chaos may exist.

In the multiplicative coupling setting, the mathematical derivation is divided into three steps.

*Step 1: Rewrite* (**W**^rec^⨀**L**)**r**^RNN^ *as a simpler form*

We first write ***y*** ≜ (**W**^rec^⨀**L**)**r**^RNN^ = *diag*(**W**^rec^**D**_***r***_**L**^*T*^) = *diag*(**W**^rec^**D**_***r***_**L**). Because **W**^rec^**D**_***r***_**L** = **W**^rec^**D**_***r***_**hh**^*T*^, taking the diagonal of an *N*_RNN_-by-*N*_RNN_ matrix yields an *N*_RNN_-dimensional vector. In index notation, we have

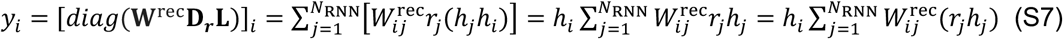

Let ***s*** ≜ **W**^rec^(**r**^RNN^⨀**h**) be a function of **x**^RNN^, then ***y*** = *diag*(**h**)***s*** because *y*_*i*_ = *h*_*i*_*s*_*i*_ and 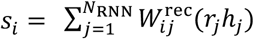. Therefore, we have the following equation:

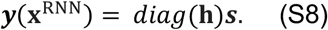

To compute the partial derivative 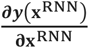, for notation simplicity we use **x** in place of **x**^RNN^ and use *N* in place of *N*_RNN_ in the remaining derivation. Then we write the partial derivative in vector form

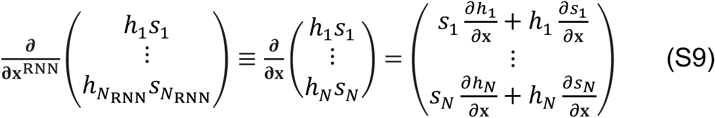

Therefore, by the standard product rule (elementwise product rule *y*_*i*_ = *h*_*i*_*s*_*i*_), we have

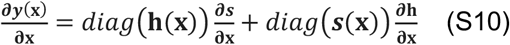

*Step 2: Derivative of* ***s*** = **W**^rec^(**r**^RNN^⨀**h**)

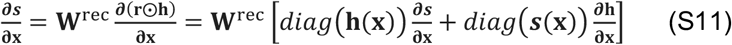

Substituting (S11) into (S10) yields

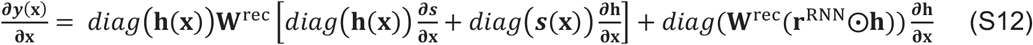

*Step 3: Derivative of* **h** = **W**^FNN→RNN^**W**^RNN→FNN^**r**^RNN^ = **W**^FNN→RNN^**W**^RNN→FNN^***ϕ***(**x**^RNN^) ≜ **C*ϕ***(**x**^RNN^)

Let **Φ**^RNN^ = *diag*(***ϕ***^′^(**x**^RNN^)), we have 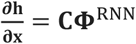.

Putting *Steps 1-3* together, we have

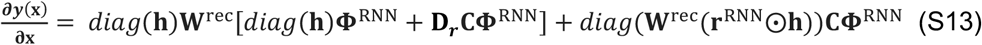

Finally, we compute the Jacobian of RNN-FNN with multiplicative coupling as follows:

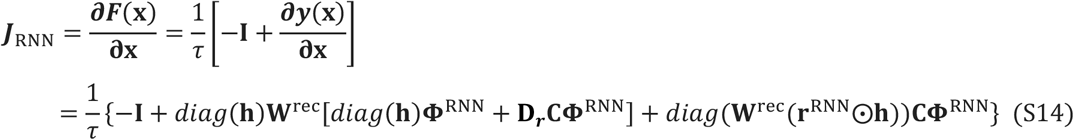

Put together, we summarize the Jacobian derivation in the following table.

| Network | Equation | Jacobian $J_{\text{RNN}}$ |
| --- | --- | --- |
| vanilla RNN | $\tau \dot{\mathbf{x}}^{\text{RNN}} = -\mathbf{x}^{\text{RNN}} + \mathbf{W}^{\text{rec}}\mathbf{r}^{\text{RNN}} + \mathbf{W}^{\text{in}}\mathbf{u}$ | $\frac{1}{\tau} [-\mathbf{I}_{N_{\text{RNN}}} + \mathbf{W}^{\text{rec}}\Phi^{\text{RNN}}]$ |
| RNN-FNN additive coupling | $\tau \dot{\mathbf{x}}^{\text{RNN}} = -\mathbf{x}^{\text{RNN}} + \mathbf{W}^{\text{rec}}\mathbf{r}^{\text{RNN}} + \mathbf{h} + \mathbf{W}^{\text{in}}\mathbf{u}$ | $\frac{1}{\tau} [-\mathbf{I}_{N_{\text{RNN}}} + (\mathbf{W}^{\text{rec}} + \mathbf{C})\Phi^{\text{RNN}}]$, Eq. (S6) |
| RNN-FNN multiplicative coupling | $\tau \dot{\mathbf{x}}^{\text{RNN}} = -\mathbf{x}^{\text{RNN}} + (\mathbf{W}^{\text{rec}} \odot \mathbf{L})\mathbf{r}^{\text{RNN}} + \mathbf{W}^{\text{in}}\mathbf{u}$ | Eq. (S14) |

Finally, without adding additional model complexity, we may consider combining both additive and multiplicative couplings into the RNN dynamics as follows:

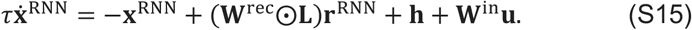

The Jacobian of Eq. (S15) can be readily derived by combining Eq. (S6) and Eq. (S14).

### Supplementary Note 7: Lyapunov exponents of gated RNN-FNN models

Considering Supplementary Notes 4 and 6, we can compute and compare the Lyapunov exponent (LE) of RNN-FNN models based on their Jacobian matrices. In nonlinear dynamical systems, LEs quantify the average rate of divergence or convergence of nearby trajectories in a dynamical system. A positive LE indicates chaotic behavior of the system, where small differences in initial conditions lead to drastically different outcomes over time. Conversely, a negative LE suggests stability of the dynamical system, where trajectories converge. For any dynamical system, the Jacobian matrix captures the local behavior of the system around a specific point in phase space. At each time step, the Jacobian matrix is used to calculate the eigenvalues. The logarithms of these eigenvalues (in modulus), when averaged over time, yield the local LEs. The largest LE is often of particular interest, providing the most significant measure of the system’s sensitivity to initial conditions.

In the 3×3 RNN-FNN system (***F***(**x**): ℝ^3^ → ℝ^3^) for the integration task, we computed the accumulative product of the Jacobian in time: 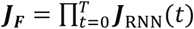. The determinant of Jacobian, or Jacobian determinant, measured at specific point **x**, gives important information about the behavior of a dynamical system near that point. If the Jacobian determinant is positive (or negative), then the system preserves (or reverses) orientation near the point; the absolute value of the Jacobian determinant at that point gives us the factor by which the function expands or shrinks volumes.

In practice, the LE estimate was approximated as the mean rate of separation of latent trajectories in time and then further averaged across *K* trials based on different input sequences with length *T*

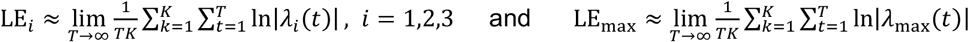

where |*λ*_max_(*t*)| denotes the modulus or absolute value of the maximum eigenvalue of the Jacobian matrix ***J***_RNN_(*t*). If LE_max_ is positive, the stationary dynamics is chaotic and small perturbations will explode, otherwise it is stable, and small perturbations will vanish (Vogt et al., 2022; Vogt et al., 2024).

We further used the integration benchmark and 3×3 RNN-FNN models to make comparisons. We adopted a similar analysis (described in **Supplementary Note 4**) and computed the Jacobian trace (size: 3×3×*T*; *T*=100) of each pretrained RNN-FNN model and applied dimensionality reduction (PCA followed by t-SNE embedding analysis) for visualization. Our results revealed two separated classes between additive and multiplicative gating models (**Extended Data Fig. 2c**). Furthermore, we computed the LE_max_ for each model and correlated it with the accuracy statistics (**Extended Data Fig. 2d**). At the population level, the accuracy had a negative Spearman’s rank correlation (*ρ* = −0.578, *p*=1.3×10^−6^, *n*=60) with the largest Lyapunov exponent. While all estimated LE_max_ values were negative, the multiplicative gating models (mean±SD: −0.0504 ± 0.0006) had lower values than the additive ones (mean±SD: −0.0463 ± 0.0009).

---

**Supplementary Table 1.**
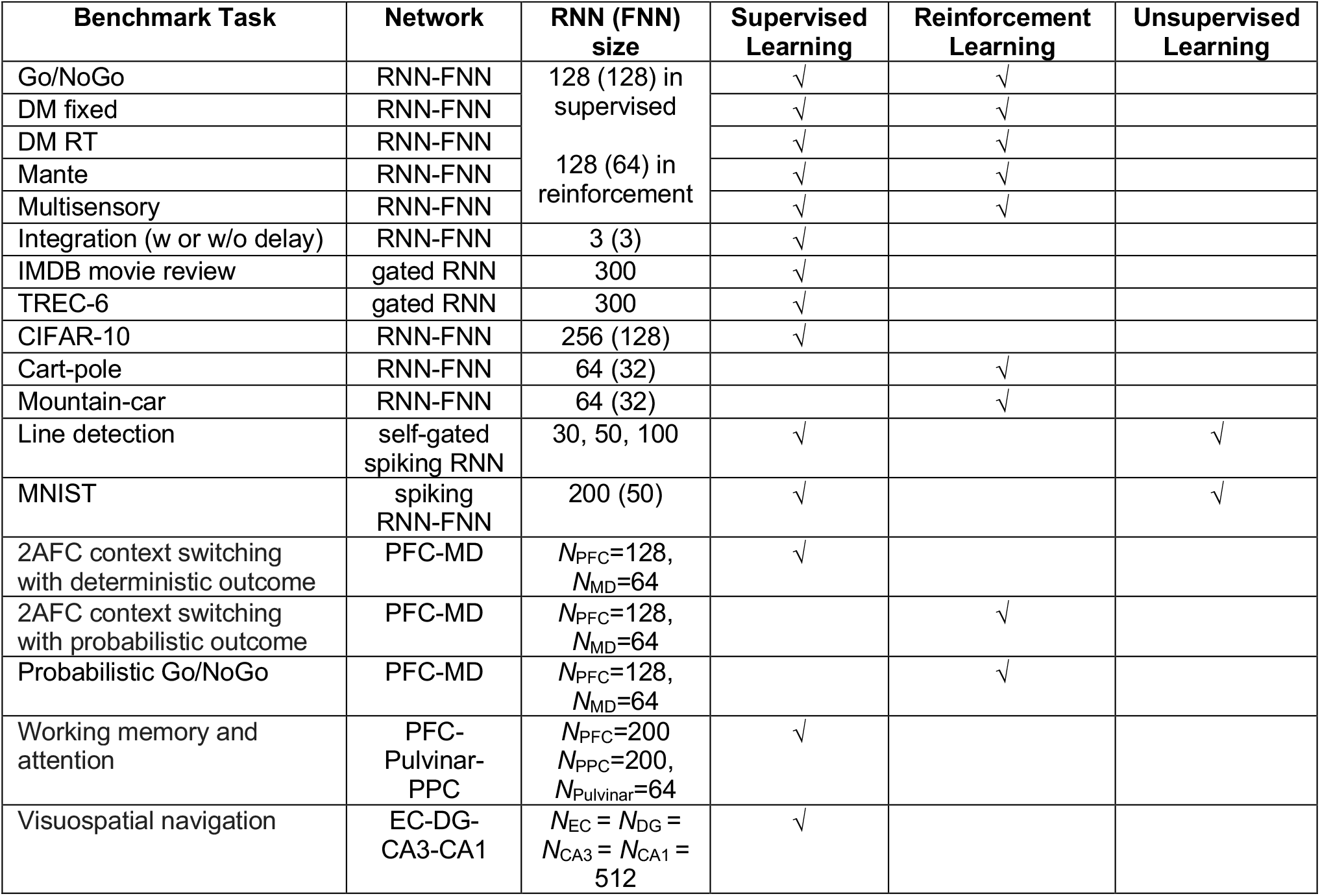
Summary of network configuration for all benchmark experiments.

**Supplementary Table 2.** Default hyperparameter setup in supervised learning benchmark tasks.

| Benchmark task | Time constant $\tau$ & bin size $dt$ | Batch size | Learning rate | Regularization parameter |
| --- | --- | --- | --- | --- |
| Go/NoGo | $\tau = 80 \text{ ms}, dt = 20 \text{ ms}$ | 8 | 0.0005 | 0.1 |
| DM fixed | $\tau = 80 \text{ ms}, dt = 20 \text{ ms}$ | 8 | 0.0005 | 0.1 |
| DM RT | $\tau = 80 \text{ ms}, dt = 20 \text{ ms}$ | 8 | 0.0005 | 0.1 |
| Multisensory | $\tau = 80 \text{ ms}, dt = 20 \text{ ms}$ | 8 | 0.0005 | 0.1 |
| Mante | $\tau = 80 \text{ ms}, dt = 20 \text{ ms}$ | 8 | 0.0005 | 0.1 |
| Integration | $\tau = 2, dt = 0.1$ | 4 | 0.001 | 0 |
| IMDB review | $\tau = 1, dt = 1$ | 256 | 0.0005 | 0 |
| TREC-6 | $\tau = 1, dt = 1$ | 32 | 0.0005 | 0 |
| CIFAR-10 | $\tau = 1, dt = 1$ | 128 | 0.001 | 0 |
| Line detection | $\tau = 1, dt = 1$ | 1, 30, 50, 100 | 0.1 | 0 |
| MNIST | $\tau = 1, dt = 1$ | 256 | 0.1 | 0 |
| 2AFC context switching | $\tau = 80 \text{ ms}, dt = 20 \text{ ms}$ | 16 | 0.0005 | 0 |
| Probabilistic Go/NoGo | $\tau = 80 \text{ ms}, dt = 20 \text{ ms}$ | 16 | 0.0005 | 0 |
| Working memory and attention | $\tau = 80 \text{ ms}$ (PPC),<br>$\tau = 40 \text{ ms}$ (PFC)<br>$dt = 20 \text{ ms}$ | 512 | 0.0005 | 0 |
| Visuospatial navigation | $\tau = 30 \text{ ms}, dt = 20 \text{ ms}$ | 512 | 0.0005 | 0.0001 |

**Supplementary Table 3.** Hyperparameter setup in the SRNN.

| Hyperparameters | MNIST task | MNIST task | MNIST task | Line detection | Line detection |
| --- | --- | --- | --- | --- | --- |
|  | STDP (hidden) | STDP (Inhibitory) | BPTT (Vanilla) | STDP (Vanilla) | BPTT (Vanilla) |
| **Simulation setup** |  |  |  |  |  |
| Time steps per sample ($T$) | 200 | 200 | 20 | 50 | 50 |
| Integration time step ($\Delta t$) | 1.0 ms | 1.0 ms | 1.0 ms | 1.0 ms | 1.0 ms |
| **Neuron parameters** |  |  |  |  |  |
| Membrane time constant ($\tau_m$) | 100.5 ms | 100.5 ms | 20 ms | 20 ms | 20 ms |
| Synaptic time constant ($\tau_{syn}$) | * | * | 10 ms | 10 ms | 10 ms |
| Rest potential ($V_{rest}$) | -65 mV | -60 mV | 0.0 mV | 0.0 mV | 0.0 mV |
| Threshold potential ($V_{thr}$) | -52 mV | -40 mV | 0.5 mV | 0.5 mV | 0.5 mV |
| Reset potential ($V_{reset}$) | -60 mV | -45 mV | 0.0 mV | 0.0 mV | 0.0 mV |
| Membrane resistance ($R_m$) | 100.5 | 100.5 | 1.0 | 1.0 | 1.0 |
| Adaptive threshold time constant ($\tau_\theta$) | $1 \times 10^7$ ms | fixed threshold | fixed threshold | fixed threshold | fixed threshold |
| Threshold increment ($\theta_+$) | 0.05 mV | | | | |
| Refractory period | 5 ms | 5 ms | none | none | none |
| **STDP parameters** |  |  |  |  |  |
| Trace time constants ($\tau_{pre}, \tau_{post}$) | 20 ms | none | N/A | 2000 ms | N/A |
| Learning rate $\eta_+$ (potentiation rate) | 0.01 | none | N/A | 0.01 | N/A |
| Learning rate $\eta_-$ (depression rate) | 0.0001 | none | N/A | 0.0001 | N/A |
| $a_{pre}$ (trace amplitude factors) | 0.1 | none | N/A | 0.001 | N/A |
| $a_{post}$ (trace amplitude factors) | 0.05 | none | N/A | 0.001 | N/A |

\* **Footnote:** For the MNIST problem, we modified the SDTP rule and used the instant update for the current equation of the LIP neuron $I_j^{RNN}(t) = \sum_i W_{ij}^{in} s_i^{in}(t) + \sum_k W_{jk}^{rec} s_k^{hid}(t - 1)$ .

